# Processing strategies for improving cortical thickness correspondence between low-field and high-field MRI in young people

**DOI:** 10.64898/2026.07.08.737238

**Authors:** Sunah Choi, Julia Shaw, Rebecca Cooper, Mary Corcoran, Soumya Sathe, Rebecca Hayes, Isabelle Elder, Alfredo Lucas, Chetan Vadali, Joel M. Stein, Maria Jalbrzikowski

**Author notes:** **Corresponding author** Maria Jalbrzikowski, PhD, Department of Psychiatry and Behavioral Sciences, Boston Children’s Hospital, 1 Brookline Pl, 02445, Brookline, MA, United States.

## Abstract

Portable low-field MRI systems are a promising complement to conventional high-field systems, enabling broader access to MRI. However, correspondence in cortical thickness estimates between low- and high-field MRI in young people remains limited despite its importance for neurodevelopment and psychopathology. To evaluate how multiple low-field image processing approaches improve cortical thickness correspondence with high-field MRI in a large sample of young individuals, we collected T1-weighted (T1w) and T2-weighted (T2w) data using both low-field (64mT) and high-field (3T) MRI within one week of each other from a community sample of young people. We applied deep learning–based super-resolution methods (SynthSR v1 and SynthSR v2) followed by recon-all, or other reconstruction methods (recon-all-clinical and recon-any), to low-field data acquired across multiple sequences (T1w and T2w) and orientations (axial, coronal, sagittal, and multi-orientation average), with and without resampling and/or co-registration. We assessed global, lobar, and regional cortical thickness correspondence with 3T MRI measures based on the Desikan–Killiany atlas using Pearson and intraclass correlations. We compared pipelines using Steiger’s Z-tests and Fisher’s Z-tests. A total of 150 individuals (mean age, 18.63±5.07; 80 female) were included. We observed the highest global correspondence with recon-all-clinical applied to coronal T1w images (*r*=0.40, *p*_FDR_=3.02e-05), which significantly exceeded the best global correspondence in our previous study (Fisher’s *Z*=1.99, *p*=0.047). At the lobar and regional levels, multi-orientation T2w images processed with recon-all-clinical showed the highest correspondence across the greatest number of regions (4/12 lobes; 13/68 regions). This pipeline showed the highest correspondence and largest improvements relative to recon-all alone in frontal, cingulate, and temporal regions, including the right pars triangularis (*r*=0.52, *p*_FDR_=4.78e-11; Steiger’s *Z*=4.78, *p*_FDR_=4.25e-06), right caudal anterior cingulate (*r*=0.47, *p*_FDR_=3.83e-09; Steiger’s *Z*=5.46, *p*_FDR_=1.32e-07), and left parahippocampal regions (*r*=0.58, *p*_FDR_=2.98e-14; Steiger’s *Z*=5.17, *p*_FDR_=6.01e-07). We observed significantly improved cortical thickness correspondence in low-field MRI in young people relative to recon-all alone and our prior work. The *recon-all-clinical* pipeline yielded moderate correspondence, particularly in frontal, cingulate, and temporal regions. Our results demonstrate a methodological improvement in the use of low-field MRI for assessing cortical thickness in young people, providing a quantitative benchmark for current tools in this area.

## 1 Introduction

MRI is an essential tool used in pediatric research and clinical care (Figueiro Longo et al., 2022; Lee-Jayaram et al., 2020), offering insights into the relationships between brain maturation and cognition, behavior, and psychopathology (Andreou & Borgwardt, 2020; Ellis et al., 2020; Lerch et al., 2017). However, conventional high-field (HF)-MRI systems are expensive (at least 1 million USD per Tesla; Sarracanie et al., 2015), require specialized infrastructure and expertise, and are largely immobile, limiting their accessibility (Arnold et al., 2023). As a result, opportunities to conduct neuroimaging research within diverse populations remain limited, hindering the translation of promising findings into broader clinical and community settings. Low-field (LF)-MRI, a promising complement to conventional HF-MRI systems, offers advantages in cost (∼$20,000-400,000; Hyperfine representative, personal communication, October 24, 2024; Sarracanie et al., 2015; Zhao et al., 2024), portability, and accessibility. Although definitions of LF-MRI vary across the literature, field strengths below 0.3 T have been broadly described as LF (Arnold et al., 2023). Portable LF-MRI scanners in this range consume less power, require less operative expertise, occupy less physical space, and do not require cryogenic cooling (Arnold et al., 2023; Kimberly et al., 2023). These features, as well as designs focused on ease of use, enable their use in diverse settings (e.g., intensive care units, offices, and/or low-resource environments) (Abate et al., 2024). The lower acoustic noise and open designs of LF-MRI scanners may also improve tolerability and reduce the need for sedation in pediatric populations (Deoni et al., 2021; Rupprecht et al., 2000).

Previous studies have reported strong correspondence between LF- and HF-MRI in surface area and volumetric measures in adults (Iglesias et al., 2022; Lucas et al., 2025; Sorby-Adams et al., 2024; Váša et al., 2025) and youth (Baljer, Zhang, et al., 2025; Cooper et al., 2024; Deoni et al., 2021), particularly when super-resolution methods are used. Super-resolution refers to techniques that convert low-resolution images into high-resolution images (Ayde et al., 2025), including multi-orientation image averaging, which combines multiple low-resolution images into one high-resolution image (Arnold et al., 2022; Askin Incebacak et al., 2022; Sui et al., 2021). They also include image synthesis techniques, which use deep learning models such as convolutional neural networks (CNNs) (Iglesias et al., 2022; Lucas et al., 2025). CNNs learn a transformation between input and target images by automatically extracting hierarchical visual features. However, demonstrating strong cortical thickness correspondence between LF- and HF-MRI measures remains challenging in young people (Cooper et al., 2024). These findings highlight both the promise and current limitations of LF-MRI for research and clinical applications.

Cortical thickness is a key marker of both neurodevelopment and psychopathology (Frangou et al., 2022; Hettwer et al., 2022). Across the first two to three decades of life, cortical thickness undergoes substantial age-related changes, with age explaining up to 59% of its variance (Frangou et al., 2022). In addition, lower cortical thickness is a well-established finding across multiple psychiatric disorders (Hettwer et al., 2022; Rimol et al., 2012; Van Erp et al., 2018). Therefore, accurate estimation of cortical thickness is critical for characterizing brain development and identifying alterations associated with psychopathology.

Recently, several analytic approaches have been introduced to improve LF-HF MRI correspondence (Ayde et al., 2025). SynthSR (Iglesias et al., 2022) is a CNN-based super-resolution method that synthesizes high-resolution T1-weighted (T1w) images from LF-MRI scans. Compared with the original version (SynthSR v1), SynthSR v2 uses domain-randomized training to accommodate more diverse inputs, incorporates an auxiliary segmentation task, and inpaints abnormalities with normal-appearing tissue. Adult studies have reported similar or modestly improved performance with SynthSR v2 based on limited volumetric comparisons (Iglesias et al., 2023; Sorby-Adams et al., 2024), but did not directly compare correlation strengths between versions to assess how well individual variation was preserved. Moreover, to our knowledge, no study has directly compared global, lobar, and regional cortical thickness estimates between SynthSR versions in young people. Recon-all-clinical (Gopinath et al., 2023; Gopinath, Greve, et al., 2025) and recon-any (Gopinath, Sorby-Adams, et al., 2025) are neuroimaging processing pipelines that implement brain segmentation and cortical surface reconstruction—standard components of conventional pipelines—using CNN-based approaches. Unlike SynthSR, the recon-all-clinical and recon-any pipelines do not generate super-resolution images but can operate directly on raw images. While both pipelines can accommodate diverse inputs, recon-all-clinical is designed for heterogeneous clinical HF-MRI, whereas recon-any was specifically trained to account for LF-MRI characteristics, including lower signal-to-noise ratio (SNR), reduced spatial resolution, and altered tissue contrast.

These approaches have improved correspondence between LF- and HF-MRI cortical thickness estimates. One study in adults processed LF scans using the recon-any pipeline and observed strong cortical thickness correspondence with HF scans, achieving Pearson correlations of *r*=0.70 with T2-weighted (T2w) scans (Gopinath, Sorby-Adams, et al., 2025). The superior CSF-gray matter contrast of LF T2w scans may enable more accurate delineation of the pial surface (Gopinath, Sorby-Adams, et al., 2025); however, the quality of CSF-gray matter contrast could vary across LF-MRI scanner types. Most studies comparing LF-HF cortical thickness correspondence have focused on adults. Our initial work in young people found that applying multi-orientation image averaging and SynthSR improved the correspondence for brain structure surface area and volume, but not cortical thickness (Cooper et al., 2024). Another study tested multiple processing pipelines in a youth sample (N=12, ages 10–12 years), including super-resolution and FreeSurfer-based methods on T2w images, but cortical thickness correspondence remained low (*r*≤0.3) (Pretzsch et al., 2025).

Here, we tested how diverse MRI processing approaches improved LF-HF MRI cortical thickness correspondence in a large community sample of young people. First, we examined the degree to which our previous cortical thickness findings hold in a larger sample, which expanded over twofold (N=150 vs. 70). Next, we applied either deep learning-based super-resolution methods (SynthSR v1 and SynthSR v2) followed by recon-all or deep learning-based reconstruction pipelines (recon-all-clinical and recon-any) to multiple LF-MRI sequences and orientations and assessed their correspondence with 3T MRI. We also examined whether preprocessing approaches, including co-registration of LF-MRI scans and isotropic resampling, further improve the correspondence. Finally, we visually assessed pipeline output quality and examined whether restricting analyses to good-quality outputs affected cortical thickness correspondence.

## 2 Methods

### 2.1 Participants

We collected data from 152 young individuals (9-26 years, N<18 years=57) from the Boston metro area between July 2022 and November 2025, including 70 participants from our previous study (Cooper et al., 2024). Consistent with our previous study (Cooper et al., 2024), exclusion criteria included a history of brain infection, the presence of a neurological or neurodegenerative disorder, a neurodevelopmental disorder that might interfere with study participation, a major psychiatric disorder other than Attention Deficit-Hyperactivity Disorder or a past episode of Depression, and/or any MRI contraindications. We obtained written informed consent from all participants; participants under 18 provided written assent and parental consent. All procedures were approved by the Institutional Review Board of Boston Children’s Hospital (IRB No. P00041773).

### 2.2 MRI acquisition

We acquired T1w and T2w brain MRI scans for each participant using a Siemens Magnetom Prisma 3T scanner at Boston Children’s Hospital (T1w: scan duration 6:54 min, resolution 0.8 mm isotropic; T2w: scan duration 5:57 min, resolution 0.8 mm isotropic). We used a Hyperfine Swoop 64 mT scanner (NeuroPod Mk 1.8; software v8.4.0) to acquire LF brain scans. We acquired T1w gray/white and T2w fast sequences in axial, coronal, and sagittal orientations, with two acquisitions per sequence and orientation (2 sequences x 3 orientations x 2 acquisition=12 total scans, total scan duration 52:25 min, resolution 1.6×1.6×5 mm) (Figure 1). We chose the T2w fast sequences based on results from internal pilot testing, as we found that the standard and fast T2w returned scans of similar visual quality. We chose to acquire T1w gray/white sequences because these scans enhance the gray/white matter boundary contrast. 3T and LF-MRI scans were acquired within one week. Scan parameters are provided in Table S1. A trained researcher reviewed LF and 3T scans during acquisition and repeated sequences when artifacts or other quality issues were detected. After image processing, output volumes were visually assessed for artifacts by two independent raters (S.C. and J.S.; see Supplementary Methods and Figure S1 for details).

**Figure 1.**
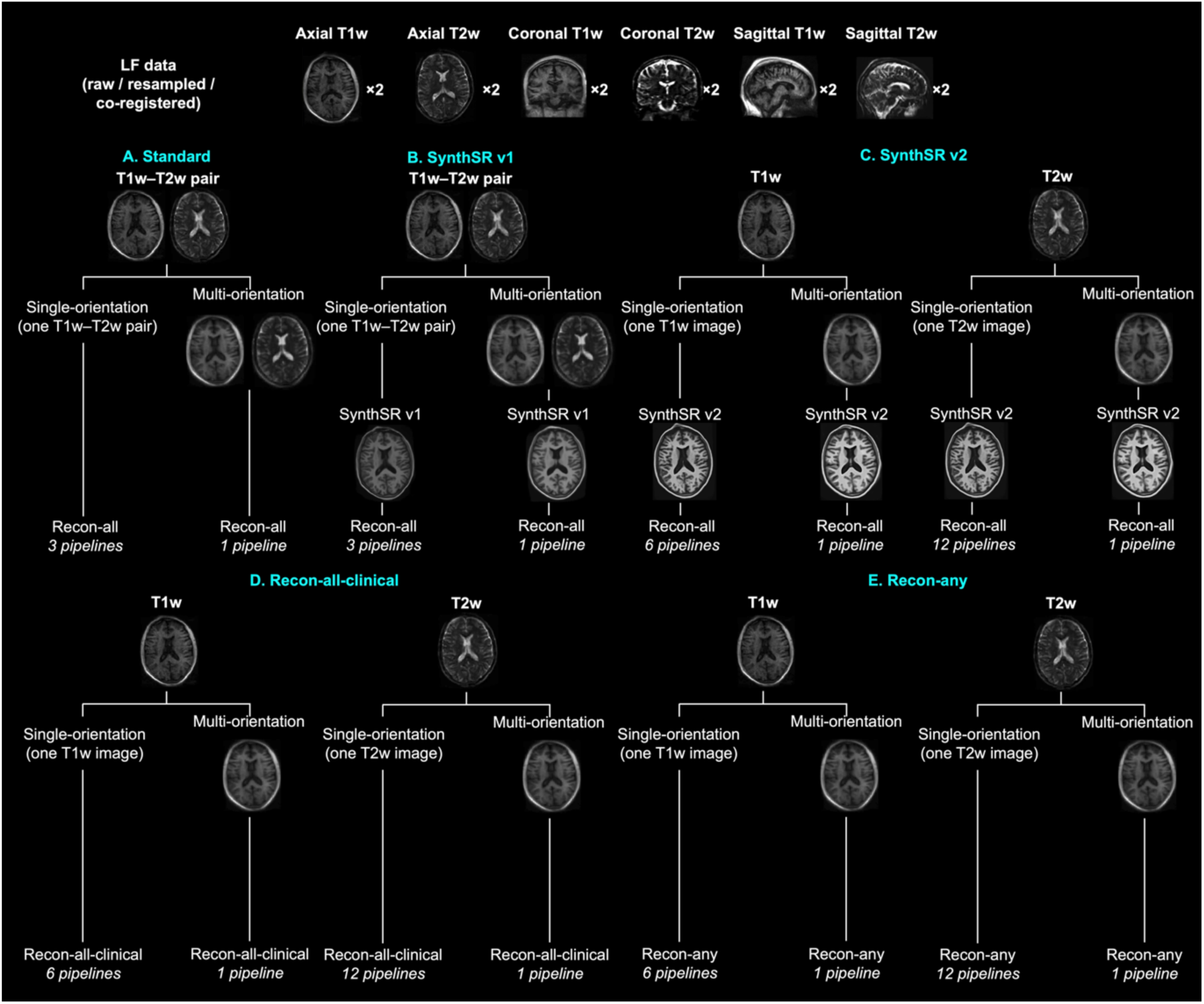
Flowchart of the low-field (LF) image processing. The top row shows the types of LF images used for image processing. Single-orientation images refer to T1w or T2w images acquired in one direction (i.e, axial, coronal, or sagittal). We also generated multi-orientation images by averaging all available images for the T1w sequences and also averaging all available images for the T2w sequences. A T1w–T2w pair refers to T1w and T2w images from the same orientation used together as inputs. Images were input into pipelines (A)–(E) either with or without resampling and/or co-registration. Pipelines were defined by processing method and input type: (A) standard recon-all with a T1w–T2w pair; (B) SynthSR v1–based processing with a T1w– T2w pair; (C) SynthSR v2–based processing with T1w or T2w images; (D) recon-all-clinical with T1w or T2w images; and (E) recon-any with T1w or T2w images. The number beneath each branch indicates the number of processing pipelines represented, with 68 pipelines in total (Table S3). Representative axial images are shown to illustrate each pipeline.

### 2.3 MRI processing

We performed resampling and co-registration of LF scans using the ANTsPy Python module (v0.3.9) (Tustison et al., 2021). We resampled T1w and T2w scans to an isotropic voxel size of 1.5 mm. We then co-registered T2w scans to T1w scans acquired in the same orientation using affine registration and applied the transformations to the T2w scans using linear interpolation. For pipelines using both T1w and T2w images, we used resampled and co-registered image pairs from the same orientation. For pipelines using T1w images alone, we applied the pipeline with and without resampling. For pipelines using T2w images alone, we evaluated four preprocessing approaches: co-registration to the corresponding T1w image and resampling, co-registration only, resampling only, and neither. Co-registration to T1w was retained for T2w-only pipelines to maximize consistency with pipelines using both T1w and T2w images. We refer to pipelines that included neither resampling nor co-registration as ‘raw’ pipelines. Using the same ANTsPy module, we generated multi-orientation images by combining multiple low-resolution images into a single isotropic high-resolution image for each sequence, following our previous method (Cooper et al., 2024). This procedure combines complementary spatial information across orientations and can reduce random image noise. Specifically, we selected the first acquired resampled axial image from each sequence (T1w and T2w) as the sequence-specific reference. We first aligned the T2w reference image to the T1w reference image using affine registration. We then used affine registration to align each of the remaining resampled T1w and T2w images to their corresponding sequence-specific reference. We applied all transformations using linear interpolation. Finally, we separately averaged the registered T1w and T2w images voxel-wise across orientations and repeated acquisitions to generate one multi-orientation image for each sequence. We refer to these images as multi T1w and multi T2w images, respectively.

We performed subsequent LF image processing using tools implemented in FreeSurfer (Figure 1). First, we estimated cortical thickness from T1w and T2w images without additional image synthesis or alternative processing pipelines using the recon-all command in FreeSurfer v7.3.2. We refer to this pipeline as the “standard” pipeline. Recon-all performs cortical reconstruction and structural analysis and can use T2w images as auxiliary inputs alongside T1w images to improve reconstruction quality (Fischl et al., 2002). We also applied SynthSR v1 (Iglesias et al., 2022), SynthSR v2 (Iglesias et al., 2023), recon-all-clinical (Gopinath, Greve, et al., 2025), or recon-any (Gopinath, Sorby-Adams, et al., 2025) to further process LF images. The project began in January 2024, shortly after we learned of the availability of SynthSR v2, following the publication of our first manuscript on LF-MRI, which focused on improvements in cortical reconstruction using SynthSR v1 (Cooper et al., 2024). Further details on the rationale for tool selection are reported in the Supplementary Text. We applied SynthSR v1 with the Hyperfine option, which requires both T1w and T2w images as inputs, and analyzed the synthesized output volume (a single T1w image) using recon-all in FreeSurfer v7.3.2. We processed T1w and T2w images separately using SynthSR v2 and analyzed each resulting output volume (a T1w image) using recon-all in FreeSurfer v7.4.1. We applied recon-all-clinical separately to T1w and T2w images in FreeSurfer v7.4.1 to perform image processing and structural analysis. Similarly, we applied recon-any separately to T1w and T2w images in the FreeSurfer development version (downloaded 8/25/2025). Because some tools are only available in newer FreeSurfer versions and recon-all yields identical results across 7.X releases, different versions were used across pipelines. Descriptions of each approach are detailed in Table S2. Table S3 lists the 68 pipelines examined.

We used 3T scans as the reference standard to evaluate correspondence with LF images. We processed T1w and T2w 3T scans using recon-all in FreeSurfer v7.3.2. As mentioned above, recon-all can use T2w images as auxiliary inputs to improve reconstruction quality; therefore, estimates derived from both T1w and T2w 3T images served as the primary reference, and we confirmed their high correspondence with T1w-only 3T estimates (see Supplementary Results). We examined global, lobar, and regional estimates of cortical thickness. We obtained 68 cortical region estimates from the Desikan-Killiany atlas. The global measure was defined as the average of the mean cortical thickness values from the left and right hemispheres. Lobe-level measures were derived using the FreeSurfer --lobesStrict annotation. Analyses were restricted to pipelines that produced valid outputs for all participants, excluding 3 pipelines that failed for a subset of participants (Figure 2A-B).

**Figure 2.**
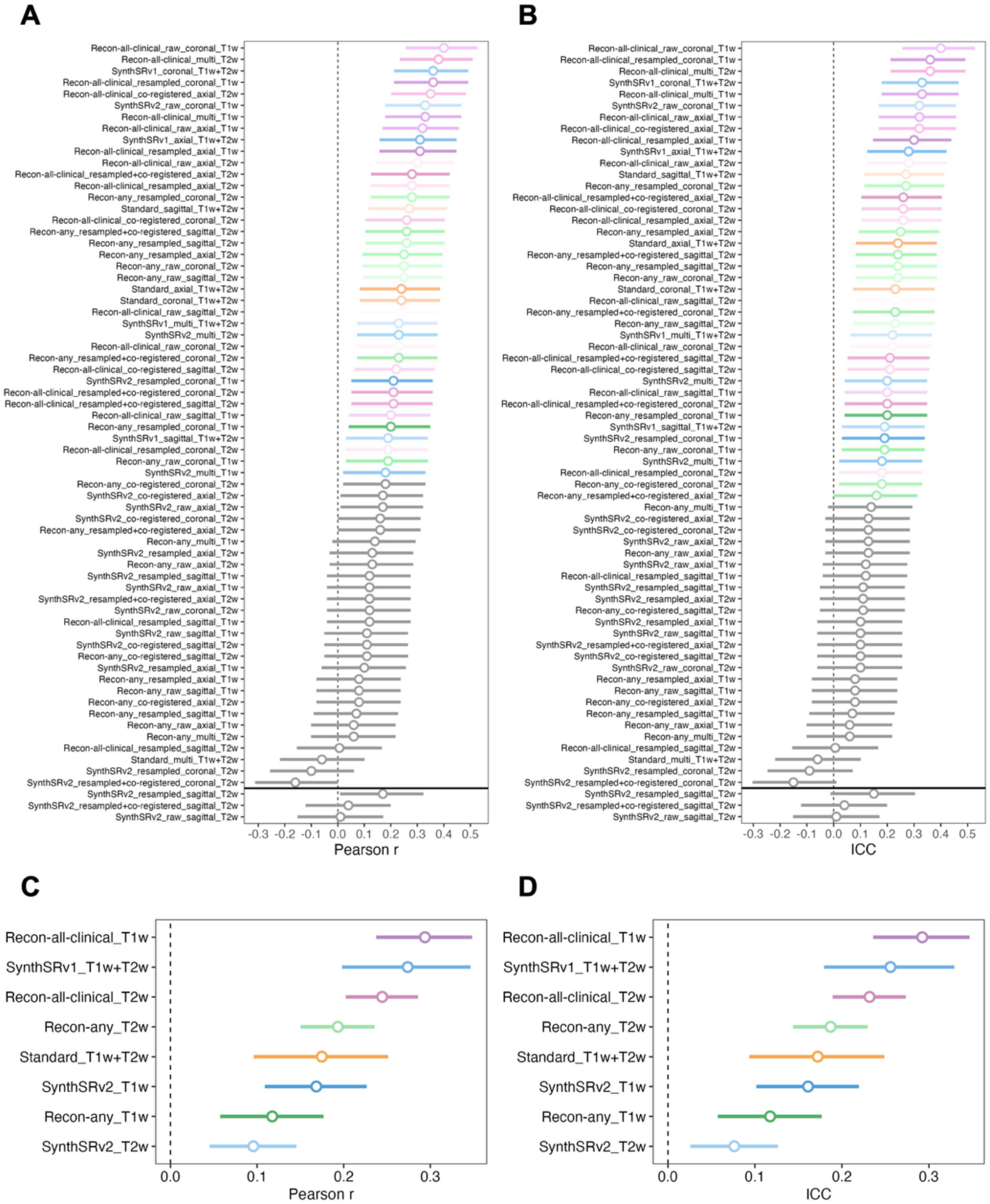
Global correspondence by pipeline and pipeline group. Pearson correlation coefficients (*r*; A) and intraclass correlation coefficients (ICCs; B) for each pipeline. Pipelines above the horizontal divider were successfully completed for all participants, whereas those below the divider failed for a subset of participants. Pearson *r* (C) and ICC (D) values averaged across pipelines within each pipeline group. Pipelines and pipeline groups are ordered by descending global correspondence. Circles indicate correspondence coefficients, whiskers denote 95% confidence intervals, and grey indicates pipelines with non-significant correspondence.

### 2.4 Statistical analysis

We conducted all analyses in R (v4.3.2). Correspondence between LF and 3T scans was examined using Pearson correlation coefficients and intraclass correlation coefficients (ICCs). Pearson correlations assessed the linear association between LF and 3T measurements and the preservation of participants’ relative rankings, whereas ICCs assessed the consistency of measurements across field strengths relative to between-participant variability. For single-orientation pipelines, we used the first good quality scan acquired to calculate correspondence. We assessed global correspondence across all pipelines and restricted lobar and regional analyses to pipelines with valid outputs for all participants and significant global correspondence (*p*_FDR_<0.05). We also examined global correspondence using only volumes rated as “good” based on the quality assessment. To summarize pipeline correspondence, we grouped individual pipelines based on the processing method and input sequence (e.g., all values came from recon-all-clinical processing on T1w images) and averaged global correspondence across pipelines within each group (i.e., across orientations and preprocessing methods), with Fisher’s Z-transformation applied prior to averaging to approximate a normal distribution. We re-ran analyses in a subset of participants under 18 years of age (N=57) to confirm that similar LF–3T correspondence that was observed in the full age sample was also observed in youth.

To compare correspondence across processing approaches, we used Steiger’s Z-tests for individual pipelines and Fisher’s Z-tests for pipeline groups and comparisons with a previous study. These tests assessed whether a given processing approach statistically improved correspondence relative to a comparison approach. For regional measures, we separately identified the pipeline with the highest Pearson correlation coefficient and the pipeline with the highest ICC among all pipelines in each region. We then counted the number of regions in which each pipeline achieved the highest Pearson correlation coefficient or ICC. We used a chi-square test to determine if specific pipelines had a greater frequency of top-performing regions. To determine which pipelines performed better than others, we examined standardized residuals from the chi-square test, with positive residuals indicating more dominant regions than expected and negative residuals indicating fewer. Standardized residuals with |*z*| > 1.96 were considered statistically significant (*p* < 0.05). To satisfy chi-square test frequency assumptions, we excluded pipelines with only one region achieving the highest coefficient.

We performed multiple testing corrections using the false discovery rate (FDR; *p*_FDR_<0.05) within each relevant family of tests. Detailed correction procedures are provided in the Supplementary Methods.

### 2.5 Secondary analyses

Based on the primary statistical analyses, we identified the three best-performing LF processing pipelines and then conducted secondary analyses, further investigating scan quality and agreement with 3T measurements. As a measure of scan quality, we extracted the Euler number from all 3T images, as well as images for the best-performing LF processing pipelines. The Euler number reflects the topological complexity of the reconstructed cortical surface, and has been used as an index of data quality (Dale et al., 1999). A more negative Euler number indicates greater topological complexity and a greater number of topological defects, whereas numbers closer to 2 (the maximum) indicate a smooth cortical reconstruction. Euler numbers less than -217 indicate poor image quality (Rosen et al., 2018). Consistent with previous work (Rosen et al., 2018), we averaged Euler numbers across both hemispheres.

Within the three best performing LF processing pipelines, we also used Bland-Altman analyses to assess levels of agreement between global, lobar, and regional estimates from LF processing pipelines and 3T values. Bland-Altman analyses evaluate overall levels of agreement and identify systematic biases between measurement methods.

## 3 Results

### 3.1 Demographics

After excluding two participants due to motion artifacts in the MRI data, the final sample included 150 young people. Table 1 reports sample demographic characteristics.

**Table 1.** Demographic features of the participants.

|  | Female | Male | Total |
| --- | --- | --- | --- |
| Total N | 80 | 70 | 150 |
| Mean Age (SD) | 18.41 (5.00) | 18.87 (5.17) | 18.63 (5.07) |
| Age Range | 9–26 | 9–26 | 9–26 |
| Race N (%) |  |  |  |
| American Indian/<br>Alaskan Native | 2 | 1 | 3 (2.0%) |
| Asian | 19 | 20 | 39 (26.0%) |
| Black | 4 | 2 | 6 (4.0%) |
| White | 43 | 41 | 84 (56.0%) |
| Native Hawaiian/<br>Pacific Islander | 0 | 0 | 0 (0.0%) |
| Interracial | 12 | 6 | 18 (12.0%) |
| Ethnicity N (%) |  |  |  |
| Hispanic/Latino | 9 | 8 | 17 (11.3%) |
| Not<br>Hispanic/Latino | 71 | 62 | 133 (88.7%) |

### 3.2 LF quality assessment

Most pipelines produced good-quality output volumes (Table S4), with the pipeline using SynthSR v2 on resampled and co-registered sagittal T2w images having the highest proportion of poor-quality volumes (8.7%). Poor-quality volumes typically exhibited blurring or signal loss artifacts, often in regions near the cerebellum (Figure S2).

### 3.3 Global correspondence

For Pearson correlation analyses, 38 of 68 pipelines showed statistically significant LF-3T cortical thickness correspondence (Table 2, Table S5). Recon-all-clinical with raw coronal T1w (*r*=0.40, *p*_FDR_=3.02e-05) and recon-all-clinical with multi T2w (*r*=0.38, *p*_FDR_=4.94e-05) showed the numerically highest coefficients (Figure 2A, Figure 3). ICC results were similar (Figure 2B, Tables S6-S7). Corresponding specification plots summarizing the characteristics of the pipelines are provided in Figures S3–S4. Three pipelines had incomplete results because FreeSurfer processing failed for some participants, as detailed in the Supplementary Results. Correspondence estimates were similar in participants under 18 years of age, with the recon-all-clinical raw coronal T1w pipeline showing *r*=0.41 (*p*_FDR_=0.012) and recon-all-clinical multi T2w pipeline showing *r*=0.47 (*p*_FDR_=0.006). At the pipeline average level, the recon-all-clinical T1w group showed the highest mean correspondence with 3T (*r*=0.29; ICC=0.29; Figure 2C-D). Fisher’s Z-tests indicated that this group showed significantly higher correspondence with 3T than the standard T1w+T2w group (*Z*=2.46, *p*_FDR_=0.026), SynthSR v2 T1w group (*Z*=3.04, *p*_FDR_=0.006), SynthSR v2 T2w group (*Z*=5.13, *p*_FDR_=3.73e-06), recon-any T1w group (*Z*=4.22, *p*_FDR_=1.04e-04), and recon-any T2w group (*Z*=2.79, *p*_FDR_=0.011) (Table S8). ICC comparisons were similar (Table S9). Euler numbers, which provide an estimate of data quality, showed that images for each of the best-performing pipelines were of sufficient quality (Figure S5). When analyses were restricted to “good” volumes based on the visual quality assessment, LF-3T global correspondence results matched the full-sample results (Tables S10-S11, Figure S6).

**Figure 3.**
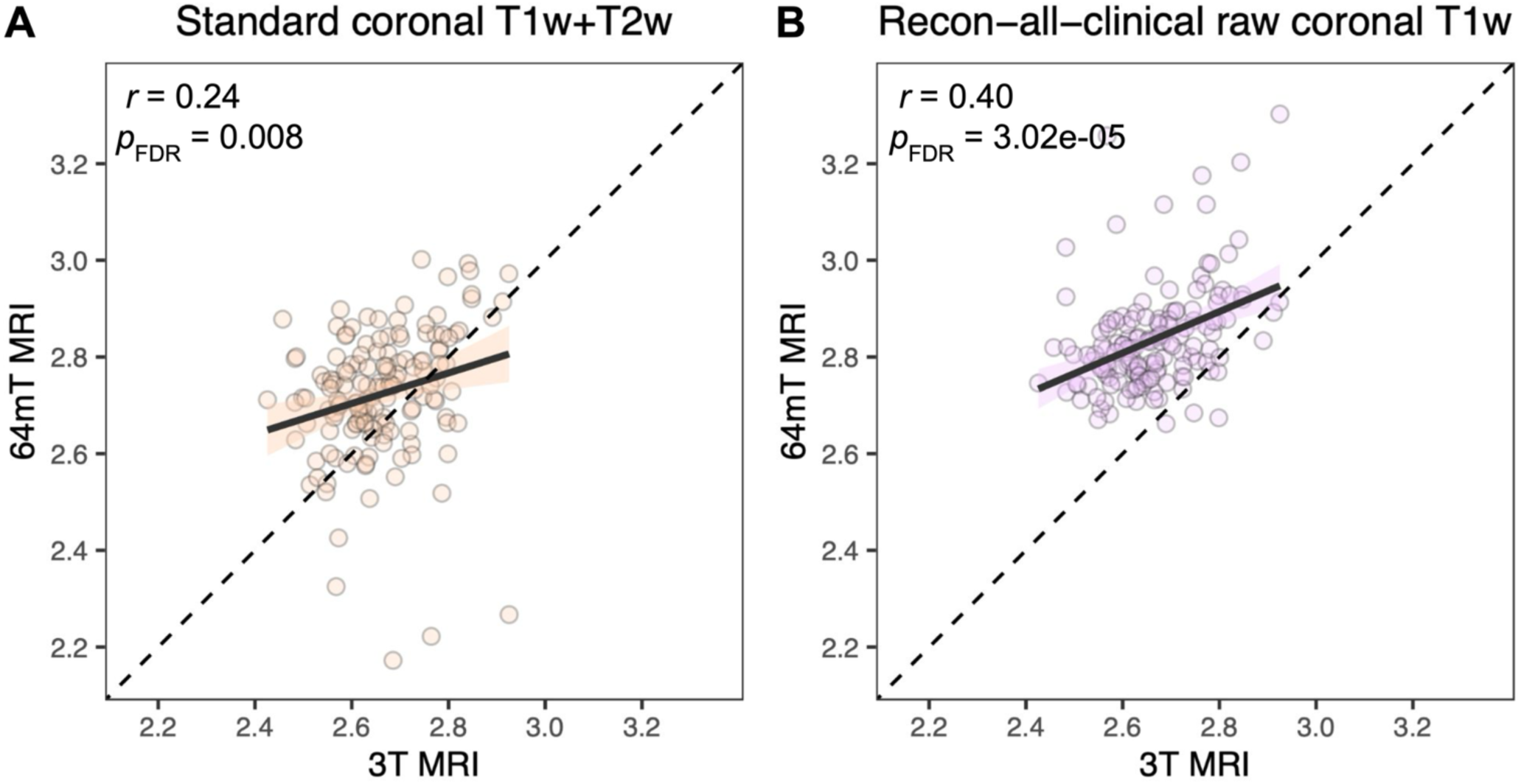
Global correlations between low-field and 3T measures. Scatter plots showing the relationship between global mean cortical thickness estimates derived from 3T and low-field MRI. The panel labels indicate the processing applied to the low-field images: (A) the standard coronal T1w+T2w pipeline and (B) the recon-all-clinical raw coronal T1w pipeline. The recon-all-clinical raw coronal T1w pipeline showed the highest global correspondence, whereas the standard coronal T1w+T2w pipeline was included to illustrate ‘baseline’ results from an image obtained with the same orientation and processed through recon-all without additional image processing. In both panels, the 3T reference estimates were derived from T1w and T2w images processed using recon-all. Each point represents one participant.

**Table 2.** Pearson correlation coefficients for the global correspondence between low-field and 3T images.

|  | Standard T1 and T2 |  |  | SynthSR v1 T1 and T2 |  |  |  |  |  |  |  |  |  |  |  |  |  |  |
| --- | --- | --- | --- | --- | --- | --- | --- | --- | --- | --- | --- | --- | --- | --- | --- | --- | --- | --- |
|  | *r* | *p* | *p*<sub>FDR</sub> | *r* | *p* | *p*<sub>FDR</sub> |  |  |  |  |  |  |  |  |  |  |  |  |
| axi | 0.24 | 0.003 | 0.008 | 0.31 | 9.85e-05 | 7.11e-04 |  |  |  |  |  |  |  |  |  |  |  |  |
| cor | 0.24 | 0.003 | 0.008 | 0.36 | 7.71e-06 | 1.25e-04 |  |  |  |  |  |  |  |  |  |  |  |  |
| sag | 0.27 | 7.41e-04 | 0.003 | 0.19 | 0.017 | 0.032 |  |  |  |  |  |  |  |  |  |  |  |  |
| multi | -0.06 | 0.464 | 0.472 | 0.23 | 0.005 | 0.012 |  |  |  |  |  |  |  |  |  |  |  |  |
|  | SynthSR v2 resampled T1 |  |  | SynthSR v2 raw T1 |  |  | SynthSR v2 resampled and co-registered T2 |  |  | SynthSR v2 co-registered T2 |  |  | SynthSR v2 resampled T2 |  |  | SynthSR v2 raw T2 |  |  |
|  | *r* | *p* | *p*<sub>FDR</sub> | *r* | *p* | *p*<sub>FDR</sub> | *r* | *p* | *p*<sub>FDR</sub> | *r* | *p* | *p*<sub>FDR</sub> | *r* | *p* | *p*<sub>FDR</sub> | *r* | *p* | *p*<sub>FDR</sub> |
| axi | 0.10 | 0.208 | 0.241 | 0.12 | 0.141 | 0.180 | 0.12 | 0.132 | 0.179 | 0.17 | 0.035 | 0.057 | 0.13 | 0.125 | 0.173 | 0.17 | 0.040 | 0.063 |
| cor | 0.21 | 0.011 | 0.023 | 0.33 | 2.89e-05 | 3.13e-04 | -0.16 | 0.046 | 0.070 | 0.16 | 0.045 | 0.070 | -0.10 | 0.225 | 0.257 | 0.12 | 0.137 | 0.180 |
| sag | 0.12 | 0.152 | 0.190 | 0.11 | 0.183 | 0.216 | — |  |  | 0.11 | 0.181 | 0.216 | — |  |  | — |  |  |
| multi | 0.18 | 0.026 | 0.044 | N/A |  |  | 0.23 | 0.005 | 0.012 | N/A |  |  | N/A |  |  | N/A |  |  |
|  | Recon-all-clinical resampled T1 |  |  | Recon-all-clinical raw T1 |  |  | Recon-all-clinical resampled and co-registered T2 |  |  | Recon-all-clinical co-registered T2 |  |  | Recon-all-clinical resampled T2 |  |  | Recon-all-clinical raw T2 |  |  |
|  | *r* | *p* | *p*<sub>FDR</sub> | *r* | *p* | *p*<sub>FDR</sub> | *r* | *p* | *p*<sub>FDR</sub> | *r* | *p* | *p*<sub>FDR</sub> | *r* | *p* | *p*<sub>FDR</sub> | *r* | *p* | *p*<sub>FDR</sub> |
| axi | 0.31 | 1.41e-04 | 9.17e-04 | 0.32 | 6.32e-05 | 5.14e-04 | 0.28 | 4.93e-04 | 0.002 | 0.35 | 9.74e-06 | 1.27e-04 | 0.28 | 4.74e-04 | 0.002 | 0.30 | 1.93e-04 | 0.001 |
| cor | 0.36 | 4.42e-06 | 9.58e-05 | 0.40 | 4.64e-07 | 3.02e-05 | 0.21 | 0.012 | 0.024 | 0.26 | 0.001 | 0.004 | 0.19 | 0.020 | 0.035 | 0.23 | 0.005 | 0.012 |
| sag | 0.12 | 0.141 | 0.184 | 0.20 | 0.012 | 0.024 | 0.21 | 0.009 | 0.019 | 0.22 | 0.006 | 0.013 | 0.01 | 0.940 | 0.940 | 0.24 | 0.003 | 0.008 |
| multi | 0.33 | 4.22e-05 | 3.92e-04 | N/A |  |  | 0.38 | 1.52e-06 | 4.94e-05 | N/A |  |  | N/A |  |  | N/A |  |  |
|  | Recon-any resampled T1 |  |  | Recon-any raw T1 |  |  | Recon-any resampled and co-registered T2 |  |  | Recon-any co-registered T2 |  |  | Recon-any resampled T2 |  |  | Recon-any raw T2 |  |  |
|  | *r* | *p* | *p*<sub>FDR</sub> | *r* | *p* | *p*<sub>FDR</sub> | *r* | *p* | *p*<sub>FDR</sub> | *r* | *p* | *p*<sub>FDR</sub> | *r* | *p* | *p*<sub>FDR</sub> | *r* | *p* | *p*<sub>FDR</sub> |
| axi | 0.08 | 0.346 | 0.375 | 0.06 | 0.465 | 0.472 | 0.16 | 0.054 | 0.080 | 0.08 | 0.332 | 0.372 | 0.25 | 0.002 | 0.006 | 0.13 | 0.120 | 0.170 |
| cor | 0.20 | 0.014 | 0.027 | 0.19 | 0.018 | 0.033 | 0.23 | 0.004 | 0.010 | 0.18 | 0.031 | 0.052 | 0.28 | 5.92e-04 | 0.003 | 0.25 | 0.002 | 0.006 |
| sag | 0.07 | 0.416 | 0.443 | 0.08 | 0.344 | 0.375 | 0.26 | 0.001 | 0.004 | 0.11 | 0.172 | 0.211 | 0.26 | 0.001 | 0.004 | 0.25 | 0.002 | 0.006 |
| multi | 0.14 | 0.080 | 0.116 | N/A |  |  | 0.06 | 0.445 | 0.467 | N/A |  |  | N/A |  |  | N/A |  |  |

Recon-all-clinical with raw coronal T1w showed significantly higher Pearson correlation coefficients than the best-performing pipeline in our previous study (Cooper et al., 2024), SynthSR v1 with axial T1w and T2w (Fisher’s *Z*=1.99, *p*=0.047), with a trend towards higher ICC (Fisher’s *Z*=1.92, *p*=0.055).

### 3.4 Lobar correspondence

Seven pipelines achieved the highest Pearson correlation coefficient in at least one cortical lobe (Figure 4A, Table 3). The recon-all-clinical multi T2w pipeline showed the highest correlations in the most lobes (N=4), followed by the recon-all-clinical multi T1w (N=2) and recon-all-clinical raw coronal T1w (N=2) pipelines (Table 3). ICC results were similar (Figure 4B, Table S12). There were no significant differences in the number of lobes with the highest correlations across pipelines. In the lobes where the recon-all-clinical multi T2w pipeline performed best (right frontal, right cingulate, and bilateral parietal), correlation coefficients ranged from *r*=0.34-0.51. The recon-all-clinical multi T1w pipeline (left cingulate and right temporal) showed top correlation coefficients ranging from *r*=0.41-0.46, and the recon-all-clinical raw coronal T1w pipeline (left frontal and left temporal) showed *r*=0.35 across both regions. Results were similar in participants under 18 years of age, as correlation coefficients ranged from *r*=0.39–0.64 for the recon-all-clinical multi T2w pipeline, *r*=0.37–0.59 for the recon-all-clinical multi T1w pipeline, and *r*=0.31–0.32 for the recon-all-clinical raw coronal T1w pipeline for the same lobar regions. Steiger’s Z-tests found no statistically significant differences between these pipelines (Figure S7). In the occipital lobe, the standard pipeline showed the highest correspondence across metrics (Table 3, Table S12).

**Figure 4.**
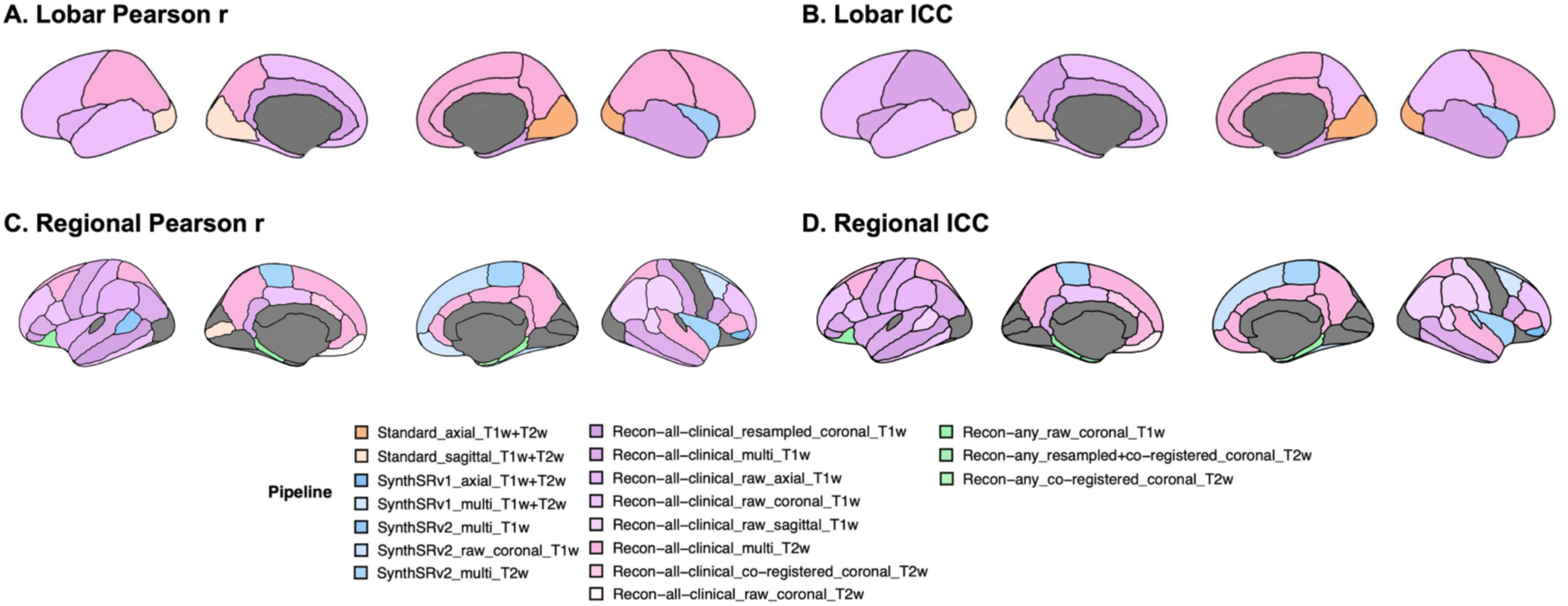
Lobar and regional correspondence between low-field (LF) and 3T measures: best performing pipeline. Lobar (A, B) and regional (C, D) correspondence were assessed using Pearson correlation coefficients (*r*) and intraclass correlation coefficients (ICCs). For each lobe and region, the processing pipeline yielding the highest cortical thickness correspondence is indicated. Analyses were restricted to pipelines that successfully processed all participants and remained significant after multiple-comparison correction at the global level (*p*_FDR_ < 0.05). Additional multiple testing correction was applied at the lobar and regional levels (*p*_FDR_ < 0.05); regions without assigned colors did not survive this correction.

**Table 3.**
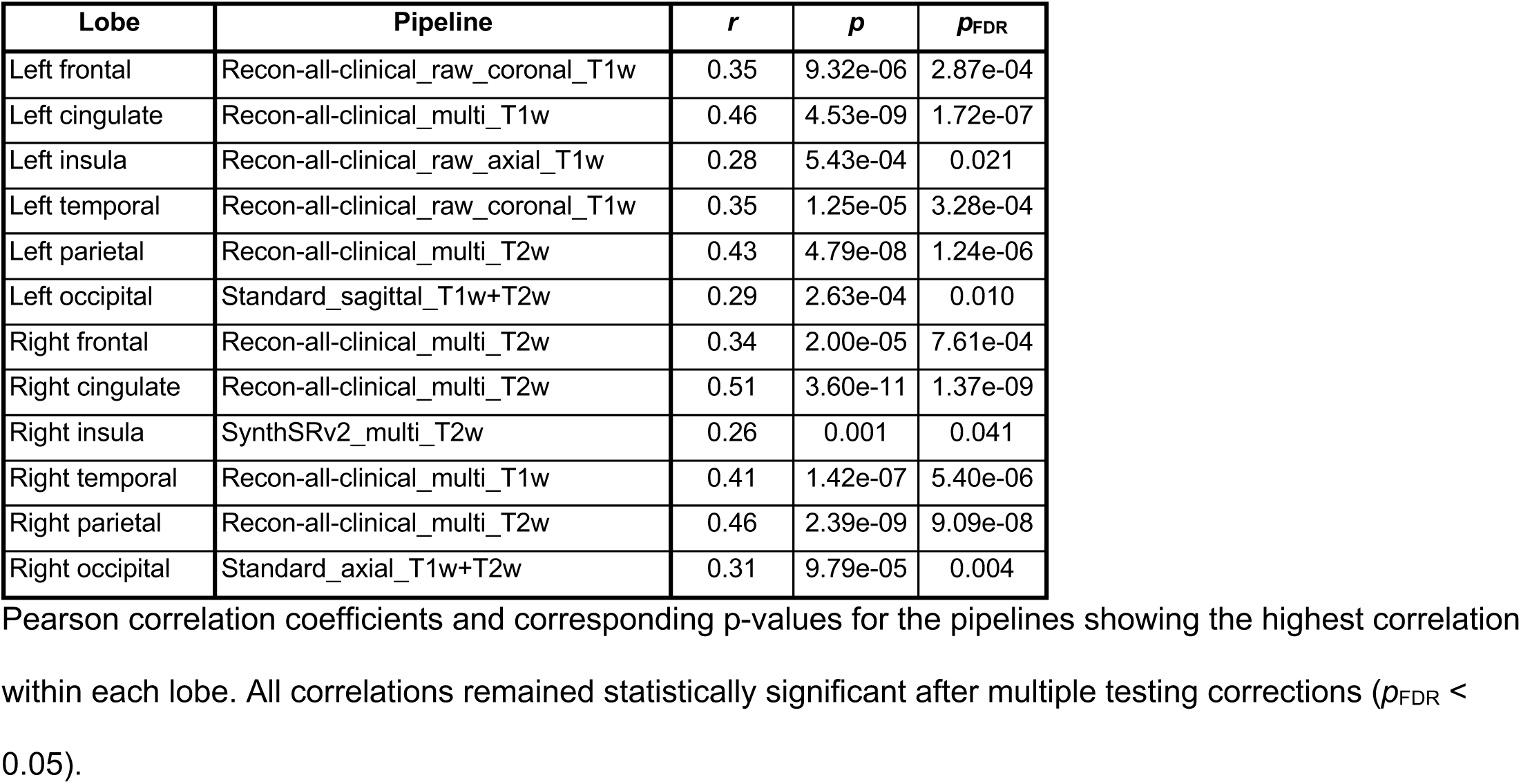
Pipelines with the highest Pearson correlation coefficients across lobes.

Compared with the standard pipeline, the recon-all-clinical multi T2w pipeline showed statistically significant improvements in the frontal and cingulate lobes (Tables S13-S14). Supplementary Results report findings for the two other top-performing pipelines.

### 3.5 Regional correspondence

Sixteen pipelines achieved the highest Pearson correlation coefficient in at least one cortical region (Figure 4C, Table 4A). The number of regions with the highest correlation coefficients differed across pipelines (*χ*²(9)=22.83, *p*=0.007); specifically, recon-all-clinical multi T2w pipeline (13 regions, *Z*=3.95, *p*=7.97e-05) and recon-all-clinical with multi T1w pipeline (9 regions, *Z*=2.02, *p*=4.33e-02) had more top-performing regions than other pipelines (Table 4A). ICC results were similar (Figure 4D, Table 4B). In the recon-all-clinical multi T2w pipeline, top correlations ranged from *r*=0.31-0.52. In the recon-all-clinical multi T1w pipeline, top correlations ranged from *r*=0.28-0.54. Comparable regional correlations were observed in participants under 18 years of age for the same regions, with correlations coefficients ranging from *r*=0.19–0.63 for the recon-all-clinical multi T2w pipeline and *r*=0.13-0.60 for the recon-all-clinical multi T1w pipeline. Direct comparisons between these two pipelines were largely non-significant (98.5%), indicating that most regions were measured with similar correspondence across pipelines (Supplemental Results).

**Table 4.** Number of regions showing the best correspondence between low-field and 3T measures for each processing pipeline.

| A. Pearson r |  |  | B. ICC |  |  |
| --- | --- | --- | --- | --- | --- |
| Pipeline | Region N | Std. residual | Pipeline | Region N | Std. residual |
| Recon-all-clinical_multi_T2w | 13 | 3.95 | Recon-all-clinical_multi_T2w | 14 | 4.07 |
| Recon-all-clinical_multi_T1w | 9 | 2.02 | Recon-all-clinical_multi_T1w | 10 | 2.22 |
| Recon-all-clinical_raw_coronal_T1w | 5 | 0.10 | Recon-all-clinical_raw_coronal_T1w | 5 | -0.09 |
| SynthSRv2_raw_coronal_T1w | 4 | -0.38 | Recon-all-clinical_raw_sagittal_T1w | 5 | -0.09 |
| SynthSRv2_multi_T2w | 3 | -0.87 | SynthSRv2_raw_coronal_T1w | 4 | -0.55 |
| SynthSRv1_multi_T1w+T2w | 3 | -0.87 | SynthSRv2_multi_T2w | 4 | -0.55 |
| Recon-any_raw_coronal_T1w | 3 | -0.87 | Recon-any_raw_coronal_T1w | 4 | -0.55 |
| Recon-all-clinical_raw_axial_T1w | 3 | -0.87 | SynthSRv1_multi_T1w+T2w | 2 | -1.48 |
| Recon-all-clinical_raw_sagittal_T1w | 3 | -0.87 | Recon-all-clinical_resampled_coronal_T1w | 2 | -1.48 |
| Recon-all-clinical_resampled_coronal_T1w | 2 | -1.35 | Recon-all-clinical_raw_axial_T1w | 2 | -1.48 |
| SynthSRv2_multi_T1w | 1 | - | SynthSRv1_axial_T1w+T2w | 1 | - |
| SynthSRv1_axial_T1w+T2w | 1 | - | Recon-any_co-registered_coronal_T2w | 1 | - |
| Recon-any_resampled+co-registered_coronal_T2w | 1 | - | Recon-all-clinical_co-registered_coronal_T2w | 1 | - |
| Recon-all-clinical_co-registered_coronal_T2w | 1 | - | Recon-all-clinical_raw_coronal_T2w | 1 | - |
| Recon-all-clinical_raw_coronal_T2w | 1 | - |  |  |  |
| Standard_sagittal_T1w+T2w | 1 | - |  |  |  |

Best correspondence was defined as the highest coefficient among the pipelines included in the regional analysis within each region, evaluated separately for Pearson correlation coefficients and intraclass correlation coefficients (ICCs). Standardized residuals were calculated from a chi-square test based on the number of dominant regions for each pipeline. Positive residuals indicate more dominant regions than expected and thus better performance in a greater number of regions, whereas negative residuals indicate fewer. Absolute values greater than 1.96 indicate a statistically significant deviation from the expected count ( $p < 0.05$ ).

The recon-all-clinical multi T2w pipeline, which performed best overall, had statistically significant improvements in 32 regions (47.1%) compared with the standard pipeline (Table S15). Greatest improvements were observed in the right caudal anterior cingulate, left parahippocampal, right medial orbitofrontal, and bilateral pars triangularis regions (Figure 5B, Table S15). ICC results were similar (Table S16). Regional improvements for recon-all-clinical with multi T1w, the second best performing pipeline, are reported in the Supplementary Results (Tables S17-S18).

**Figure 5.**
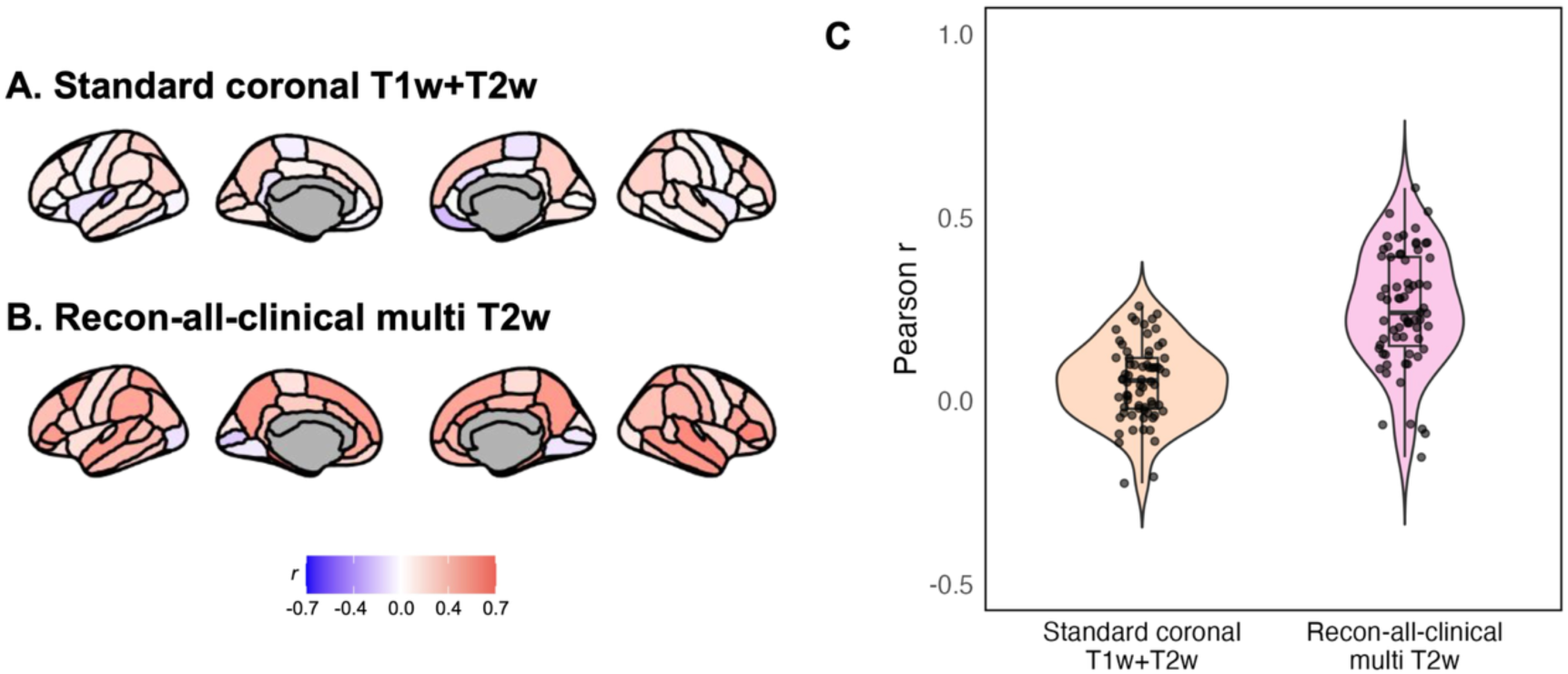
Regional correlations between low-field and 3T measures. Regional Pearson correlation coefficients (*r*) between LF and 3T measures derived from the standard coronal T1w and T2w pipeline (A) and the recon-all-clinical multi-orientation T2w pipeline (B). Correlations are color-coded, with positive values shown in red and negative values in blue. (C) Violin and box plots showing the distribution of regional Pearson correlation coefficients for the two pipelines. Each dot represents the Pearson correlation coefficient for an individual region.

Fisher’s Z-tests comparing regional correspondence between the best-performing pipeline in the current study (recon-all-clinical with multi T2w) with that from our previous study (Cooper et al., 2024) (SynthSR v1 with axial T1w and T2w) showed significant improvements in the left frontal pole, right caudal anterior cingulate, and right rostral anterior cingulate regions (Tables S19-S20).

### 3.6 Bland–Altman analyses

Bland–Altman analyses were conducted for the three best-performing pipelines: recon-all-clinical with raw coronal T1w, multi T1w, and multi T2w images. At the global level, these pipelines showed strong agreement with 3T estimates, with mean biases ranging from −0.17 to 0.00 mm (Figure S8). At the lobar level, mean biases across lobes were −0.19 mm (range=−0.29 to 0.18) for raw coronal T1w, −0.19 mm (range=−0.49 to −0.06) for multi T1w, and −0.08 mm (range=−0.42 to 0.17) for multi T2w (Figures S9–S11). At the regional level, mean biases across regions were −0.15 mm (range=−0.60 to 0.20) for raw coronal T1w, −0.20 mm (range=−0.61 to 0.19) for multi T1w, and −0.35 mm (range=−0.90 to 0.05) for multi T2w (Figures S12–S14). Overall, biases were predominantly negative, indicating that these pipelines generally overestimated cortical thickness relative to 3T, with greater biases observed in occipital areas.

## 4 Discussion

We examined 68 processing pipelines for estimating cortical thickness in LF MRI scans in a community sample of 150 young people. Recon-all-clinical pipelines using T1w or T2w images showed statistically significant improvements in global cortical thickness correspondence. At the lobar and regional levels, the recon-all-clinical multi T1w/T2w pipelines achieved the highest correlations in the greatest number of regions, with the largest improvements observed in the frontal, cingulate, and temporal regions. Top-performing pipelines showed significant improvements over the highest cortical thickness correspondence observed in our previous work (Cooper et al., 2024). In summary, our results demonstrate methodological improvements in the use of LF-MRI for assessing cortical thickness in young people, providing a quantitative benchmark for current tools in this area.

Cortical thickness is a key measure of brain development, and deviations from its normative trajectory have been proposed as biomarkers of psychopathology risk (Frangou et al., 2022; Hettwer et al., 2022; Luking et al., 2022). Improving its estimation from LF-MRI in young people is critical for expanding its applicability in research and clinical settings. Previously, we improved LF-HF correspondence in surface area and volume measures using super-resolution approaches, but not in cortical thickness (Cooper et al., 2024). Similarly, in a small sample of youth (N=12), LF-HF cortical thickness correspondence remained low (*r*≤0.3) across multiple processing methods (Pretzsch et al., 2025). However, in the current study, correlation coefficients reached up to *r*=0.40 globally, *r*=0.51 in the cingulate lobe, and *r*=0.58 in the parahippocampal region. The pipeline with the highest regional cortical thickness correspondence significantly outperformed the standard recon-all pipeline in 32 of 68 regions (*Z*=2.19-5.46; Table S15). In our previous work, SynthSR v1 significantly improved regional correspondence over the standard pipeline in 43 regions for surface area (*Z*=2.44-5.70) and 42 regions for cortical volume (*Z*=2.26-6.18). Although improvements in cortical thickness correspondence occurred in less regions in the current study, the magnitude of the improvements were similar, based on comparable Z-scores in surface area and volume estimates. Because we did not systematically evaluate surface area and cortical volume across the current pipelines, our results do not establish whether optimal processing strategies differ across cortical features. Across measures, recon-all-clinical pipelines showed overall superior performance, with only minor differences between sequences and orientations. The improvements likely reflect both methodological advances and the larger sample size, enabling more stable estimation of correspondence. Our results show that deep learning–based processing, particularly the recon-all-clinical pipeline applied to T1w or T2w images, substantially improves LF-HF correspondence.

The highest correspondence was achieved using the recon-all-clinical pipeline with multi T2w images, particularly in the frontal, cingulate, and temporal regions. These regions undergo protracted neurodevelopment (Gogtay et al., 2004) and are strongly implicated in psychiatric disorders (Collins et al., 2023; Van Erp et al., 2018), underscoring the relevance of these findings for neurodevelopmental and psychiatric research. Others have found that LF-MRI derived brain volume measures capture age effects comparable to those obtained from HF imaging (Deoni et al., 2021). Furthermore, LF-MRI estimated cortical thickness may be useful for constructing neuroimaging summary scores, which combine structural variations across multiple brain regions and have shown promise as neurobiological markers of psychosis (Rodrigue et al., 2024). Our findings demonstrate improved correspondence in regional cortical thickness and suggest potential for extending this line of work using LF-based measures. In contrast, correspondence remained relatively low in the insula and occipital regions, suggesting that estimates in these regions should be interpreted with caution.

Across cortical thickness measures, LF-HF correspondence from SynthSR v2-processed images remained low. These values were generally even lower than those obtained with SynthSR v1. SynthSR may introduce over-smoothing at the cortical surface, limiting sensitivity to individual variation (Gopinath et al., 2023). Also, conventional cortical surface reconstruction methods assign a single label per voxel (Dale et al., 1999), which may lead to over-smoothing, particularly at the gray–white matter boundary, where multiple tissue types may be present. In contrast, the recon-all-clinical pipeline does not rely on super-resolution and predicts voxel-wise signed distance functions from a given surface during surface reconstruction (Gopinath, Greve, et al., 2025). These methodological differences may explain recon-all-clinical’s superior performance in cortical thickness estimation. In addition, multi-orientation images did not improve cortical thickness correspondence relative to single-orientation images when processed without additional image processing (i.e., using the “standard” pipeline). Although multi-orientation averaging can improve isotropy and signal-to-noise ratio, these improvements may have been offset by boundary smoothing introduced through registration, interpolation, and averaging. With recon-all-clinical, however, correspondence for multi-orientation images was comparable to or higher than that obtained with single-orientation images. Recon-all-clinical, which is designed to accommodate heterogeneous image resolutions and contrasts, may better leverage the complementary information provided by multi-orientation images and produce more consistent tissue representations. In addition, certain acquisitions, processing pipelines, and brain regions remain challenging for LF cortical thickness estimation (see Supplementary Discussion).

Prior studies have suggested that T2w LF images may outperform T1w images for structural estimation (Gopinath, Sorby-Adams, et al., 2025; Váša et al., 2025), although this may vary by scanner type. Here, we did not find that one sequence consistently outperformed the other. T2w sequences often showed the highest numerical LF-HF correspondence; however, T1w-based pipelines using coronal scans or multi-orientation images frequently yielded comparable values, with overlapping confidence intervals. Therefore, T1w or T2w multi-orientation images and coronal T1w acquisitions may represent potentially optimal approaches for estimating cortical thickness using LF-MRI. When scan time is limited, coronal T1w scans may be a practical option. Furthermore, the optimal processing pipeline varies across brain regions, and pipeline selection can be tailored to the specific objective. For example, recon-all-clinical pipelines may be preferable for global and most regional estimates, whereas standard pipelines using recon-all without additional image processing may perform better in occipital regions. Figure S15 summarizes these findings to guide pipeline selection.

Several limitations should be noted. We used 3T MRI as the reference standard; however, cortical thickness estimates from 3T images are not a perfect ‘ground truth’ and are also subject to measurement error. Our top global cortical thickness correlations are in the range of *r*=0.3–0.4, and top regional correlates are *r*=0.2-0.6. These correlations represent weak-to-moderate associations, though they do demonstrate a significant improvement from what has been observed in previous LF-MRI findings in young people (Cooper et al., 2024). Importantly, the strength of these correlations is insufficient to demonstrate clinical utility or support disease-state classification. Other study limitations include the fact that newer versions of the Hyperfine software and alternative sequence options are now available. Future studies should evaluate newer versions of the Hyperfine software and alternative sequence options, as these improvements likely acquire images with higher SNR and could potentially further improve cortical thickness correspondence. It is highly likely that the 5-mm slice thickness of the current LF acquisitions increased partial-volume effects and limited accurate cortical boundary detection, reducing our ability to accurately estimate cortical thickness. Higher-resolution and isotropic acquisitions likely reduce these effects. The processing tools were not specifically developed for pediatric populations, which may limit their performance in youth samples. Processing tools tailored to youth populations may further improve correspondence in this group. Also, although we evaluated several widely used processing approaches, other methods are available. Deep learning–based tools trained on empirical LF data (Lucas et al., 2025) or specifically on pediatric empirical LF data (Baljer, Briski, et al., 2025; Baljer, Zhang, et al., 2025) may achieve better performance and should be explored in future work. Newer super-resolution and synthesis tools such as SuperSynth (https://surfer.nmr.mgh.harvard.edu), which generate multiple image contrasts, also warrant evaluation. Finally, we used a development version of recon-any (Gopinath, Sorby-Adams, et al., 2025), and its performance may differ in stabilized release. Further studies are needed to evaluate correspondence as updated versions become available.

We find that using recon-all-clinical pipelines on LF-MRIs can achieve moderate cortical thickness correspondence with 3T MRI in a large sample of young people. Correspondence varied across brain regions depending on image acquisition factors and processing approaches, highlighting the importance of selecting appropriate pipelines for specific objectives. These results demonstrate progress in LF-MRI for cortical thickness estimation in young people and highlight its potential for further development as a scalable and accessible neuroimaging tool.

## Data and code availability

The data and code that support the findings of this study are available from the corresponding author upon reasonable request.

## Author contributions

S.C. and M.J. contributed to conceptual development of the study, data collection, analysis, interpretation, and manuscript writing; J.S., R.C., M.C., S.S., and I.E. contributed to data collection, analysis, interpretation, and manuscript editing. R.H. contributed to data processing and manuscript editing. A.L., C.V., and J.M.S. contributed to interpretation and manuscript editing.

