## Supplementary Materials for "Processing strategies for improving cortical thickness correspondence between low-field and high-field MRI in young people"

**Supplementary Material**

**Supplementary Methods**

**Supplementary Results**

**Supplementary Discussion**

**Figure S1.** A representative example of outputs reviewed during visual quality control

**Figure S2.** Examples of poor-quality images due to artifacts

**Figure S3.** Pipeline characteristics for global Pearson correlation analyses

**Figure S4.** Pipeline characteristics for global intraclass correlation coefficient (ICC) analyses

**Figure S5.** Distribution of Euler numbers for 3T images and the best-performing low-field processing pipelines

**Figure S6.** Global mean cortical thickness correspondence between low-field and 3T MRI pre- and post-quality filtering

**Figure S7.** Lobar correspondence between low-field and 3T MRI for the best-performing pipelines

**Figure S8.** Bland–Altman plots of agreement between global cortical thickness estimates derived from 3T and low-field MRI

**Figure S9.** Bland–Altman plots of agreement between lobar cortical thickness estimates derived from 3T and low-field MRI processed using recon-all-clinical with raw coronal T1w images

**Figure S10.** Bland–Altman plots of agreement between lobar cortical thickness estimates derived from 3T and low-field MRI processed using recon-all-clinical with multi T1w images

**Figure S11.** Bland–Altman plots of agreement between lobar cortical thickness estimates derived from 3T and low-field MRI processed using recon-all-clinical with multi T2w images

**Figure S12.** Bland–Altman plots of agreement between regional cortical thickness estimates derived from 3T and low-field MRI processed using recon-all-clinical with raw coronal T1w images

**Figure S13.** Bland–Altman plots of agreement between regional cortical thickness estimates derived from 3T and low-field MRI processed using recon-all-clinical with multi T1w images

**Figure S14.** Bland–Altman plots of agreement between regional cortical thickness estimates derived from 3T and low-field MRI processed using recon-all-clinical with multi T2w images

**Figure S15.** Decision flowchart for selecting low-field MRI processing pipelines for cortical thickness estimation

**Table S1.** Scan parameters for the 64mT and 3T scans acquired in this study

**Table S2.** Description of processing approaches

**Table S3.** Low-field pipelines tested in this study

**Table S4.** Proportion of poor-quality images by processing pipeline

**Table S5.** Low-field pipelines with significant global Pearson correlations with 3T

**Table S6.** Low-field pipelines with significant global intraclass correlations with 3T

**Table S7.** Intraclass correlation coefficients (ICCs) for the global correspondence between low-field and 3T images

**Table S8.** Comparison of low-field pipeline group–average Pearson correlation coefficients

**Table S9.** Comparison of low-field pipeline group–average intraclass correlation coefficients (ICCs)

**Table S10.** Pearson correlations between processed low-field and 3T global measures using only good-quality images

**Table S11.** Intraclass correlations between processed low-field and 3T global measures using only good-quality images

**Table S12.** Pipelines with the highest Intraclass correlation coefficients (ICCs) across lobes

**Table S13.** Pearson correlations of lobar measures from standard versus recon-all-clinical-processed multi T2w with 3T scans

**Table S14.** Intraclass correlations of lobar measures from standard versus recon-all-clinical-processed multi T2w with 3T scans

**Table S15.** Pearson correlations of regional measures from standard versus recon-all-clinical-processed multi T2w with 3T scans

**Table S16.** Intraclass correlations of regional measures from standard versus recon-all-clinical-processed multi T2w with 3T scans

**Table S17.** Pearson correlations of regional measures from standard versus recon-all-clinical-processed multi T1w with 3T scans

**Table S18.** Intraclass correlations of regional measures from standard versus recon-all-clinical-processed multi T1w with 3T scans

**Table S19.** Regional Pearson correlation improvements: current vs. previous best-performing pipelines

**Table S20.** Regional intraclass correlation improvements: current vs. previous best-performing pipelines

**Supplementary References**

### Supplementary Methods

*Selection of processing approaches*

The selection of tools and processing approaches was guided by the iterative development of the study rather than by a formal a priori ranking or a predefined comprehensive list of available methods. The project initially began as a direct comparison of SynthSR v1 and SynthSR v2, motivated by the release of SynthSR v2 and the limited comparison between the two approaches in the original SynthSR manuscript (Iglesias et al., 2023), which we learned of after submitting our first low-field MRI manuscript (Cooper et al., 2024). Although previous adult studies reported similar or modestly improved volumetric performance with SynthSR v2 (Iglesias et al., 2023; Sorby-Adams et al., 2024), they did not directly compare correlation strengths between versions, and no prior study had compared the two versions for cortical thickness estimation in youth.

As the project developed, we consulted the SynthSR and FreeSurfer documentation to identify recommended approaches for key preprocessing decisions, including image resampling, co-registration, and the incorporation of multiple MRI sequence types and image orientations. To our knowledge, none of these methodological decisions had been systematically tested in the past; thus, we expanded our analyses to systematically evaluate the effects of these choices within processing pipelines.

During data processing and analysis, we also became aware of the FreeSurfer recon-all-clinical and recon-any reconstruction approaches and incorporated these methods into our comparisons. The resulting combination of methodological choices yielded 68 distinct processing pipelines. At that stage, we considered the study scope sufficiently broad and restricted the study to these pipelines, rather than continuing to incorporate additional tools or approaches as they became available.

*Post-processing quality assessment of LF outputs*

J.S. conducted the primary assessment, and ambiguous cases were resolved by consensus. We reviewed the output volumes (derived from resampled images for T1-weighted (T1w)-only pipelines with a single orientation and from resampled and co-registered images for all other pipelines) and evaluated image quality across imaging sequences, orientations, and processing approaches (Figure S1). For SynthSR-processed pipelines, we reviewed the output volumes generated after image synthesis. For recon-all-clinical and recon-any-processed pipelines, we reviewed the cortical reconstruction. We rated image quality on a three-point scale (1=poor, 2=fair, 3=good), with good-quality volumes defined by clearly identifiable cortical anatomy and absence of noticeable artifacts, signal loss within the cerebrum, or excessive background noise. Poor-quality volumes were defined by substantial artifacts, signal loss, or excessive noise that obscured cortical anatomy. Based on these ratings, we examined correspondence with 3T using (i) all outputs and (ii) only successful outputs, excluding volumes rated as poor or fair.

*Multiple testing correction*

For global correspondence, we applied the false discovery rate (FDR) correction across pipelines. For analyses examining correspondence across all lobes or regions within each pipeline, FDR correction was applied across lobes or regions. For analyses evaluating multiple pipelines within each lobe or region, FDR correction was applied across pipelines. Similarly, when testing improvements over the standard pipeline or the best-performing pipeline from our previous study across lobes or regions, we applied correction across lobes or regions for each comparison.

**Supplementary Results**

*Comparison of 3T reference pipelines with and without auxiliary T2w input*

To evaluate the effect of including T2-weighted (T2w) images as auxiliary inputs in the 3T reference pipeline, we compared cortical thickness estimates derived using both T1w and T2w images with those derived from T1w images alone. The two approaches showed high correspondence globally ($r$=0.91; ICC=0.91), across lobes ($r$=0.86-0.96; ICC=0.86-0.96), and across cortical regions ($r$=0.74–0.98; ICC=0.73-0.98), indicating that the 3T estimates were highly consistent with and without auxiliary T2w input.

*Pipeline failures*

Cortical thickness estimation failed for one participant in the SynthSR v2 resampled and co-registered sagittal T2w pipeline, three participants in the SynthSR v2 resampled sagittal T2w pipeline, and two participants in the SynthSR v2 raw sagittal T2w pipeline. The errors occurred during topology correction, Talairach registration, and skull stripping or white matter identification.

*Lobar correspondence*

We used Steiger's Z tests to compare the recon-all-clinical multi T1w and recon-all-clinical raw coronal T1w pipelines—which showed the second-best performance in lobar correspondence—with the standard processing pipeline. The recon-all-clinical multi T1w pipeline showed statistically significant improvements in the bilateral cingulate (left: $Z$=4.10, $p_{FDR}$=5.01e-04; right: $Z$=3.82, $p_{FDR}$=7.86e-04), bilateral temporal lobes (left: $Z$=2.44, $p_{FDR}$=0.037; right: $Z$=2.42, $p_{FDR}$=0.037), and left insula ($Z$=2.96, $p_{FDR}$=0.012) based on Pearson correlation coefficients. ICC results showed a similar pattern (left cingulate: $Z$=4.09, $p_{FDR}$=5.14e-04; left insula: $Z$=2.78, $p_{FDR}$=0.021; left temporal: $Z$=2.46, $p_{FDR}$=0.035; right cingulate: $Z$=3.81, $p_{FDR}$=8.27e-04; right temporal: $Z$=2.44, $p_{FDR}$=0.035). The recon-all-clinical raw coronal T1w pipeline showed statistically significant improvements in the bilateral cingulate (left: $Z$=3.85, $p_{FDR}$=7.02e-04; right: $Z$=4.15, $p_{FDR}$=3.94e-04), bilateral temporal lobes (left: $Z$=2.48, $p_{FDR}$=0.038; right: $Z$=2.38, $p_{FDR}$=0.038), bilateral insula (left: $Z$=2.51, $p_{FDR}$=0.038; right: $Z$=2.29, $p_{FDR}$=0.038), and left frontal lobe ($Z$=2.31, $p_{FDR}$=0.038) based on Pearson correlation coefficients. ICC results showed a similar pattern (left frontal: $Z$=2.35, $p_{FDR}$=0.046; left cingulate: $Z$=3.82, $p_{FDR}$=7.97e-04; left temporal: $Z$=2.46, $p_{FDR}$=0.044; left insula: $Z$=2.26, $p_{FDR}$=0.046; right cingulate: $Z$=4.05, $p_{FDR}$=6.14e-04; right temporal: $Z$=2.44, $p_{FDR}$=0.044; right parietal: $Z$=2.16, $p_{FDR}$=0.046; right insula: $Z$=2.17, $p_{FDR}$=0.046).

*Regional correspondence*

We performed Steiger's Z tests to compare regional correspondence between the recon-all-clinical multi T2w pipeline, which showed the best performance, and the recon-all-clinical multi T1w pipeline, which showed the second-best performance. A statistically significant difference between the two pipelines was observed only in the right banks of the superior temporal sulcus, where the recon-all-clinical multi T1w pipeline showed higher correspondence than the recon-all-clinical multi T2w pipeline (Pearson's $r$: Steiger's $Z$=3.71, $p_{FDR}$=0.014; ICC: Steiger's $Z$=3.68, $p_{FDR}$=0.016).

When we used Steiger's Z tests to compare regional improvements relative to the standard processing pipeline, the recon-all-clinical multi T1w pipeline (the second-best-performing pipeline) showed statistically significant improvements in 33 regions (48.5% of all regions), including ten frontal, six cingulate, nine temporal, seven parietal, and one occipital regions (Table S17). The five strongest improvements were observed in the bilateral parahippocampal regions, left pars opercularis, right caudal anterior cingulate, and right middle temporal regions. ICC results were similar (Table S18).

**Supplementary Discussion**

This study also identified acquisitions, processing pipelines, and brain regions that remain challenging for LF cortical thickness estimation. Sagittal acquisitions showed lower correspondence after processing, with more output artifacts and processing failures. Preprocessing effects were minimal for T1w images, whereas processing failures were more frequent for T2w images without co-registration. Compared to recon-all-clinical pipelines, SynthSR v2 and recon-any pipelines produced significantly lower cortical thickness correspondence. SynthSR v2 and recon-any showed correspondence comparable to that of the standard pipeline using recon-all without additional image processing, providing no clear evidence that these alternative approaches offered an advantage over the standard pipeline. Quality assessment revealed blurring and signal loss in outputs from some pipelines, particularly in occipital regions near the cerebellum, suggesting that current processing methods are not well optimized for cerebellar regions and may affect adjacent cortical areas. Lobar and regional analyses further showed that standard pipelines without additional processing yielded the highest correspondence in these regions, indicating that they remain particularly challenging to process. Based on quality assessment, we compared correspondence using all processed volumes versus only those of good quality and found largely consistent results. This suggests that when high-quality data are selected prior to processing, post-processing quality control has limited impact on improving group-level estimation accuracy.

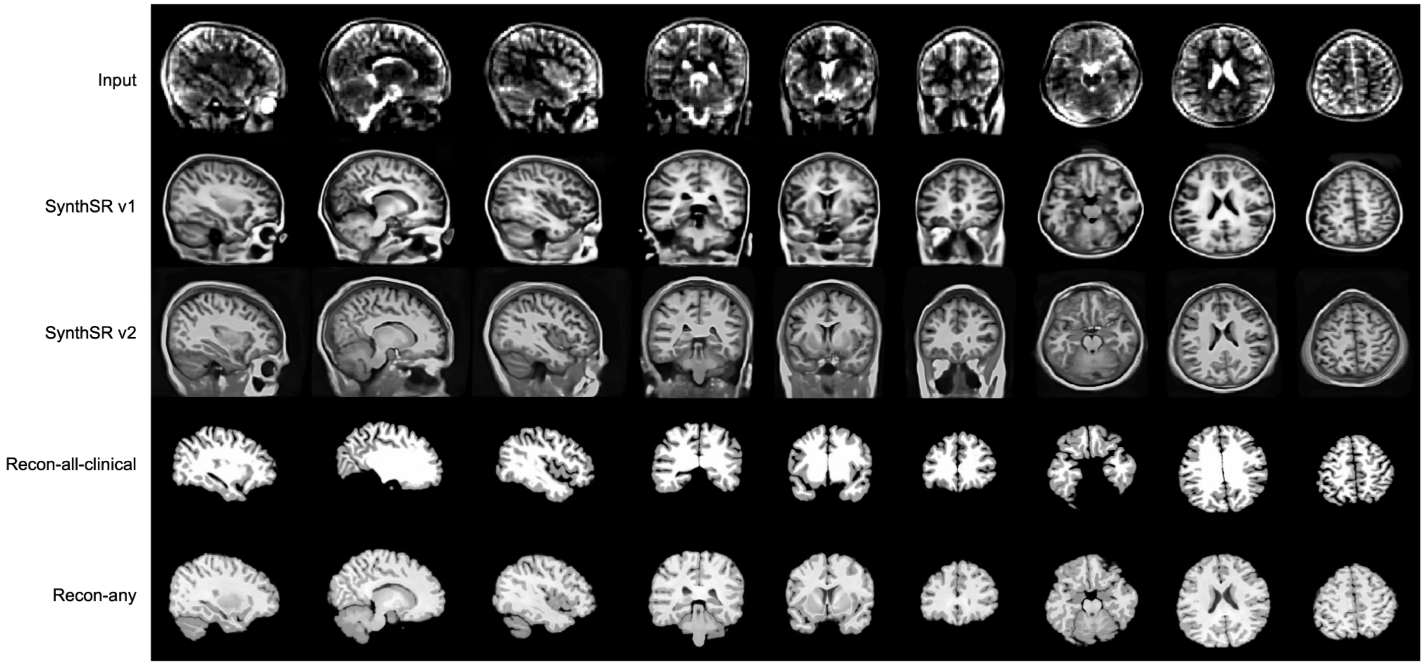


**Figure S1. A representative example of outputs reviewed during visual quality control.** The first row shows the input images, and subsequent rows show the corresponding outputs generated using SynthSR v1, SynthSR v2, recon-all-clinical, and recon-any, respectively. These outputs were visually assessed for the clarity of cortical anatomy and the presence of artifacts, signal loss within the cerebrum, or excessive background noise.

A

B

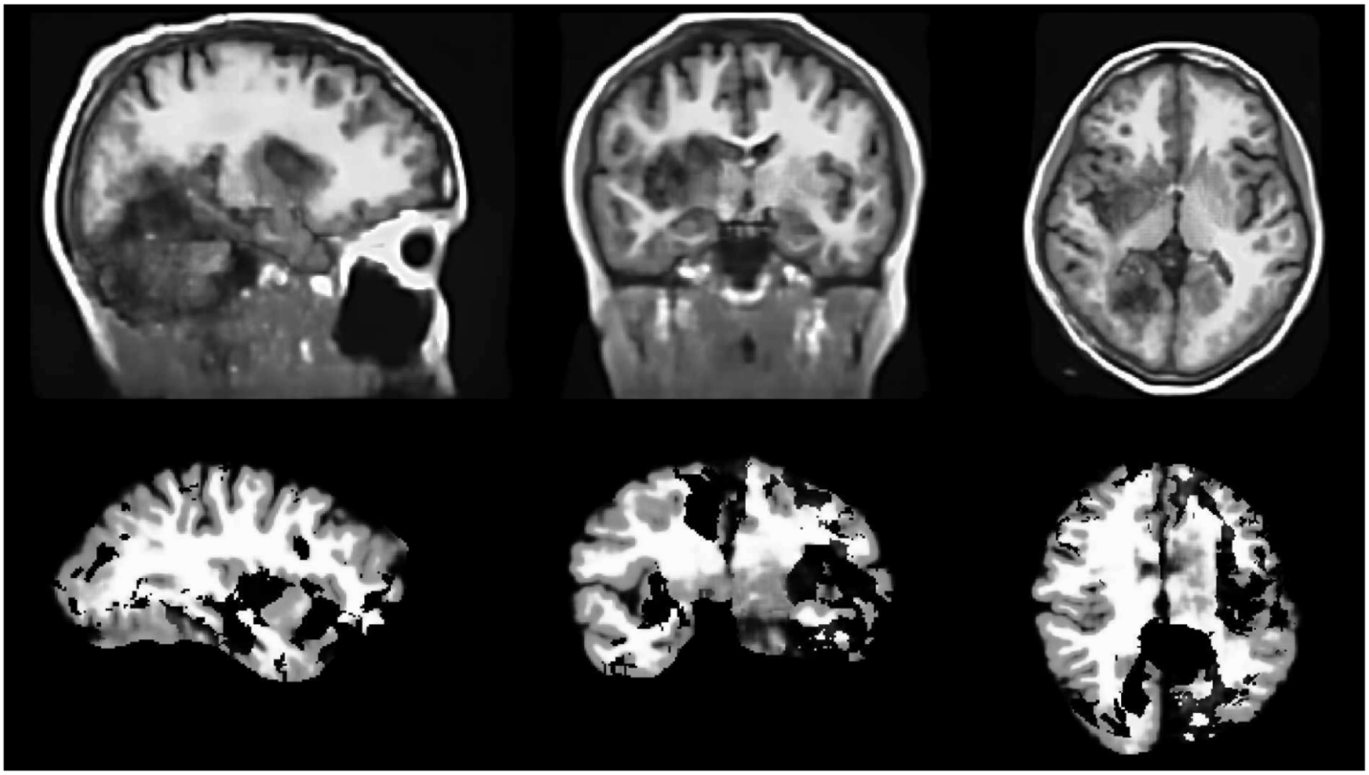

**Figure S2. Examples of poor-quality images due to artifacts.** (A) Blurring in occipital and subcortical regions following SynthSR v2 synthesis. (B) Widespread signal loss across the brain after recon-all-clinical reconstruction.

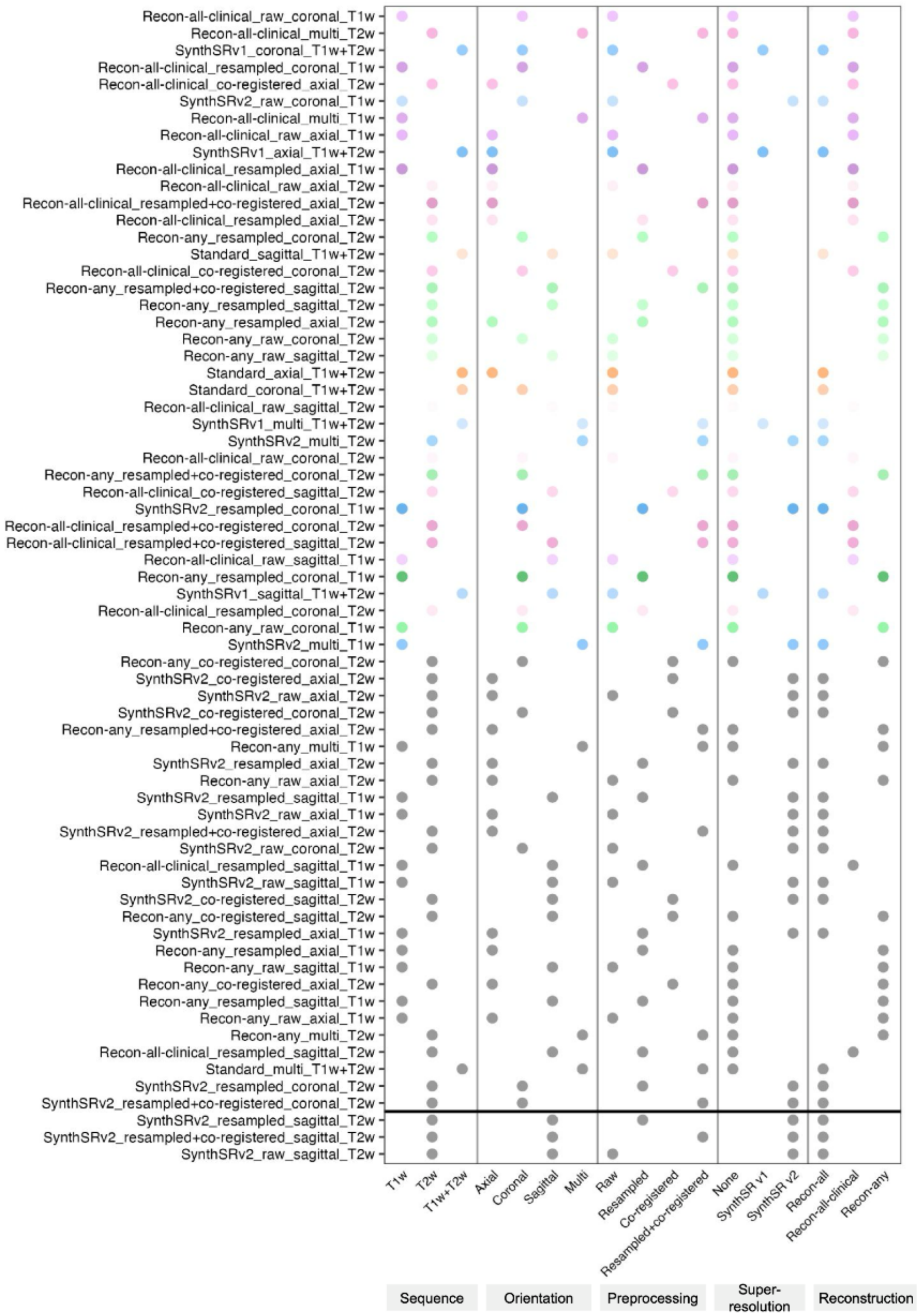


**Figure S3. Pipeline characteristics for global Pearson correlation analyses.** Pipelines are ordered by descending global Pearson correlation coefficient, consistent with the order in Figure 2A. Pipelines above the horizontal divider were successfully completed for all participants, whereas those below the divider failed for a subset of participants. Gray indicates pipelines with non-significant correspondence.

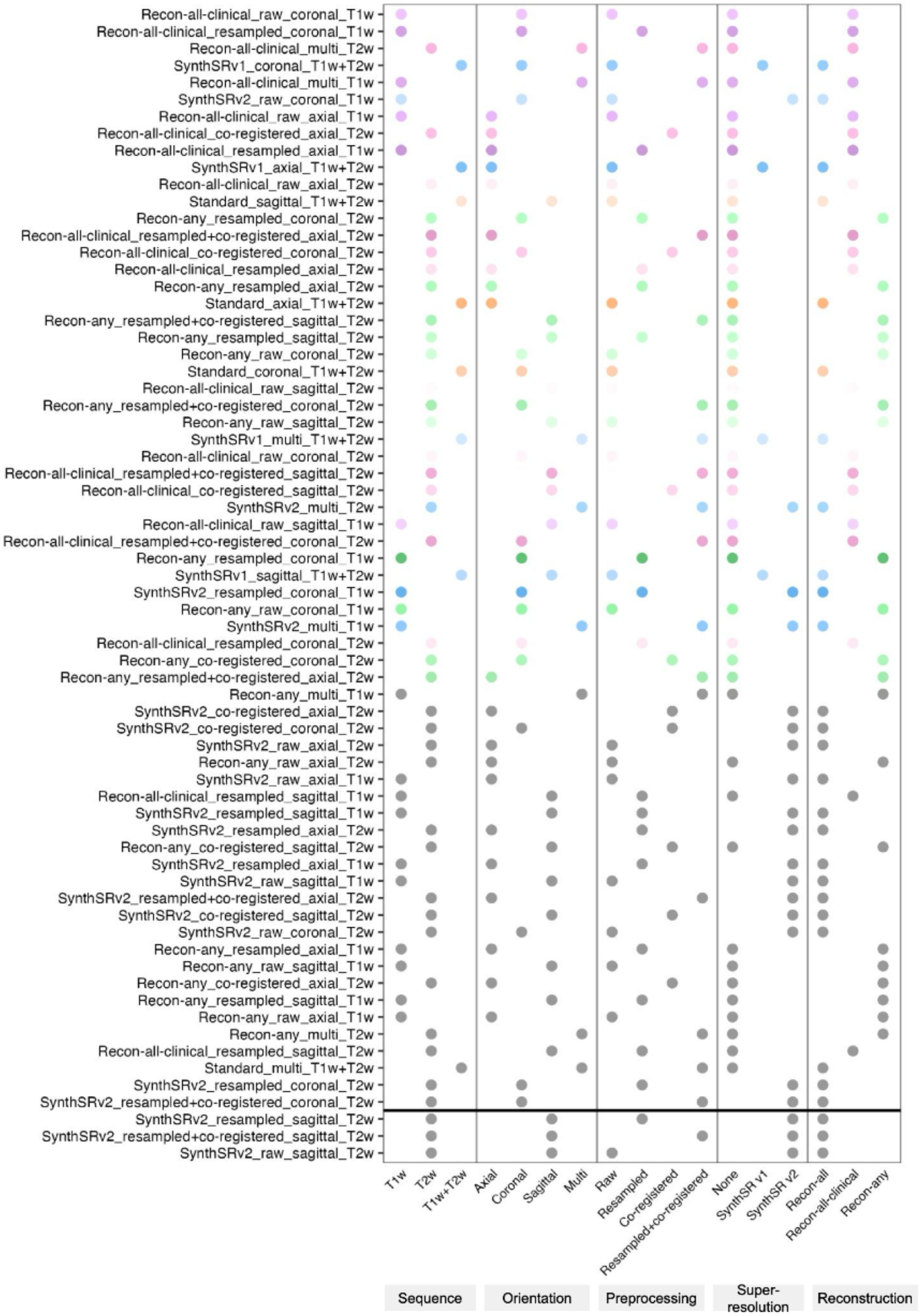


**Figure S4. Pipeline characteristics for global intraclass correlation coefficient (ICC) analyses.** Pipelines are ordered by descending global ICC, consistent with the order in Figure 2B. Pipelines above the horizontal divider were successfully completed for all participants, whereas those below the divider failed for a subset of participants. Gray indicates pipelines with non-significant correspondence.

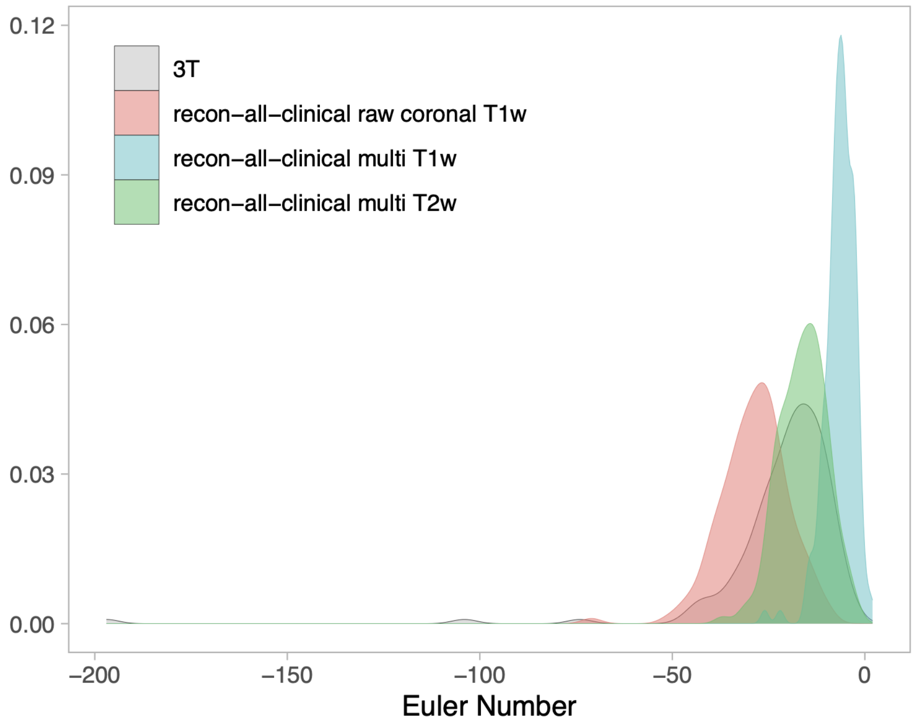


**Figure S5. Distribution of Euler numbers for 3T images and the best-performing low-field processing pipelines.** Density plots show Euler numbers for the 3T reference images and low-field images processed using recon-all-clinical with raw coronal T1w, multi T1w, or multi T2w images. The Euler number reflects topological complexity of the reconstructed cortical surface, and can be used as an index of data quality. A Euler number less than -217 indicates poor image quality (Rosen et al., 2018).

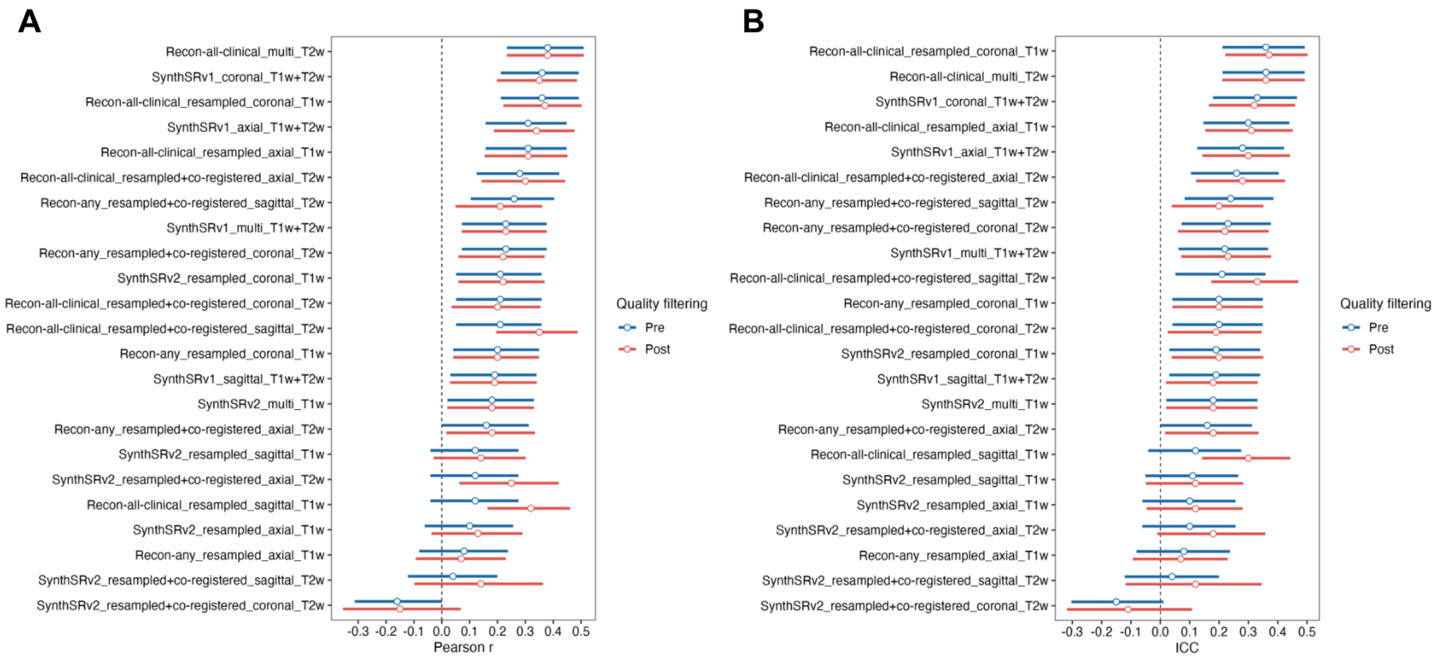


**Figure S6. Global mean cortical thickness correspondence between low-field and 3T MRI pre- and post-quality filtering.** For low-field MRI pipelines that included fair or poor-quality images, analyses were performed both including all images and after restricting to good-quality images only. (A) Pearson correlation coefficients and (B) intraclass correlation coefficients (ICCs).

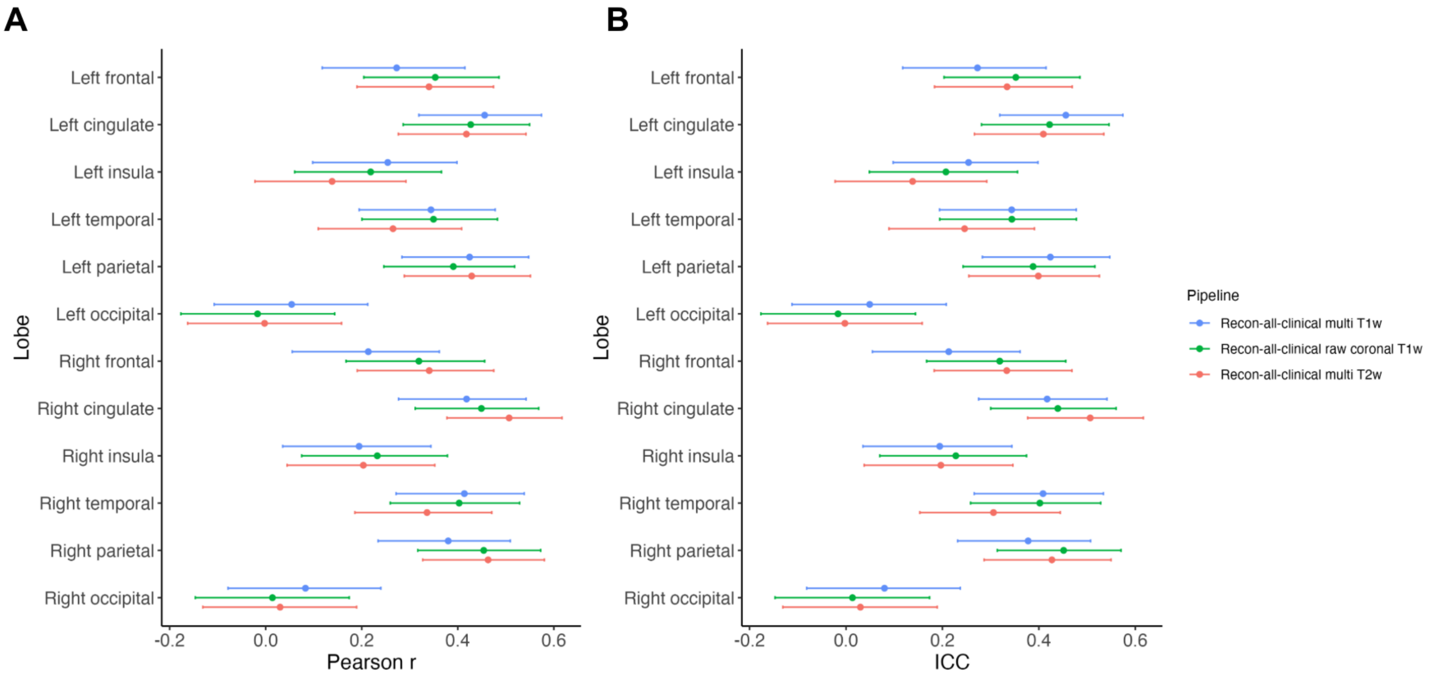


**Figure S7. Lobar correspondence between low-field and 3T MRI for the best-performing pipelines.**

Pipelines that showed the highest correspondence across the greatest number of lobes are presented. (A) Pearson correlation coefficients and (B) intraclass correlation coefficients (ICCs).

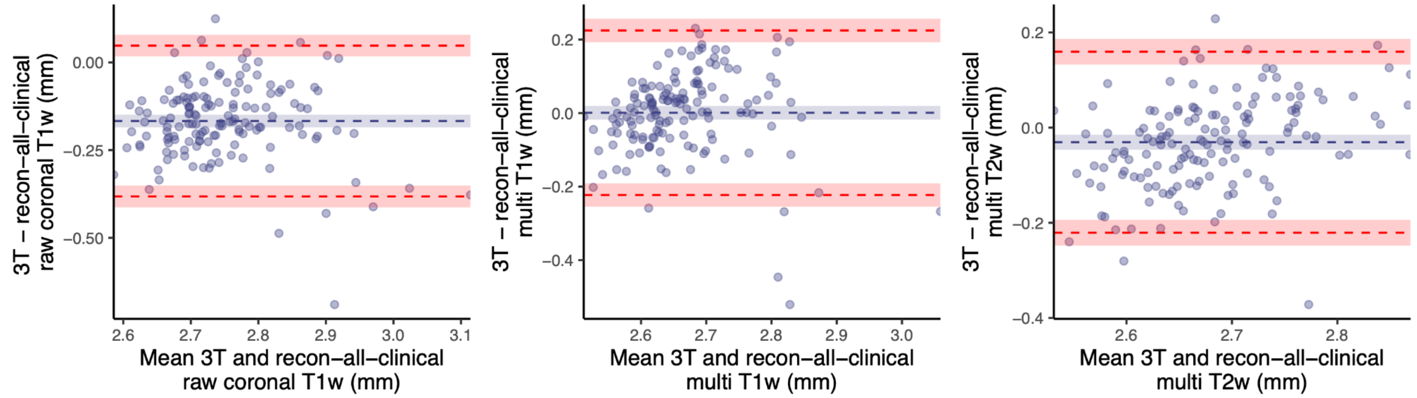


**Figure S8. Bland–Altman plots of agreement between global cortical thickness estimates derived from 3T and low-field MRI.** Panels show results for recon-all-clinical with (A) raw coronal T1w, (B) multi T1w, and (C) multi T2w images. Each point represents one participant. The blue dashed line indicates the mean bias, and the upper and lower red dashed lines indicate the 95% limits of agreement. Shaded regions indicate the corresponding 95% confidence intervals. Differences were calculated as 3T minus low-field estimates; therefore, negative values indicate higher cortical thickness estimates from low-field MRI than from 3T MRI.

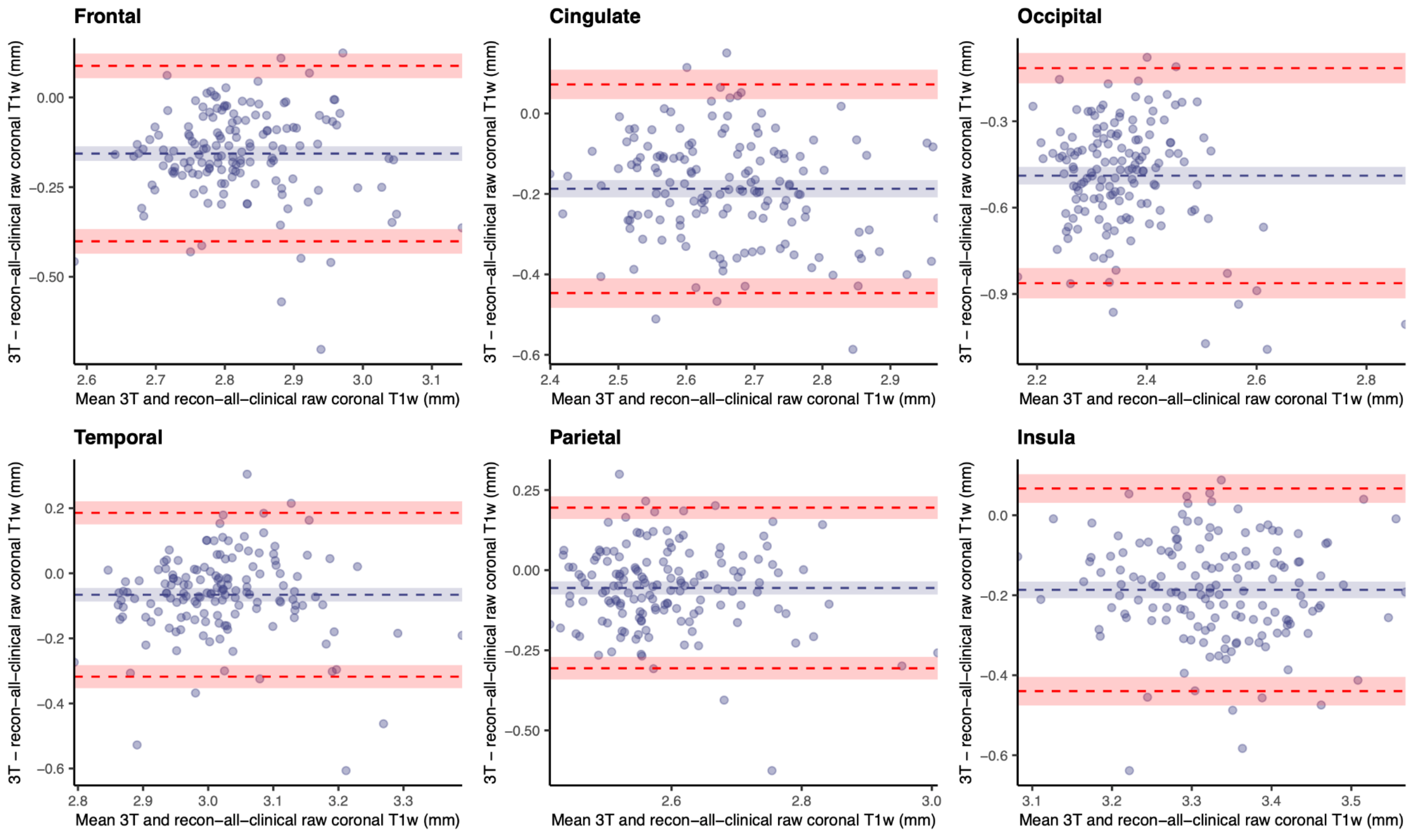


**Figure S9. Bland–Altman plots of agreement between lobar cortical thickness estimates derived from 3T and low-field MRI processed using recon-all-clinical with raw coronal T1w images.** Plots show results for the frontal, cingulate, occipital, temporal, parietal, and insular lobes. Each point represents one participant. The blue dashed line indicates the mean bias, and the upper and lower red dashed lines indicate the 95% limits of agreement. Shaded regions indicate the corresponding 95% confidence intervals. Differences were calculated as 3T minus low-field estimates; therefore, negative values indicate higher cortical thickness estimates from low-field MRI than from 3T MRI.

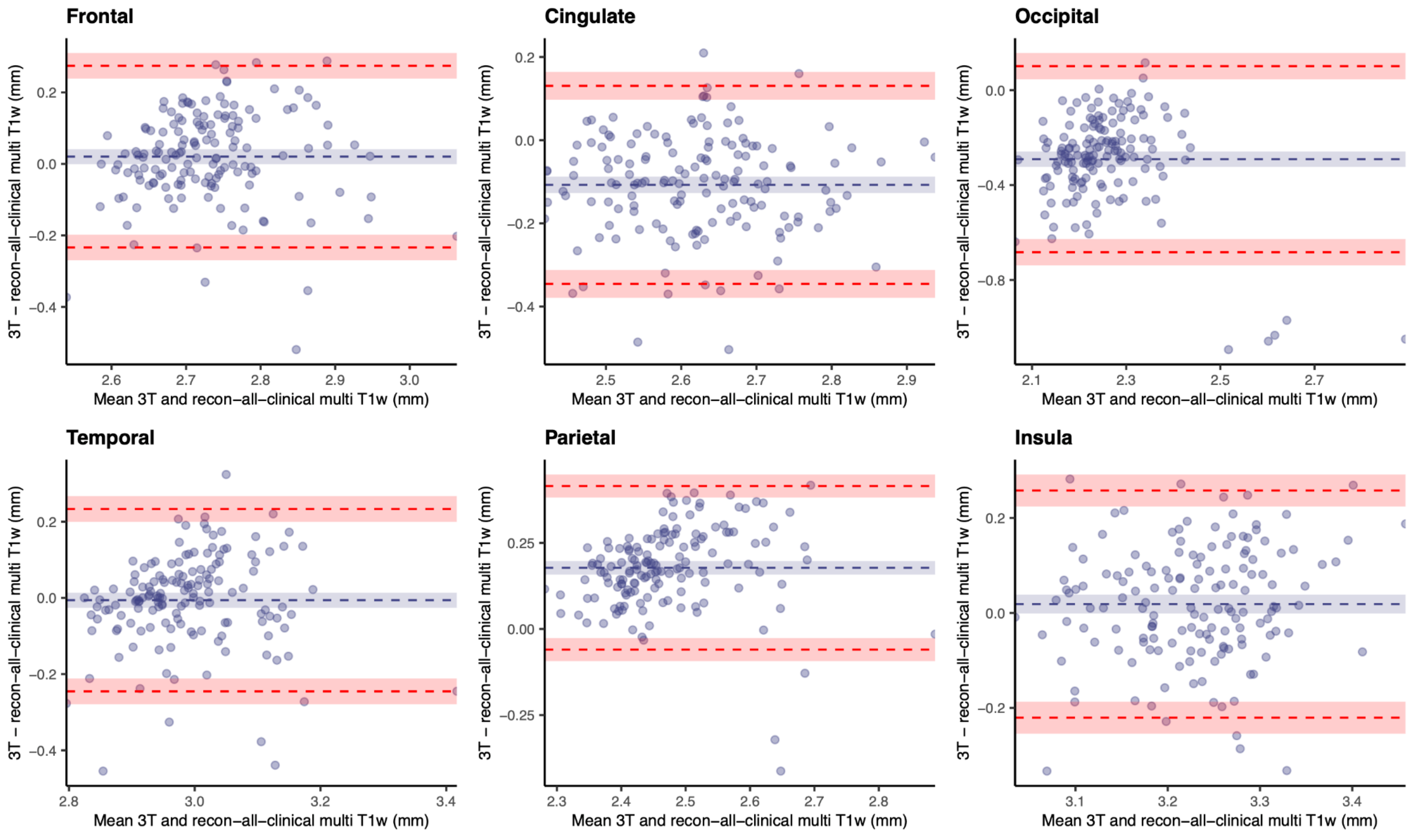


**Figure S10. Bland–Altman plots of agreement between lobar cortical thickness estimates derived from 3T and low-field MRI processed using recon-all-clinical with multi T1w images.** Plots show results for the frontal, cingulate, occipital, temporal, parietal, and insular lobes. Each point represents one participant. The blue dashed line indicates the mean bias, and the upper and lower red dashed lines indicate the 95% limits of agreement. Shaded regions indicate the corresponding 95% confidence intervals. Differences were calculated as 3T minus low-field estimates; therefore, negative values indicate higher cortical thickness estimates from low-field MRI than from 3T MRI.

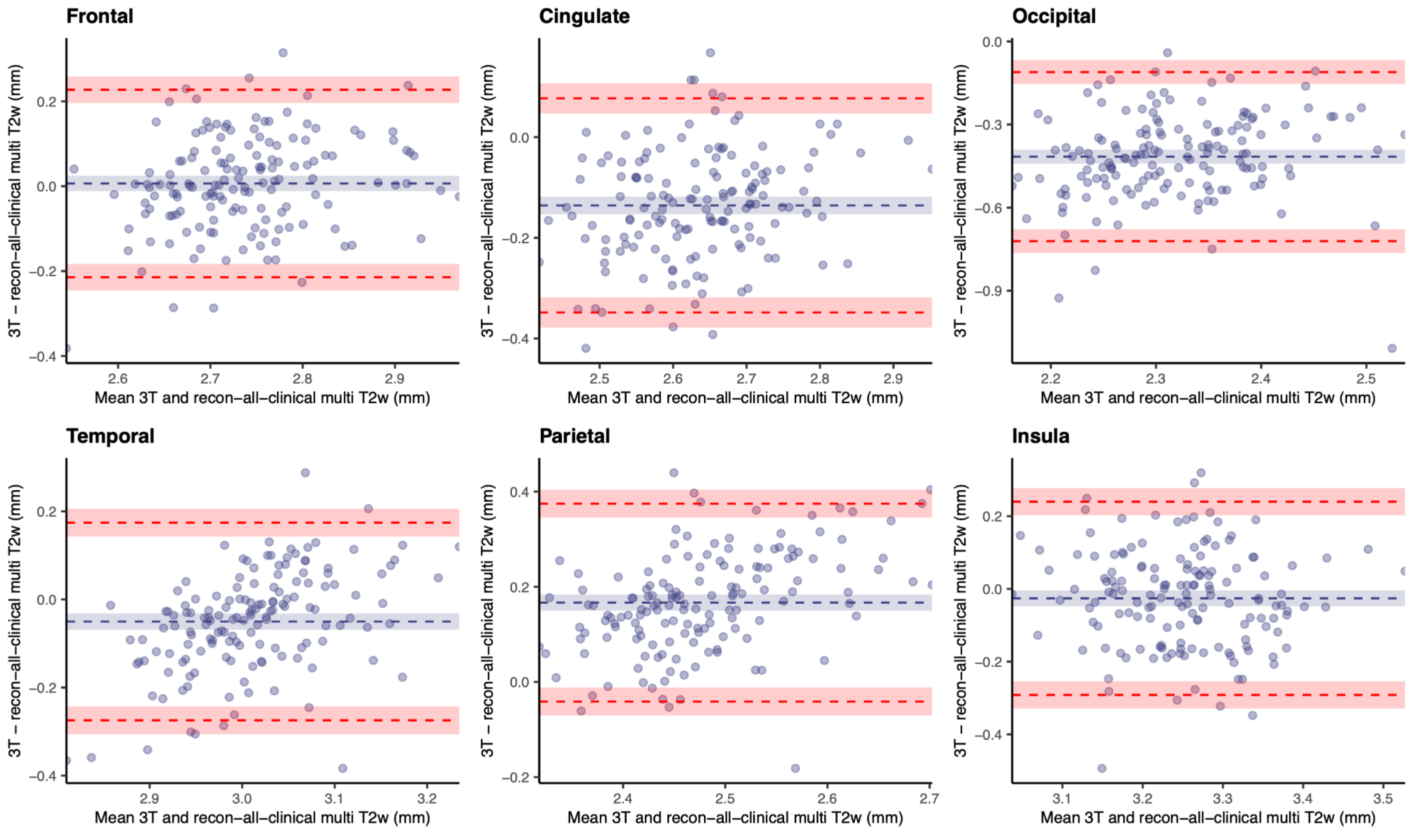


**Figure S11. Bland–Altman plots of agreement between lobar cortical thickness estimates derived from 3T and low-field MRI processed using recon-all-clinical with multi T2w images.** Plots show results for the frontal, cingulate, occipital, temporal, parietal, and insular lobes. Each point represents one participant. The blue dashed line indicates the mean bias, and the upper and lower red dashed lines indicate the 95% limits of agreement. Shaded regions indicate the corresponding 95% confidence intervals. Differences were calculated as 3T minus low-field estimates; therefore, negative values indicate higher cortical thickness estimates from low-field MRI than from 3T MRI.

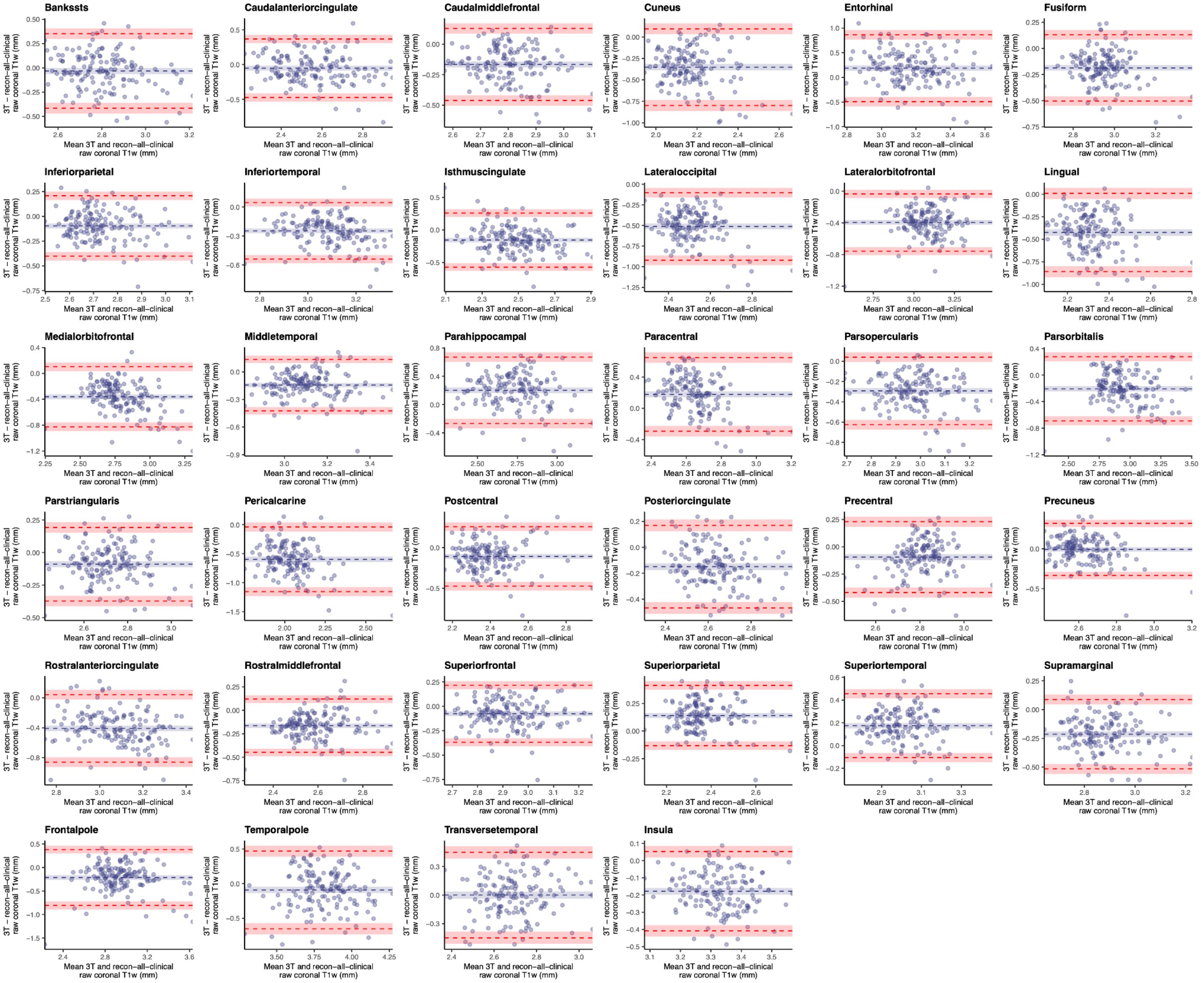


**Figure S12. Bland–Altman plots of agreement between regional cortical thickness estimates derived from 3T and low-field MRI processed using recon-all-clinical with raw coronal T1w images.** Plots show results across cortical regions defined using the Desikan–Killiany atlas. Each point represents one participant. The blue dashed line indicates the mean bias, and the upper and lower red dashed lines indicate the 95% limits of agreement. Shaded regions indicate the corresponding 95% confidence intervals. Differences were calculated as 3T minus low-field estimates; therefore, negative values indicate higher cortical thickness estimates from low-field MRI than from 3T MRI.

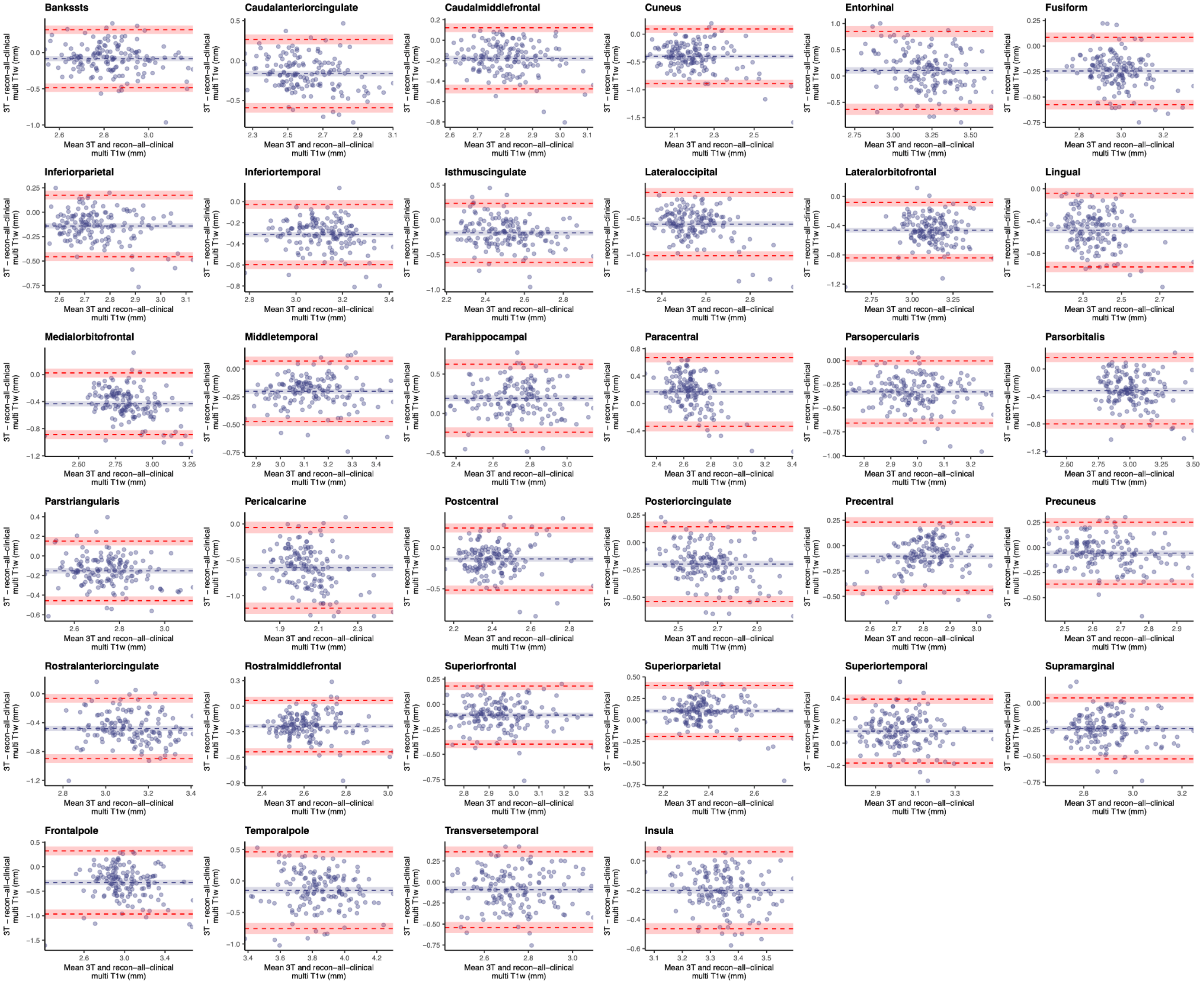


**Figure S13. Bland–Altman plots of agreement between regional cortical thickness estimates derived from 3T and low-field MRI processed using recon-all-clinical with multi T1w images.** Plots show results across cortical regions defined using the Desikan–Killiany atlas. Each point represents one participant. The blue dashed line indicates the mean bias, and the upper and lower red dashed lines indicate the 95% limits of agreement. Shaded regions indicate the corresponding 95% confidence intervals. Differences were calculated as 3T minus low-field estimates; therefore, negative values indicate higher cortical thickness estimates from low-field MRI than from 3T MRI.

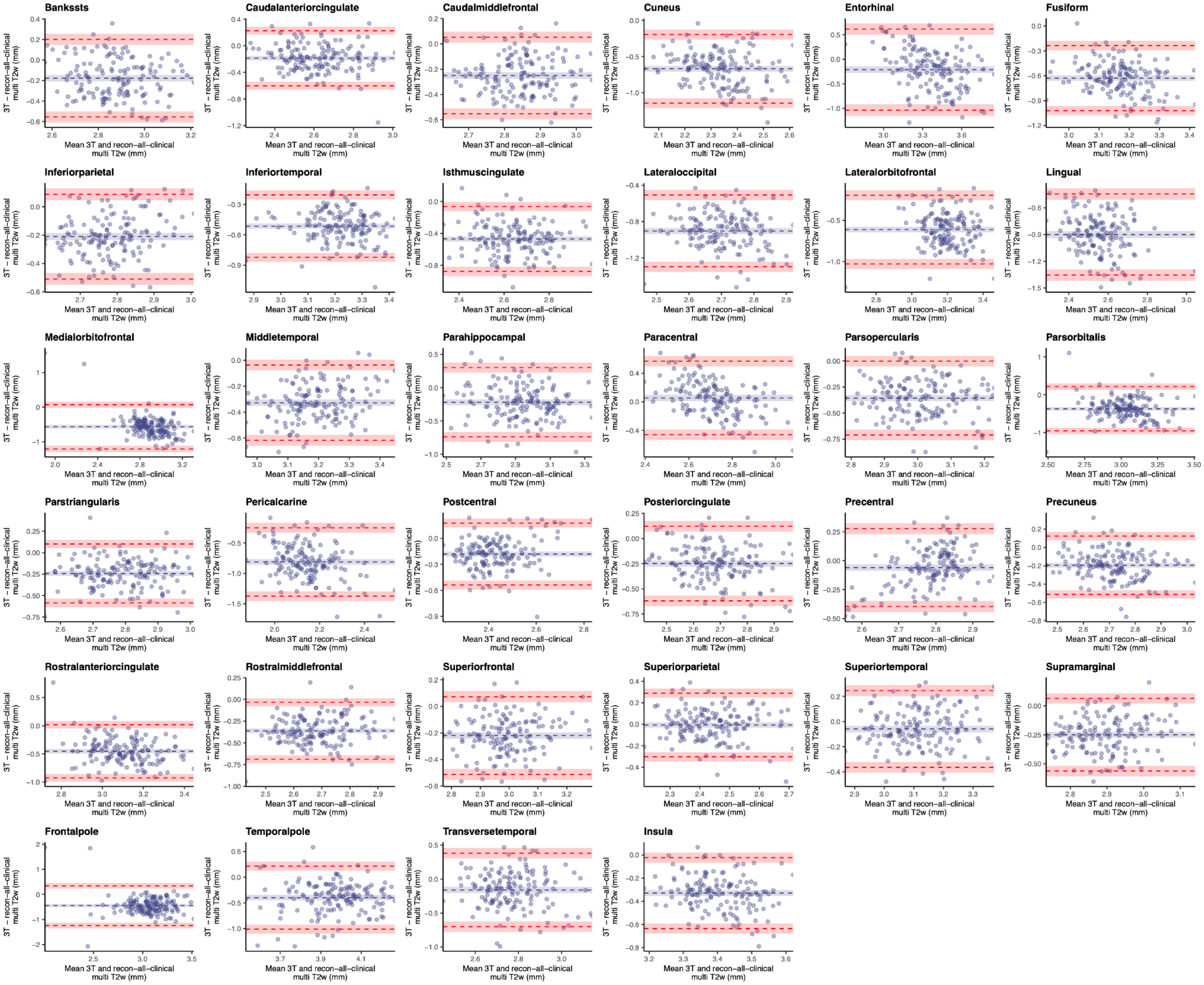


**Figure S14. Bland–Altman plots of agreement between regional cortical thickness estimates derived from 3T and low-field MRI processed using recon-all-clinical with multi T2w images.** Plots show results across cortical regions defined using the Desikan–Killiany atlas. Each point represents one participant. The blue dashed line indicates the mean bias, and the upper and lower red dashed lines indicate the 95% limits of agreement. Shaded regions indicate the corresponding 95% confidence intervals. Differences were calculated as 3T minus low-field estimates; therefore, negative values indicate higher cortical thickness estimates from low-field MRI than from 3T MRI.

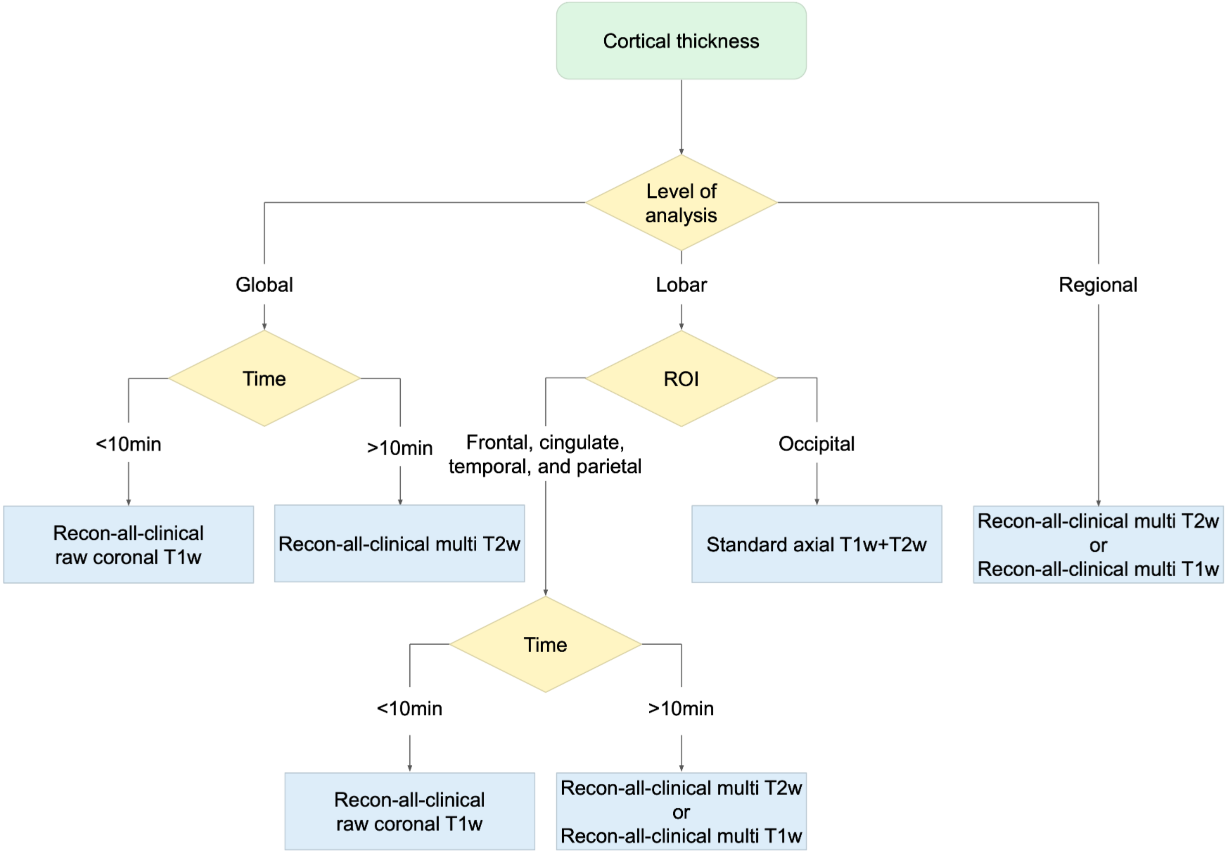


**Figure S15. Decision flowchart for selecting low-field MRI processing pipelines for cortical thickness estimation.** The flowchart summarizes recommended processing pipelines according to the level of analysis, ROI, and available acquisition time. Where two pipelines are shown, either may be considered. For detailed lobe- and region-specific best-performing pipelines, see Figure 4. ROI, region of interest.

**Table S1. Scan parameters for the 64mT and 3T scans acquired in this study**

| Scan Type | Scanner Manufacturer | Scanner Model | Repetition Time (s) | Echo Time (s) | Inversion Time (s) | Flip Angle (deg) | Percent Phase FOV | Slice Thickness (mm) | Slice Resolution (mm) | Total Acquisition Time (min:s) |
|---|---|---|---|---|---|---|---|---|---|---|
| 64mT Axial T1w | Hyperfine | Swoop | 1.5 | 0.00596 | 0.3 | 90 | 100.00 | 5.0 | 1.6 x 1.6 | 5:39 |
| 64mT Coronal T1w | Hyperfine | Swoop | 1.5 | 0.00559 | 0.3 | 90 | 100.00 | 5.0 | 1.6 x 1.6 | 5:32 |
| 64mT Sagittal T1w | Hyperfine | Swoop | 1.5 | 0.00588 | 0.3 | 90 | 100.00 | 5.0 | 1.6 x 1.6 | 5:39 |
| 3T T1w | Siemens | Prisma | 2.5 | 0.00207 | 1 | 8 | 93.75 | 0.8 | 0.8 x 0.8 | 6:34 |
| 64mT Axial T2w | Hyperfine | Swoop | 2.0 | 0.18240 | • | 90 | 100.00 | 5.0 | 1.6 x 1.6 | 2:53 |
| 64mT Coronal T2w | Hyperfine | Swoop | 2.0 | 0.21760 | • | 90 | 65.00 | 5.0 | 1.6 x 1.6 | 2:21 |
| 64mT Sagittal T2w | Hyperfine | Swoop | 2.0 | 0.23000 | • | 90 | 65.00 | 5.0 | 1.6 x 1.6 | 1:59 |
| 3T T2w | Siemens | Prisma | 3.2 | 0.56400 | • | 120 | 93.75 | 0.8 | 0.8 x 0.8 | 5:57 |

Note: Four participants were scanned using a slightly modified T1-weighted 64mT protocol (repetition time = 0.88 s; inversion time = 0.354 s). Excluding these participants from analyses did not significantly alter the results.

**Table S2. Description of processing approaches**

| Tools | Published / released year | Processing approach | Key improvements |
|---|---|---|---|
| SynthSR v1 (+ recon-all) | 2022 | Original version of the deep learning-based super-resolution method for transforming LF-MRI scans into higher-resolution images. In this process, LF-MRI scans are input into a pretrained CNN to generate synthetic HF images. T1w and T2w images acquired with the standard Hyperfine sequence (1.5 × 1.5 × 1.5 mm axial) are used to generate a 1-mm isotropic T1w image, which meets the prerequisites for existing neuroimaging analysis tools such as recon-all. We reported results using this method in our previous manuscript (Cooper et al., 2024). | Performs super-resolution using a deep learning–based image synthesis approach. The resulting synthetic HF outputs meet the resolution and contrast prerequisites required by existing neuroimaging analysis tools and can be used for downstream structural analysis. |
| SynthSR v2 (+ recon-all) | 2023 | Updated super-resolution method based on the original SynthSR. This tool also transforms low-resolution clinical MRI scans into synthetic HF images via a pretrained CNN, with improved generalizability, more realistic synthesis, and better handling of brain abnormalities. A single input scan of any contrast, resolution, or orientation can be used as input to generate a 1-mm isotropic T1w image, which is directly usable by existing neuroimaging analysis tools. | (1) Improves generalizability across input scans by using a domain randomization strategy that generates synthetic training data with fully randomized acquisition parameters; (2) Produces more realistic images by incorporating a pretrained auxiliary segmentation CNN into the model architecture; (3) Handles abnormalities by inpainting them with normal-looking tissue, enabling structural analysis with existing neuroimaging tools that typically cannot cope with various forms of pathology (particularly large lesions). |
| Recon-all-clinical | 2023 | Deep learning–based analysis pipeline for clinical MRI scans. This tool employs CNN-based approaches for brain segmentation and cortical surface reconstruction—tasks that are traditionally performed in neuroimaging analysis pipelines. It can analyze low-resolution images faster, without requiring a separate super-resolution step, and helps prevent the over-smoothing that can occur during super-resolution and conventional surface reconstruction. Clinical MRI scans of any contrast, resolution, or orientation can be used as input, although the pipeline is optimized for HF scans. | SynthSeg brain segmentation method and signed distance function-based cortical surface reconstruction method have been integrated into an existing geometry processing pipeline. Both components are based on CNN models trained with domain randomization. This approach can process heterogeneous clinical MRI scans faster, without requiring super-resolution, and enables more accurate surface reconstruction. (SynthSR is performed within the pipeline, but its output is used only for visualization and is not incorporated into subsequent processing steps.) |
| Recon-any | 2024 | Based on the recon-all-clinical approach, recon-any was trained on LF MRI data. This tool also employs CNN-based approaches for brain segmentation and cortical surface reconstruction as part of the neuroimaging analysis pipeline. LF MRI scans with arbitrary contrast, resolution, or orientation can be used for structural analysis with this tool. | (1) Trained to generalize across heterogeneous LF-MRI acquisitions; (2) Validated on postmortem LF-MRI data |

LF, low-field; HF, high-field; CNN, convolutional neural network.

**Table S3. Low-field pipelines tested in this study**

| | Low-field pipeline name | Super-resolution method | Reconstruction and segmentation pipeline | Preprocessing method | Orientation | Pulse sequence |
|---|---|---|---|---|---|---|
| 1 | Standard_axial_T1w+T2w | None | Recon-all | resampled+co-registered | axial | T1w+T2w |
| 2 | Standard_coronal_T1w+T2w | None | Recon-all | resampled+co-registered | coronal | T1w+T2w |
| 3 | Standard_sagittal_T1w+T2w | None | Recon-all | resampled+co-registered | sagittal | T1w+T2w |
| 4 | Standard_multi_T1w+T2w | None | Recon-all | resampled+co-registered | multi | T1w+T2w |
| 5 | SynthSRv1_axial_T1w+T2w | SynthSRv1 | Recon-all | resampled+co-registered | axial | T1w+T2w |
| 6 | SynthSRv1_coronal_T1w+T2w | SynthSRv1 | Recon-all | resampled+co-registered | coronal | T1w+T2w |
| 7 | SynthSRv1_sagittal_T1w+T2w | SynthSRv1 | Recon-all | resampled+co-registered | sagittal | T1w+T2w |
| 8 | SynthSRv1_multi_T1w+T2w | SynthSRv1 | Recon-all | resampled+co-registered | multi | T1w+T2w |
| 9 | SynthSRv2_resampled_axial_T1w | SynthSRv2 | Recon-all | resampled | axial | T1w |
| 10 | SynthSRv2_resampled_coronal_T1w | SynthSRv2 | Recon-all | resampled | coronal | T1w |
| 11 | SynthSRv2_resampled_sagittal_T1w | SynthSRv2 | Recon-all | resampled | sagittal | T1w |
| 12 | SynthSRv2_multi_T1w | SynthSRv2 | Recon-all | resampled | multi | T1w |
| 13 | SynthSRv2_raw_axial_T1w | SynthSRv2 | Recon-all | raw | axial | T1w |
| 14 | SynthSRv2_raw_coronal_T1w | SynthSRv2 | Recon-all | raw | coronal | T1w |
| 15 | SynthSRv2_raw_sagittal_T1w | SynthSRv2 | Recon-all | raw | sagittal | T1w |
| 16 | SynthSRv2_resampled+co-registered_axial_T2w | SynthSRv2 | Recon-all | resampled+co-registered | axial | T2w |
| 17 | SynthSRv2_resampled+co-registered_coronal_T2w | SynthSRv2 | Recon-all | resampled+co-registered | coronal | T2w |
| 18 | SynthSRv2_resampled+co-registered_sagittal_T2w | SynthSRv2 | Recon-all | resampled+co-registered | sagittal | T2w |
| 19 | SynthSRv2_multi_T2w | SynthSRv2 | Recon-all | resampled+co-registered | multi | T2w |
| 20 | SynthSRv2_co-registered_axial_T2w | SynthSRv2 | Recon-all | co-registered | axial | T2w |
| 21 | SynthSRv2_co-registered_coronal_T2w | SynthSRv2 | Recon-all | co-registered | coronal | T2w |
| 22 | SynthSRv2_co-registered_sagittal_T2w | SynthSRv2 | Recon-all | co-registered | sagittal | T2w |
| 23 | SynthSRv2_resampled_axial_T2w | SynthSRv2 | Recon-all | resampled | axial | T2w |
| 24 | SynthSRv2_resampled_coronal_T2w | SynthSRv2 | Recon-all | resampled | coronal | T2w |
| 25 | SynthSRv2_resampled_sagittal_T2w | SynthSRv2 | Recon-all | resampled | sagittal | T2w |
| 26 | SynthSRv2_raw_axial_T2w | SynthSRv2 | Recon-all | raw | axial | T2w |
| 27 | SynthSRv2_raw_coronal_T2w | SynthSRv2 | Recon-all | raw | coronal | T2w |
| 28 | SynthSRv2_raw_sagittal_T2w | SynthSRv2 | Recon-all | raw | sagittal | T2w |
| 29 | Recon-all-clinical_resampled_axial_T1w | None | Recon-all-clinical | resampled | axial | T1w |
| 30 | Recon-all-clinical_resampled_coronal_T1w | None | Recon-all-clinical | resampled | coronal | T1w |
| 31 | Recon-all-clinical_resampled_sagittal_T1w | None | Recon-all-clinical | resampled | sagittal | T1w |
| 32 | Recon-all-clinical_multi_T1w | None | Recon-all-clinical | resampled | multi | T1w |

| | | | | | | |
|---|---|---|---|---|---|---|
| 33 | Recon-all-clinical_raw_axial_T1w | None | Recon-all-clinical | raw | axial | T1w |
| 34 | Recon-all-clinical_raw_coronal_T1w | None | Recon-all-clinical | raw | coronal | T1w |
| 35 | Recon-all-clinical_raw_sagittal_T1w | None | Recon-all-clinical | raw | sagittal | T1w |
| 36 | Recon-all-clinical_resampled+co-registered_axial_T2w | None | Recon-all-clinical | resampled+co-registered | axial | T2w |
| 37 | Recon-all-clinical_resampled+co-registered_coronal_T2w | None | Recon-all-clinical | resampled+co-registered | coronal | T2w |
| 38 | Recon-all-clinical_resampled+co-registered_sagittal_T2w | None | Recon-all-clinical | resampled+co-registered | sagittal | T2w |
| 39 | Recon-all-clinical_multi_T2w | None | Recon-all-clinical | resampled+co-registered | multi | T2w |
| 40 | Recon-all-clinical_co-registered_axial_T2w | None | Recon-all-clinical | co-registered | axial | T2w |
| 41 | Recon-all-clinical_co-registered_coronal_T2w | None | Recon-all-clinical | co-registered | coronal | T2w |
| 42 | Recon-all-clinical_co-registered_sagittal_T2w | None | Recon-all-clinical | co-registered | sagittal | T2w |
| 43 | Recon-all-clinical_resampled_axial_T2w | None | Recon-all-clinical | resampled | axial | T2w |
| 44 | Recon-all-clinical_resampled_coronal_T2w | None | Recon-all-clinical | resampled | coronal | T2w |
| 45 | Recon-all-clinical_resampled_sagittal_T2w | None | Recon-all-clinical | resampled | sagittal | T2w |
| 46 | Recon-all-clinical_raw_axial_T2w | None | Recon-all-clinical | raw | axial | T2w |
| 47 | Recon-all-clinical_raw_coronal_T2w | None | Recon-all-clinical | raw | coronal | T2w |
| 48 | Recon-all-clinical_raw_sagittal_T2w | None | Recon-all-clinical | raw | sagittal | T2w |
| 49 | Recon-any_resampled_axial_T1w | None | Recon-any | resampled | axial | T1w |
| 50 | Recon-any_resampled_coronal_T1w | None | Recon-any | resampled | coronal | T1w |
| 51 | Recon-any_resampled_sagittal_T1w | None | Recon-any | resampled | sagittal | T1w |
| 52 | Recon-any_multi_T1w | None | Recon-any | resampled | multi | T1w |
| 53 | Recon-any_raw_axial_T1w | None | Recon-any | raw | axial | T1w |
| 54 | Recon-any_raw_coronal_T1w | None | Recon-any | raw | coronal | T1w |
| 55 | Recon-any_raw_sagittal_T1w | None | Recon-any | raw | sagittal | T1w |
| 56 | Recon-any_resampled+co-registered_axial_T2w | None | Recon-any | resampled+co-registered | axial | T2w |
| 57 | Recon-any_resampled+co-registered_coronal_T2w | None | Recon-any | resampled+co-registered | coronal | T2w |
| 58 | Recon-any_resampled+co-registered_sagittal_T2w | None | Recon-any | resampled+co-registered | sagittal | T2w |
| 59 | Recon-any_multi_T2w | None | Recon-any | resampled+co-registered | multi | T2w |
| 60 | Recon-any_co-registered_axial_T2w | None | Recon-any | co-registered | axial | T2w |
| 61 | Recon-any_co-registered_coronal_T2w | None | Recon-any | co-registered | coronal | T2w |
| 62 | Recon-any_co-registered_sagittal_T2w | None | Recon-any | co-registered | sagittal | T2w |
| 63 | Recon-any_resampled_axial_T2w | None | Recon-any | resampled | axial | T2w |
| 64 | Recon-any_resampled_coronal_T2w | None | Recon-any | resampled | coronal | T2w |
| 65 | Recon-any_resampled_sagittal_T2w | None | Recon-any | resampled | sagittal | T2w |
| 66 | Recon-any_raw_axial_T2w | None | Recon-any | raw | axial | T2w |

| | | | | | | |
|---|---|---|---|---|---|---|
| 67 | Recon-any_raw_coronal_T2w | None | Recon-any | raw | coronal | T2w |
| 68 | Recon-any_raw_sagittal_T2w | None | Recon-any | raw | sagittal | T2w |

All 68 low-field image processing pipelines tested in this study. The first column lists the pipeline names, and the remaining columns describe the imaging sequences and orientations used, along with the preprocessing and processing methods applied in each pipeline.

**Table S4. Proportion of poor-quality images by processing pipeline**

| Low-field pipeline | Poor-quality image (%) |
|---|---|
| SynthSRv1_coronal_T1w+T2w | 0.67 |
| SynthSRv2_resampled_axial_T1w | 0.33 |
| SynthSRv2_resampled_coronal_T1w | 0.33 |
| SynthSRv2_resampled_sagittal_T1w | 1.67 |
| SynthSRv2_resampled+co-registered_axial_T2w | 2.33 |
| SynthSRv2_resampled+co-registered_coronal_T2w | 7.33 |
| SynthSRv2_resampled+co-registered_sagittal_T2w | 8.67 |
| Recon-all-clinical_resampled_axial_T1w | 1.00 |
| Recon-all-clinical_resampled_sagittal_T1w | 3.33 |
| Recon-all-clinical_resampled+co-registered_coronal_T2w | 0.33 |
| Recon-all-clinical_resampled+co-registered_sagittal_T2w | 2.00 |

Processing pipelines with images rated as poor quality (score = 1) and the percentage of poor-quality images within each pipeline.

**Table S5. Low-field pipelines with significant global Pearson correlations with 3T**

| | Low-field pipeline name | Super-resolution method | Reconstruction and segmentation pipeline | Preprocessing method | Orientation | Pulse sequence |
|---|---|---|---|---|---|---|
| 1 | Standard_axial_T1w+T2w | None | Recon-all | resampled+co-registered | axial | T1w+T2w |
| 2 | Standard_coronal_T1w+T2w | None | Recon-all | resampled+co-registered | coronal | T1w+T2w |
| 3 | Standard_sagittal_T1w+T2w | None | Recon-all | resampled+co-registered | sagittal | T1w+T2w |
| 4 | SynthSRv1_axial_T1w+T2w | SynthSRv1 | Recon-all | resampled+co-registered | axial | T1w+T2w |
| 5 | SynthSRv1_coronal_T1w+T2w | SynthSRv1 | Recon-all | resampled+co-registered | coronal | T1w+T2w |
| 6 | SynthSRv1_sagittal_T1w+T2w | SynthSRv1 | Recon-all | resampled+co-registered | sagittal | T1w+T2w |
| 7 | SynthSRv1_multi_T1w+T2w | SynthSRv1 | Recon-all | resampled+co-registered | multi | T1w+T2w |
| 8 | SynthSRv2_resampled_coronal_T1w | SynthSRv2 | Recon-all | resampled | coronal | T1w |
| 9 | SynthSRv2_multi_T1w | SynthSRv2 | Recon-all | resampled | multi | T1w |
| 10 | SynthSRv2_raw_coronal_T1w | SynthSRv2 | Recon-all | raw | coronal | T1w |
| 11 | SynthSRv2_multi_T2w | SynthSRv2 | Recon-all | resampled+co-registered | multi | T2w |
| 12 | Recon-all-clinical_resampled_axial_T1w | None | Recon-all-clinical | resampled | axial | T1w |
| 13 | Recon-all-clinical_resampled_coronal_T1w | None | Recon-all-clinical | resampled | coronal | T1w |
| 14 | Recon-all-clinical_resampled_multi_T1w | None | Recon-all-clinical | resampled | multi | T1w |
| 15 | Recon-all-clinical_raw_axial_T1w | None | Recon-all-clinical | raw | axial | T1w |
| 16 | Recon-all-clinical_raw_coronal_T1w | None | Recon-all-clinical | raw | coronal | T1w |
| 17 | Recon-all-clinical_raw_sagittal_T1w | None | Recon-all-clinical | raw | sagittal | T1w |
| 18 | Recon-all-clinical_resampled+co-registered_axial_T2w | None | Recon-all-clinical | resampled+co-registered | axial | T2w |
| 19 | Recon-all-clinical_resampled+co-registered_coronal_T2w | None | Recon-all-clinical | resampled+co-registered | coronal | T2w |
| 20 | Recon-all-clinical_resampled+co-registered_sagittal_T2w | None | Recon-all-clinical | resampled+co-registered | sagittal | T2w |
| 21 | Recon-all-clinical_resampled+co-registered_multi_T2w | None | Recon-all-clinical | resampled+co-registered | multi | T2w |
| 22 | Recon-all-clinical_co-registered_axial_T2w | None | Recon-all-clinical | co-registered | axial | T2w |
| 23 | Recon-all-clinical_co-registered_coronal_T2w | None | Recon-all-clinical | co-registered | coronal | T2w |
| 24 | Recon-all-clinical_co-registered_sagittal_T2w | None | Recon-all-clinical | co-registered | sagittal | T2w |
| 25 | Recon-all-clinical_resampled_axial_T2w | None | Recon-all-clinical | resampled | axial | T2w |
| 26 | Recon-all-clinical_resampled_coronal_T2w | None | Recon-all-clinical | resampled | coronal | T2w |
| 27 | Recon-all-clinical_raw_axial_T2w | None | Recon-all-clinical | raw | axial | T2w |
| 28 | Recon-all-clinical_raw_coronal_T2w | None | Recon-all-clinical | raw | coronal | T2w |
| 29 | Recon-all-clinical_raw_sagittal_T2w | None | Recon-all-clinical | raw | sagittal | T2w |
| 30 | Recon-any_resampled_coronal_T1w | None | Recon-any | resampled | coronal | T1w |
| 31 | Recon-any_raw_coronal_T1w | None | Recon-any | raw | coronal | T1w |
| 32 | Recon-any_resampled+co-registered_coronal_T2w | None | Recon-any | resampled+co-registered | coronal | T2w |
| 33 | Recon-any_resampled+co-registered_sagittal_T2w | None | Recon-any | resampled+co-registered | sagittal | T2w |
| 34 | Recon-any_resampled_axial_T2w | None | Recon-any | resampled | axial | T2w |

| | | | | | | |
|---|---|---|---|---|---|---|
| 35 | Recon-any_resampled_coronal_T2w | None | Recon-any | resampled | coronal | T2w |
| 36 | Recon-any_resampled_sagittal_T2w | None | Recon-any | resampled | sagittal | T2w |
| 37 | Recon-any_raw_coronal_T2w | None | Recon-any | raw | coronal | T2w |
| 38 | Recon-any_raw_sagittal_T2w | None | Recon-any | raw | sagittal | T2w |

Low-field image processing pipelines showing significant Pearson correlations in global cortical thickness with 3T image ($p_{FDR}$ < 0.05). The first column lists the pipeline names, and the remaining columns describe the imaging sequences and orientations used, along with the preprocessing and processing methods applied in each pipeline.

**Table S6. Low-field pipelines with significant global intraclass correlations with 3T**

| | Low-field pipeline name | Super-resolution method | Reconstruction and segmentation pipeline | Preprocessing method | Orientation | Pulse sequence |
|---|---|---|---|---|---|---|
| 1 | Standard_axial_T1w+T2w | None | Recon-all | resampled+co-registered | axial | T1w+T2w |
| 2 | Standard_coronal_T1w+T2w | None | Recon-all | resampled+co-registered | coronal | T1w+T2w |
| 3 | Standard_sagittal_T1w+T2w | None | Recon-all | resampled+co-registered | sagittal | T1w+T2w |
| 4 | SynthSRv1_axial_T1w+T2w | SynthSRv1 | Recon-all | resampled+co-registered | axial | T1w+T2w |
| 5 | SynthSRv1_coronal_T1w+T2w | SynthSRv1 | Recon-all | resampled+co-registered | coronal | T1w+T2w |
| 6 | SynthSRv1_sagittal_T1w+T2w | SynthSRv1 | Recon-all | resampled+co-registered | sagittal | T1w+T2w |
| 7 | SynthSRv1_multi_T1w+T2w | SynthSRv1 | Recon-all | resampled+co-registered | multi | T1w+T2w |
| 8 | SynthSRv2_resampled_coronal_T1w | SynthSRv2 | Recon-all | resampled | coronal | T1w |
| 9 | SynthSRv2_multi_T1w | SynthSRv2 | Recon-all | resampled | multi | T1w |
| 10 | SynthSRv2_raw_coronal_T1w | SynthSRv2 | Recon-all | raw | coronal | T1w |
| 11 | SynthSRv2_multi_T2w | SynthSRv2 | Recon-all | resampled+co-registered | multi | T2w |
| 12 | Recon-all-clinical_resampled_axial_T1w | None | Recon-all-clinical | resampled | axial | T1w |
| 13 | Recon-all-clinical_resampled_coronal_T1w | None | Recon-all-clinical | resampled | coronal | T1w |
| 14 | Recon-all-clinical_resampled_multi_T1w | None | Recon-all-clinical | resampled | multi | T1w |
| 15 | Recon-all-clinical_raw_axial_T1w | None | Recon-all-clinical | raw | axial | T1w |
| 16 | Recon-all-clinical_raw_coronal_T1w | None | Recon-all-clinical | raw | coronal | T1w |
| 17 | Recon-all-clinical_raw_sagittal_T1w | None | Recon-all-clinical | raw | sagittal | T1w |
| 18 | Recon-all-clinical_resampled+co-registered_axial_T2w | None | Recon-all-clinical | resampled+co-registered | axial | T2w |
| 19 | Recon-all-clinical_resampled+co-registered_coronal_T2w | None | Recon-all-clinical | resampled+co-registered | coronal | T2w |
| 20 | Recon-all-clinical_resampled+co-registered_sagittal_T2w | None | Recon-all-clinical | resampled+co-registered | sagittal | T2w |
| 21 | Recon-all-clinical_resampled+co-registered_multi_T2w | None | Recon-all-clinical | resampled+co-registered | multi | T2w |
| 22 | Recon-all-clinical_co-registered_axial_T2w | None | Recon-all-clinical | co-registered | axial | T2w |
| 23 | Recon-all-clinical_co-registered_coronal_T2w | None | Recon-all-clinical | co-registered | coronal | T2w |
| 24 | Recon-all-clinical_co-registered_sagittal_T2w | None | Recon-all-clinical | co-registered | sagittal | T2w |
| 25 | Recon-all-clinical_resampled_axial_T2w | None | Recon-all-clinical | resampled | axial | T2w |
| 26 | Recon-all-clinical_resampled_coronal_T2w | None | Recon-all-clinical | resampled | coronal | T2w |
| 27 | Recon-all-clinical_raw_axial_T2w | None | Recon-all-clinical | raw | axial | T2w |
| 28 | Recon-all-clinical_raw_coronal_T2w | None | Recon-all-clinical | raw | coronal | T2w |
| 29 | Recon-all-clinical_raw_sagittal_T2w | None | Recon-all-clinical | raw | sagittal | T2w |
| 30 | Recon-any_resampled_coronal_T1w | None | Recon-any | resampled | coronal | T1w |
| 31 | Recon-any_raw_coronal_T1w | None | Recon-any | raw | coronal | T1w |
| 32 | Recon-any_resampled+co-registered_axial_T2w | None | Recon-any | resampled+co-registered | axial | T2w |
| 33 | Recon-any_resampled+co-registered_coronal_T2w | None | Recon-any | resampled+co-registered | coronal | T2w |

| | | | | | | |
|---|---|---|---|---|---|---|
| 34 | Recon-any_resampled+co-registered_sagittal_T2w | None | Recon-any | resampled+co-registered | sagittal | T2w |
| 35 | Recon-any_co-registered_coronal_T2w | None | Recon-any | co-registered | coronal | T2w |
| 36 | Recon-any_resampled_axial_T2w | None | Recon-any | resampled | axial | T2w |
| 37 | Recon-any_resampled_coronal_T2w | None | Recon-any | resampled | coronal | T2w |
| 38 | Recon-any_resampled_sagittal_T2w | None | Recon-any | resampled | sagittal | T2w |
| 39 | Recon-any_raw_coronal_T2w | None | Recon-any | raw | coronal | T2w |
| 40 | Recon-any_raw_sagittal_T2w | None | Recon-any | raw | sagittal | T2w |

Low-field image processing pipelines showing significant intraclass correlations in global cortical thickness with 3T image ($p_{FDR} < 0.05$). The first column lists the pipeline names, and the remaining columns describe the imaging sequences and orientations used, along with the preprocessing and processing methods applied in each pipeline.

**Table S7. Intraclass correlation coefficients (ICCs) for the global correspondence between low-field and 3T images**

| | Standard T1 and T2 | | | SynthSR v1 T1 and T2 | | |
|---|---|---|---|---|---|---|
| | ICC | $p$ | $p_{FDR}$ | ICC | $p$ | $p_{FDR}$ |
| axi | **0.24** | **0.001** | **0.003** | **0.28** | **2.47e-04** | **0.002** |
| cor | **0.23** | **0.002** | **0.005** | **0.33** | **2.01e-05** | **2.61e-04** |
| sag | **0.27** | **4.29e-04** | **0.002** | **0.19** | **0.011** | **0.020** |
| multi | -0.06 | 0.768 | 0.792 | **0.22** | **0.003** | **0.007** |

| | SynthSR v2 resampled T1 | | | SynthSR v2 raw T1 | | | SynthSR v2 resampled and co-registered T2 | | | SynthSR v2 co-registered T2 | | | SynthSR v2 resampled T2 | | | SynthSR v2 raw T2 | | |
|---|---|---|---|---|---|---|---|---|---|---|---|---|---|---|---|---|---|---|
| | ICC | $p$ | $p_{FDR}$ | ICC | $p$ | $p_{FDR}$ | ICC | $p$ | $p_{FDR}$ | ICC | $p$ | $p_{FDR}$ | ICC | $p$ | $p_{FDR}$ | ICC | $p$ | $p_{FDR}$ |
| axi | 0.10 | 0.118 | 0.139 | 0.12 | 0.077 | 0.106 | 0.10 | 0.113 | 0.139 | 0.13 | 0.059 | 0.087 | 0.11 | 0.083 | 0.112 | 0.13 | 0.062 | 0.090 |
| cor | **0.19** | **0.010** | **0.019** | **0.32** | **3.50e-05** | **3.18e-04** | -0.15 | 0.965 | 0.965 | 0.13 | 0.055 | 0.085 | -0.09 | 0.862 | 0.875 | 0.10 | 0.118 | 0.139 |
| sag | 0.11 | 0.093 | 0.121 | 0.10 | 0.114 | 0.139 | — | — | — | 0.10 | 0.108 | 0.138 | — | — | — | — | — | — |
| multi | **0.18** | **0.014** | **0.024** | N/A | N/A | N/A | **0.20** | **0.006** | **0.013** | N/A | N/A | N/A | N/A | N/A | N/A | N/A | N/A | N/A |

| | Recon-all-clinical resampled T1 | | | Recon-all-clinical raw T1 | | | Recon-all-clinical resampled and co-registered T2 | | | Recon-all-clinical co-registered T2 | | | Recon-all-clinical resampled T2 | | | Recon-all-clinical raw T2 | | |
|---|---|---|---|---|---|---|---|---|---|---|---|---|---|---|---|---|---|---|
| | ICC | $p$ | $p_{FDR}$ | ICC | $p$ | $p_{FDR}$ | ICC | $p$ | $p_{FDR}$ | ICC | $p$ | $p_{FDR}$ | ICC | $p$ | $p_{FDR}$ | ICC | $p$ | $p_{FDR}$ |
| axi | **0.30** | **7.30e-05** | **5.27e-04** | **0.32** | **3.91e-05** | **3.18e-04** | **0.26** | **5.22e-04** | **0.002** | **0.32** | **2.74e-05** | **2.97e-04** | **0.26** | **6.53e-04** | **0.003** | **0.28** | **2.78e-04** | **0.002** |
| cor | **0.36** | **2.16e-06** | **6.39e-05** | **0.40** | **2.29e-07** | **1.49e-05** | **0.20** | **0.008** | **0.016** | **0.26** | **7.74e-04** | **0.003** | **0.18** | **0.014** | **0.024** | **0.22** | **0.003** | **0.007** |
| sag | 0.12 | 0.072 | 0.102 | **0.20** | **0.007** | **0.014** | **0.21** | **0.005** | **0.011** | **0.21** | **0.004** | **0.009** | 0.01 | 0.470 | 0.493 | **0.23** | **0.002** | **0.005** |
| multi | **0.33** | **1.99e-05** | **2.61e-04** | N/A | N/A | N/A | **0.36** | **2.95e-06** | **6.39e-05** | N/A | N/A | N/A | N/A | N/A | N/A | N/A | N/A | N/A |

| | Recon-any resampled T1 | | | Recon-any raw T1 | | | Recon-any resampled and co-registered T2 | | | Recon-any co-registered T2 | | | Recon-any resampled T2 | | | Recon-any raw T2 | | |
|---|---|---|---|---|---|---|---|---|---|---|---|---|---|---|---|---|---|---|
| | ICC | $p$ | $p_{FDR}$ | ICC | $p$ | $p_{FDR}$ | ICC | $p$ | $p_{FDR}$ | ICC | $p$ | $p_{FDR}$ | ICC | $p$ | $p_{FDR}$ | ICC | $p$ | $p_{FDR}$ |
| axi | 0.08 | 0.175 | 0.197 | 0.06 | 0.236 | 0.251 | **0.16** | **0.028** | **0.045** | 0.08 | 0.167 | 0.194 | **0.25** | **0.001** | **0.003** | 0.13 | 0.059 | 0.087 |
| cor | **0.20** | **0.007** | **0.014** | **0.19** | **0.010** | **0.019** | **0.23** | **0.003** | **0.007** | **0.18** | **0.016** | **0.027** | **0.27** | **4.83e-04** | **0.002** | **0.24** | **0.001** | **0.003** |
| sag | 0.07 | 0.211 | 0.232 | 0.08 | 0.176 | 0.197 | **0.24** | **0.002** | **0.005** | 0.11 | 0.088 | 0.117 | **0.24** | **0.001** | **0.003** | **0.23** | **0.002** | **0.005** |
| multi | 0.14 | 0.045 | 0.071 | N/A | N/A | N/A | 0.06 | 0.228 | 0.247 | N/A | N/A | N/A | N/A | N/A | N/A | N/A | N/A | N/A |

Global mean cortical thickness correspondence between 3T and low-field images across different pipelines. N/A indicates pipeline combinations that were not applicable because resampling and co-registration are required to generate multi-orientation images. An em dash (—) indicates pipelines for which cortical thickness estimation failed for some participants. Bold values survived multiple testing corrections at $p_{FDR} < 0.05$.

**Table S8. Comparison of low-field pipeline group–average Pearson correlation coefficients**

| Pipeline 1 | Pipeline 2 | *r*1 | *r*2 | Fisher's *Z* | *p* | $p_{FDR}$ |
|---|---|---|---|---|---|---|
| Recon-all-clinical_T1w | Standard_T1w+T2w | 0.29 | 0.17 | **2.46** | **0.014** | **0.026** |
| Recon-all-clinical_T1w | SynthSRv1_T1w+T2w | 0.29 | 0.27 | 0.42 | 0.672 | 0.672 |
| Recon-all-clinical_T1w | SynthSRv2_T1w | 0.29 | 0.17 | **3.04** | **0.002** | **0.006** |
| Recon-all-clinical_T1w | SynthSRv2_T2w | 0.29 | 0.10 | **5.13** | **2.87e-07** | **3.73e-06** |
| Recon-all-clinical_T1w | Recon-all-clinical_T2w | 0.29 | 0.24 | 1.49 | 0.165 | 0.195 |
| Recon-all-clinical_T1w | Recon-any_T1w | 0.29 | 0.12 | **4.22** | **2.40e-05** | **1.04e-04** |
| Recon-all-clinical_T1w | Recon-any_T2w | 0.29 | 0.19 | **2.79** | **0.005** | **0.011** |

Pipeline 1 is the pipeline group showing the highest average correspondence with 3T, whereas Pipeline 2 represents each of the remaining seven pipeline groups. *r*1 and *r*2 denote the average Pearson correlation coefficients for Pipeline 1 and Pipeline 2, respectively. *Z*, *p*, and FDR-corrected *p* values ($p_{FDR}$) were obtained from Fisher's Z tests comparing Pipeline 1 with each Pipeline 2 group. Bold values survived multiple-testing correction at $p_{FDR} < 0.05$.

**Table S9. Comparison of low-field pipeline group–average intraclass correlation coefficients (ICCs)**

| Pipeline 1 | Pipeline 2 | ICC1 | ICC2 | Fisher's *Z* | *p* | $p_{FDR}$ |
|---|---|---|---|---|---|---|
| Recon-all-clinical_T1w | Standard_T1w+T2w | 0.29 | 0.17 | **2.48** | **0.013** | **0.024** |
| Recon-all-clinical_T1w | SynthSRv1_T1w+T2w | 0.29 | 0.26 | 0.77 | 0.441 | 0.478 |
| Recon-all-clinical_T1w | SynthSRv2_T1w | 0.29 | 0.16 | **3.17** | **0.002** | **0.005** |
| Recon-all-clinical_T1w | SynthSRv2_T2w | 0.29 | 0.08 | **5.58** | **2.47e-08** | **3.21e-07** |
| Recon-all-clinical_T1w | Recon-all-clinical_T2w | 0.29 | 0.23 | 1.70 | 0.090 | 0.130 |
| Recon-all-clinical_T1w | Recon-any_T1w | 0.29 | 0.12 | **4.19** | **2.81e-05** | **1.22e-04** |
| Recon-all-clinical_T1w | Recon-any_T2w | 0.29 | 0.19 | **2.92** | **0.004** | **0.008** |

Pipeline 1 is the pipeline group showing the highest average correspondence with 3T, whereas Pipeline 2 represents each of the remaining seven pipeline groups. ICC1 and ICC2 denote the average ICC for Pipeline 1 and Pipeline 2, respectively. *Z*, *p*, and FDR-corrected *p* values ($p_{FDR}$) were obtained from Fisher's Z tests comparing Pipeline 1 with each Pipeline 2 group. Bold values survived multiple-testing correction at $p_{FDR} < 0.05$.

**Table S10. Pearson correlations between processed low-field and 3T global measures using only good-quality images**

| Low-field pipeline | *r* | *p* | $p_{FDR}$ | N Subjects |
|---|---|---|---|---|
| SynthSRv1_axial_T1w+T2w | **0.34** | **2.80e-05** | **1.29e-04** | 144 |
| SynthSRv1_coronal_T1w+T2w | **0.35** | **1.67e-05** | **1.28e-04** | 145 |
| SynthSRv1_sagittal_T1w+T2w | **0.19** | **0.021** | **0.030** | 148 |
| SynthSRv1_multi_T1w+T2w | **0.23** | **0.005** | **0.013** | 149 |
| SynthSRv2_resampled_axial_T1w | 0.13 | 0.113 | 0.130 | 140 |
| SynthSRv2_resampled_coronal_T1w | **0.22** | **0.009** | **0.018** | 146 |
| SynthSRv2_resampled_sagittal_T1w | 0.14 | 0.113 | 0.130 | 136 |
| SynthSRv2_multi_T1w | **0.18** | **0.026** | **0.035** | 149 |
| SynthSRv2_resampled+co-registered_axial_T2w | **0.25** | **0.010** | **0.018** | 107 |
| SynthSRv2_resampled+co-registered_coronal_T2w | -0.15 | 0.186 | 0.204 | 83 |
| SynthSRv2_resampled+co-registered_sagittal_T2w | 0.14 | 0.259 | 0.271 | 70 |
| Recon-all-clinical_resampled_axial_T1w | **0.31** | **1.71e-04** | **5.62e-04** | 143 |
| Recon-all-clinical_resampled_coronal_T1w | **0.37** | **3.30e-06** | **3.80e-05** | 147 |
| Recon-all-clinical_resampled_sagittal_T1w | **0.32** | **1.06e-04** | **4.06e-04** | 142 |
| Recon-all-clinical_resampled+co-registered_axial_T2w | **0.30** | **2.78e-04** | **7.99e-04** | 143 |
| Recon-all-clinical_resampled+co-registered_coronal_T2w | **0.20** | **0.017** | **0.026** | 140 |
| Recon-all-clinical_resampled+co-registered_sagittal_T2w | **0.35** | **2.78e-05** | **1.29e-04** | 140 |
| Recon-all-clinical_multi_T2w | **0.38** | **1.70e-06** | **3.80e-05** | 149 |
| Recon-any_resampled_axial_T1w | 0.07 | 0.420 | 0.420 | 146 |
| Recon-any_resampled_coronal_T1w | **0.20** | **0.014** | **0.023** | 150 |
| Recon-any_resampled+co-registered_axial_T2w | **0.18** | **0.031** | **0.040** | 143 |
| Recon-any_resampled+co-registered_coronal_T2w | **0.22** | **0.007** | **0.016** | 146 |
| Recon-any_resampled+co-registered_sagittal_T2w | **0.21** | **0.010** | **0.018** | 146 |

Pearson correlation coefficients between processed low-field and 3T global mean cortical thickness, computed using only images rated as good quality (score=3). Images rated as poor (score=1) or fair (score=2) were excluded from the analysis. The number of subjects remaining after exclusion is reported. Pipelines in which all images were rated as good quality were not included in the table. Results significant at $p_{FDR} < 0.05$ after multiple-comparison correction are shown in bold.

**Table S11. Intraclass correlations between processed low-field and 3T global measures using only good-quality images**

| Low-field pipeline | ICC | *p* | $p_{FDR}$ | N Subjects |
|---|---|---|---|---|
| SynthSRv1_axial_T1w+T2w | **0.30** | **1.03e-04** | **3.61e-04** | 144 |
| SynthSRv1_coronal_T1w+T2w | **0.32** | **3.72e-05** | **2.14e-04** | 145 |
| SynthSRv1_sagittal_T1w+T2w | **0.18** | **0.012** | **0.018** | 148 |
| SynthSRv1_multi_T1w+T2w | **0.23** | **0.003** | **0.008** | 149 |
| SynthSRv2_resampled_axial_T1w | 0.12 | 0.081 | 0.098 | 140 |
| SynthSRv2_resampled_coronal_T1w | **0.20** | **0.006** | **0.013** | 146 |
| SynthSRv2_resampled_sagittal_T1w | 0.12 | 0.088 | 0.101 | 136 |
| SynthSRv2_multi_T1w | **0.18** | **0.014** | **0.020** | 149 |
| SynthSRv2_resampled+co-registered_axial_T2w | **0.18** | **0.034** | **0.043** | 107 |
| SynthSRv2_resampled+co-registered_coronal_T2w | -0.11 | 0.835 | 0.835 | 83 |
| SynthSRv2_resampled+co-registered_sagittal_T2w | 0.12 | 0.166 | 0.182 | 70 |
| Recon-all-clinical_resampled_axial_T1w | **0.31** | **9.79e-05** | **3.61e-04** | 143 |
| Recon-all-clinical_resampled_coronal_T1w | **0.37** | **1.74e-06** | **3.55e-05** | 147 |
| Recon-all-clinical_resampled_sagittal_T1w | **0.30** | **1.10e-04** | **3.61e-04** | 142 |
| Recon-all-clinical_resampled+co-registered_axial_T2w | **0.28** | **3.65e-04** | **0.001** | 143 |
| Recon-all-clinical_resampled+co-registered_coronal_T2w | **0.19** | **0.012** | **0.018** | 140 |
| Recon-all-clinical_resampled+co-registered_sagittal_T2w | **0.33** | **3.50e-05** | **2.14e-04** | 140 |
| Recon-all-clinical_multi_T2w | **0.36** | **3.09e-06** | **3.55e-05** | 149 |
| Recon-any_resampled_axial_T1w | 0.07 | 0.210 | 0.220 | 146 |
| Recon-any_resampled_coronal_T1w | **0.20** | **0.007** | **0.013** | 150 |
| Recon-any_resampled+co-registered_axial_T2w | **0.18** | **0.016** | **0.022** | 143 |
| Recon-any_resampled+co-registered_coronal_T2w | **0.22** | **0.004** | **0.009** | 146 |
| Recon-any_resampled+co-registered_sagittal_T2w | **0.20** | **0.008** | **0.014** | 146 |

Intraclass correlation coefficients (ICCs) between processed low-field and 3T global mean cortical thickness, computed using only images rated as good quality (score=3). Images rated as poor (score=1) or fair (score=2) were excluded from the analysis. The number of subjects remaining after exclusion is reported. Pipelines in which all images were rated as good quality were not included in the table. Results significant at $p_{FDR} < 0.05$ after multiple-comparison correction are shown in bold.

**Table S12. Pipelines with the highest Intraclass correlation coefficients (ICCs) across lobes**

| Lobe | Pipeline | ICC | *p* | $p_{FDR}$ |
|---|---|---|---|---|
| Left frontal | Recon-all-clinical_raw_coronal_T1w | 0.35 | 4.60e-06 | 1.84e-04 |
| Left cingulate | Recon-all-clinical_multi_T1w | 0.46 | 2.02e-09 | 8.07e-08 |
| Left insula | Recon-all-clinical_multi_T1w | 0.25 | 8.49e-04 | 0.018 |
| Left temporal | Recon-all-clinical_raw_coronal_T1w | 0.34 | 8.17e-06 | 1.67e-04 |
| Left parietal | Recon-all-clinical_multi_T1w | 0.42 | 3.25e-08 | 1.30e-06 |
| Left occipital | Standard_sagittal_T1w+T2w | 0.27 | 2.63e-04 | 0.011 |
| Right frontal | Recon-all-clinical_multi_T2w | 0.33 | 1.44e-05 | 5.75e-04 |
| Right cingulate | Recon-all-clinical_multi_T2w | 0.51 | 1.63e-11 | 6.52e-10 |
| Right insula | SynthSRv2_multi_T2w | 0.26 | 7.66e-04 | 0.031 |
| Right temporal | Recon-all-clinical_multi_T1w | 0.41 | 9.52e-08 | 3.11e-06 |
| Right parietal | Recon-all-clinical_raw_coronal_T1w | 0.45 | 3.01e-09 | 1.20e-07 |
| Right occipital | Standard_axial_T1w+T2w | 0.28 | 9.79e-05 | 0.004 |

ICCs and corresponding *p*-values for the pipelines showing the highest correlation within each lobe. All correlations remained statistically significant after multiple testing corrections ($p_{FDR} < 0.05$).

**Table S13. Pearson correlations of lobar measures from standard versus recon-all-clinical-processed multi T2w with 3T scans**

| Lobe | Standard coronal T1w+T2w vs. 3T | | | Recon-all-clinical multi T2w vs. 3T | | | Steiger's | | |
|---|---|---|---|---|---|---|---|---|---|
| | ***r*** | ***p*** | ***$p_{FDR}$*** | ***r*** | ***p*** | ***$p_{FDR}$*** | ***Z*** | ***p*** | ***$p_{FDR}$*** |
| Left frontal | 0.09 | 0.276 | 0.414 | **0.34** | **2.06e-05** | **4.12e-05** | **2.59** | **0.010** | **0.039** |
| Left cingulate | 0.10 | 0.236 | 0.405 | **0.42** | **1.02e-07** | **3.07e-07** | **3.35** | **8.05e-04** | **0.005** |
| Left insula | -0.08 | 0.314 | 0.419 | 0.14 | 0.093 | 0.111 | 1.96 | 0.049 | 0.077 |
| Left temporal | 0.07 | 0.406 | 0.487 | **0.27** | **0.001** | **0.002** | 1.92 | 0.055 | 0.077 |
| Left parietal | **0.27** | **9.47e-04** | **0.006** | **0.43** | **4.79e-08** | **1.92e-07** | 1.90 | 0.058 | 0.077 |
| Left occipital | 0.11 | 0.194 | 0.389 | 0.00 | 0.978 | 0.978 | -0.90 | 0.367 | 0.367 |
| Right frontal | 0.19 | 0.020 | 0.059 | **0.34** | **2.00e-05** | **4.12e-05** | 1.53 | 0.126 | 0.137 |
| Right cingulate | 0.01 | 0.912 | 0.912 | **0.51** | **3.60e-11** | **4.32e-10** | **4.95** | **7.57e-07** | **9.08e-06** |
| Right insula | -0.05 | 0.576 | 0.628 | **0.20** | **0.013** | **0.017** | 2.09 | 0.036 | 0.077 |
| Right temporal | 0.13 | 0.100 | 0.239 | **0.34** | **2.63e-05** | **4.51e-05** | 1.96 | 0.050 | 0.077 |
| Right parietal | **0.28** | **5.85e-04** | **0.006** | **0.46** | **2.39e-09** | **1.44e-08** | 2.12 | 0.034 | 0.077 |
| Right occipital | **0.21** | **0.011** | **0.044** | 0.03 | 0.716 | 0.781 | -1.71 | 0.087 | 0.104 |

Pearson correlation coefficients were calculated between 3T scans and 64mT scans processed using the standard coronal T1w+T2w pipeline and the recon-all-clinical multi T2w pipeline. Differences in correlation strength were assessed using Steiger's Z test. Positive *Z* values indicate stronger correspondence with 3T scans for recon-all-clinical–processed regions relative to standard regions. Results that are significant at $p_{FDR} < 0.05$ after correction for multiple comparisons are shown in bold.

**Table S14. Intraclass correlations of lobar measures from standard versus recon-all-clinical-processed multi T2w with 3T scans**

| Lobe | Standard coronal T1w+T2w vs. 3T | | | Recon-all-clinical multi T2w vs. 3T | | | Steiger's | | |
|---|---|---|---|---|---|---|---|---|---|
| | ICC | *p* | $p_{FDR}$ | ICC | *p* | $p_{FDR}$ | *Z* | *p* | $p_{FDR}$ |
| Left frontal | 0.08 | 0.150 | 0.225 | **0.33** | **1.38e-05** | **2.88e-05** | **2.52** | **0.012** | **0.047** |
| Left cingulate | 0.09 | 0.126 | 0.216 | **0.41** | **9.02e-08** | **3.61e-07** | **3.24** | **0.001** | **0.007** |
| Left insula | -0.06 | 0.781 | 0.781 | 0.14 | 0.046 | 0.055 | 1.78 | 0.075 | 0.123 |
| Left temporal | 0.07 | 0.212 | 0.283 | **0.25** | **0.001** | **0.002** | 1.72 | 0.086 | 0.123 |
| Left parietal | **0.23** | **0.002** | **0.012** | **0.4** | **2.15e-07** | **6.45e-07** | 1.77 | 0.078 | 0.123 |
| Left occipital | 0.09 | 0.124 | 0.216 | 0.00 | 0.511 | 0.511 | -0.81 | 0.42 | 0.420 |
| Right frontal | 0.18 | 0.013 | 0.038 | **0.33** | **1.44e-05** | **2.88e-05** | 1.52 | 0.129 | 0.141 |
| Right cingulate | 0.01 | 0.459 | 0.550 | **0.51** | **1.63e-11** | **1.96e-10** | **4.92** | **8.66E-07** | **1.04e-05** |
| Right insula | -0.03 | 0.663 | 0.724 | **0.20** | **0.008** | **0.010** | 1.95 | 0.051 | 0.123 |
| Right temporal | 0.13 | 0.059 | 0.141 | **0.31** | **6.71e-05** | **1.15e-04** | 1.69 | 0.092 | 0.123 |
| Right parietal | **0.25** | **0.001** | **0.012** | **0.43** | **2.28e-08** | **1.37e-07** | 1.91 | 0.056 | 0.123 |
| Right occipital | **0.19** | **0.009** | **0.037** | 0.03 | 0.358 | 0.390 | -1.54 | 0.123 | 0.141 |

Intraclass correlation coefficients (ICCs) were calculated between 3T scans and 64mT scans processed using the standard coronal T1w+T2w pipeline and the recon-all-clinical multi T2w pipeline. Differences in correlation strength were assessed using Steiger's Z test. Positive *Z* values indicate stronger correspondence with 3T scans for recon-all-clinical–processed regions relative to standard regions. Results that are significant at $p_{FDR} < 0.05$ after correction for multiple comparisons are shown in bold.

**Table S15. Pearson correlations of regional measures from standard versus recon-all-clinical-processed multi T2w with 3T scans**

| | Standard coronal T1w and T2w vs. 3T | | | Recon-all-clinical multi T2w vs. 3T | | | Steiger's | | |
|---|---|---|---|---|---|---|---|---|---|
| **Region** | ***r*** | ***p*** | $p_{FDR}$ | ***r*** | ***p*** | $p_{FDR}$ | ***Z*** | ***p*** | $p_{FDR}$ |
| left banks of superior temporal sulcus | 0.05 | 0.526 | 0.575 | **0.28** | **5.34e-04** | **9.79e-04** | **2.19** | **0.028** | **0.041** |
| left caudal anterior cingulate | 0.15 | 0.061 | 0.082 | **0.32** | **7.21e-05** | **1.49e-04** | 1.84 | 0.066 | 0.088 |
| left caudal middle frontal | 0.09 | 0.266 | 0.316 | **0.43** | **3.48e-08** | **9.84e-08** | **3.42** | **6.24e-04** | **0.001** |
| left cuneus | **0.18** | **0.025** | **0.036** | 0.10 | 0.22 | 0.266 | -0.78 | 0.435 | 0.488 |
| left entorhinal | 0.02 | 0.849 | 0.868 | **0.22** | **0.007** | **0.011** | 1.83 | 0.067 | 0.09 |
| left fusiform | -0.04 | 0.62 | 0.664 | 0.10 | 0.234 | 0.28 | 1.24 | 0.215 | 0.261 |
| left inferior parietal | **0.20** | **0.016** | **0.024** | **0.28** | **4.39e-04** | **8.10e-04** | 0.94 | 0.346 | 0.4 |
| left inferior temporal | -0.04 | 0.628 | 0.673 | **0.22** | **7.00e-03** | **1.10e-02** | **2.40** | **0.016** | **0.025** |
| left isthmus cingulate | -0.08 | 0.334 | 0.389 | **0.28** | **6.45e-04** | **0.001** | **3.28** | **0.001** | **0.002** |
| left lateral occipital | -0.03 | 0.758 | 0.788 | -0.09 | 0.275 | 0.325 | -0.54 | 0.592 | 0.639 |
| left lateral orbitofrontal | -0.03 | 0.681 | 0.72 | 0.14 | 8.50e-02 | 0.113 | 1.43 | 0.153 | 0.192 |
| left lingual | 0.10 | 0.235 | 0.282 | -0.06 | 0.438 | 0.491 | -1.32 | 0.186 | 0.23 |
| left medial orbitofrontal | -0.03 | 0.729 | 0.765 | 0.17 | 0.039 | 0.054 | 1.64 | 0.102 | 0.132 |
| left middle temporal | 0.15 | 0.068 | 0.09 | **0.31** | **1.43e-04** | **2.82e-04** | 1.53 | 0.127 | 0.162 |
| left parahippocampal | 0.09 | 0.274 | 0.324 | **0.58** | **6.63e-15** | **2.98e-14** | **5.17** | **2.29e-07** | **6.01e-07** |
| left paracentral | -0.05 | 0.566 | 0.614 | 0.13 | 1.21e-01 | 1.55e-01 | 1.72 | 0.085 | 0.111 |
| left pars triangularis | -0.01 | 0.906 | 0.918 | **0.43** | **4.05e-08** | **1.14e-07** | **4.32** | **1.58e-05** | **3.49e-05** |
| left pars opercularis | 0.06 | 0.478 | 0.531 | **0.19** | **1.80e-02** | **2.70e-02** | 1.16 | 2.46e-01 | 2.94e-01 |
| left pars orbitalis | 0.04 | 0.658 | 0.702 | **0.38** | **1.27e-06** | **3.14e-06** | **3.75** | **1.79e-04** | **3.49e-04** |
| left pericalcarine | 0.16 | 0.045 | 0.063 | -0.15 | 0.059 | 0.08 | **-2.97** | **0.003** | **0.005** |
| left postcentral | 0.14 | 0.099 | 0.129 | **0.21** | **9.00e-03** | **1.40e-02** | 0.71 | 4.78e-01 | 5.31e-01 |
| left posterior cingulate | 0.06 | 0.483 | 0.536 | **0.23** | **0.005** | **0.009** | 1.65 | 0.1 | 0.13 |
| left precentral | -0.01 | 0.871 | 0.888 | 0.13 | 0.118 | 0.152 | 1.34 | 0.179 | 0.222 |
| left precuneus | **0.22** | **0.006** | **0.01** | **0.45** | **6.62e-09** | **1.99e-08** | **2.61** | **0.009** | **0.014** |
| left rostral anterior cingulate | -0.02 | 0.823 | 0.846 | **0.39** | **8.20e-07** | **2.06e-06** | **3.91** | **9.20e-05** | **1.87e-04** |
| left rostral middle frontal | 0.09 | 0.27 | 0.319 | **0.28** | **4.20e-04** | **7.78e-04** | **2.23** | **0.026** | **0.038** |
| left superior frontal | 0.12 | 0.134 | 0.17 | **0.43** | **3.37e-08** | **9.53e-08** | **3.34** | **8.48e-04** | **2.00e-03** |
| left superior parietal | **0.21** | **0.01** | **0.016** | **0.41** | **1.42e-07** | **3.82e-07** | **2.31** | **0.021** | **0.031** |
| left superior temporal | 0.10 | 0.233 | 0.28 | **0.42** | **8.48e-08** | **2.31e-07** | **3.40** | **6.66e-04** | **0.001** |
| left supramarginal | 0.12 | 0.153 | 0.192 | **0.40** | **4.09e-07** | **1.06e-06** | **3.07** | **0.002** | **0.004** |
| left frontal pole | -0.09 | 0.268 | 0.318 | 0.10 | 2.23e-01 | 2.69e-01 | 1.68 | 9.20e-02 | 0.121 |
| left temporal pole | -0.01 | 0.886 | 0.9 | 0.17 | 3.90e-02 | 5.50e-02 | 1.57 | 0.117 | 0.15 |
| left transverse temporal | **-0.23** | **0.006** | **0.009** | **0.17** | **0.034** | **0.048** | **3.66** | **2.51e-04** | **4.77e-04** |
| left insula | -0.08 | 0.326 | 0.38 | 0.14 | 0.082 | 0.108 | 1.98 | 4.80e-02 | 6.60e-02 |
| right banks of superior temporal sulcus | 0.12 | 0.16 | 0.2 | **0.21** | **0.009** | **0.014** | 0.89 | 0.372 | 0.426 |
| right caudal anterior cingulate | -0.11 | 0.177 | 0.22 | **0.47** | **1.20e-09** | **3.83e-09** | **5.46** | **4.70e-08** | **1.32e-07** |
| right caudal middle frontal | 0.16 | 0.053 | 0.073 | **0.22** | **0.007** | **0.012** | 0.61 | 0.545 | 0.594 |

| | | | | | | | | | |
|---|---|---|---|---|---|---|---|---|---|
| right cuneus | 0.09 | 0.272 | 0.321 | 0.07 | 0.363 | 0.417 | -0.14 | 0.886 | 0.9 |
| right entorhinal | -0.05 | 0.565 | 0.614 | **0.22** | **0.007** | **0.012** | **2.19** | **0.029** | **0.041** |
| right fusiform | -0.03 | 0.684 | 0.722 | 0.05 | 0.547 | 0.596 | 0.69 | 0.492 | 0.544 |
| right inferior parietal | **0.19** | **0.017** | **0.026** | **0.28** | **5.58e-04** | **0.001** | 0.82 | 0.413 | 0.466 |
| right inferior temporal | 0.02 | 0.783 | 0.811 | **0.24** | **3.00e-03** | **0.006** | 1.96 | 0.05 | 0.069 |
| right isthmus cingulate | 0.07 | 0.379 | 0.433 | **0.41** | **1.65e-07** | **4.41e-07** | **3.09** | **0.002** | **0.003** |
| right lateral occipital | 0.09 | 0.26 | 0.309 | 0.09 | 0.29 | 0.34 | -0.05 | 0.956 | 0.963 |
| right lateral orbitofrontal | 0.01 | 0.907 | 0.919 | **0.20** | **1.40e-02** | **2.20e-02** | 1.53 | 0.126 | 0.161 |
| right lingual | 0.12 | 0.157 | 0.197 | -0.08 | 0.346 | 0.4 | -1.85 | 0.064 | 0.086 |
| right medial orbitofrontal | **-0.21** | **0.011** | **0.017** | **0.30** | **1.53e-04** | **3.02e-04** | **4.42** | **9.91e-06** | **2.24e-05** |
| right middle temporal | 0.05 | 0.54 | 0.59 | **0.45** | **8.35e-09** | **2.49e-08** | **3.65** | **2.61e-04** | **4.93e-04** |
| right parahippocampal | 0.04 | 0.597 | 0.642 | **0.40** | **3.72e-07** | **9.63e-07** | **3.23** | **1.00e-03** | **2.00e-03** |
| right paracentral | -0.08 | 0.326 | 0.38 | 0.15 | 6.40e-02 | 8.60e-02 | **2.30** | **2.10e-02** | **3.10e-02** |
| right pars triangularis | 0.04 | 0.61 | 0.655 | **0.52** | **1.34e-11** | **4.78e-11** | **4.78** | **1.74e-06** | **4.25e-06** |
| right pars opercularis | 0.07 | 0.401 | 0.455 | **0.39** | **5.97e-07** | **1.53e-06** | **3.15** | **0.002** | **0.003** |
| right pars orbitalis | 0.10 | 0.229 | 0.275 | **0.24** | **3.00e-03** | **5.00e-03** | 1.43 | 1.53e-01 | 1.93e-01 |
| right pericalcarine | 0.01 | 0.879 | 0.895 | -0.07 | 4.24e-01 | 4.77e-01 | -0.73 | 0.467 | 0.52 |
| right postcentral | 0.06 | 0.478 | 0.531 | **0.31** | **8.69e-05** | **1.78e-04** | **2.34** | **0.019** | **0.029** |
| right posterior cingulate | -0.02 | 0.8 | 0.826 | **0.32** | **6.33e-05** | **1.31e-04** | **3.23** | **0.001** | **0.002** |
| right precentral | -0.02 | 0.827 | 0.849 | 0.12 | 1.43e-01 | 1.80e-01 | 1.28 | 0.201 | 0.246 |
| right precuneus | **0.23** | **0.005** | **0.008** | **0.45** | **1.16e-08** | **3.40e-08** | **2.25** | **0.024** | **0.036** |
| right rostral anterior cingulate | 0.00 | 0.979 | 0.981 | **0.31** | **1.13e-04** | **2.26e-04** | **2.87** | **0.004** | **0.007** |
| right rostral middle frontal | **0.22** | **0.007** | **0.011** | **0.25** | **2.00e-03** | **3.00e-03** | 0.34 | 0.738 | 0.769 |
| right superior frontal | **0.26** | **0.001** | **0.002** | **0.39** | **7.14e-07** | **1.81e-06** | 1.39 | 0.163 | 0.204 |
| right superior parietal | **0.24** | **0.004** | **0.006** | **0.43** | **4.89e-08** | **1.37e-07** | **2.19** | **0.029** | **0.041** |
| right superior temporal | 0.13 | 0.103 | 0.133 | **0.51** | **2.39e-11** | **8.33e-11** | **4.19** | **2.82e-05** | **6.07e-05** |
| right supramarginal | 0.08 | 0.354 | 0.408 | **0.31** | **9.06e-05** | **1.85e-04** | **2.36** | **0.018** | **0.027** |
| right frontal pole | -0.11 | 0.162 | 0.203 | **0.17** | **3.30e-02** | **4.70e-02** | **2.59** | **1.00e-02** | **1.50e-02** |
| right temporal pole | 0.09 | 0.293 | 0.343 | **0.20** | **1.40e-02** | **2.20e-02** | 0.97 | 0.331 | 0.385 |
| right transverse temporal | 0.00 | 0.974 | 0.978 | 0.13 | 0.125 | 0.159 | 1.27 | 0.203 | 0.247 |
| right insula | -0.05 | 0.576 | 0.624 | **0.20** | **0.013** | **0.019** | 2.09 | 0.036 | 0.051 |

Pearson correlation coefficients were calculated between 3T scans and 64mT scans processed using the standard coronal T1w+T2w pipeline and the recon-all-clinical multi T2w pipeline. Differences in correlation strength were assessed using Steiger's Z test. Positive *Z* values indicate stronger correspondence with 3T scans for recon-all-clinical–processed regions relative to standard regions. Results that are significant at $p_{FDR} < 0.05$ after correction for multiple comparisons are shown in bold.

**Table S16. Intraclass correlations of regional measures from standard versus recon-all-clinical-processed multi T2w with 3T scans**

| Region | Standard coronal T1w and T2w vs. 3T | | | Recon-all-clinical multi T2w vs. 3T | | | Steiger's | | |
|---|---|---|---|---|---|---|---|---|---|
| | ICC | *p* | $p_{FDR}$ | ICC | *p* | $p_{FDR}$ | *Z* | *p* | $p_{FDR}$ |
| left banks of superior temporal sulcus | 0.04 | 0.306 | 0.369 | **0.28** | **2.78e-04** | **5.51e-04** | **2.22** | **0.027** | **0.04** |
| left caudal anterior cingulate | 0.14 | 0.04 | 0.058 | **0.32** | **3.43e-05** | **7.65e-05** | 1.92 | 0.054 | 0.077 |
| left caudal middle frontal | 0.07 | 0.191 | 0.242 | **0.41** | **6.44e-08** | **1.92e-07** | **3.29** | **9.90e-04** | **0.002** |
| left cuneus | 0.13 | 0.057 | 0.081 | 0.1 | 0.112 | 0.149 | -0.27 | 0.787 | 0.828 |
| left entorhinal | 6.50e-03 | 0.469 | 0.537 | **0.2** | **0.008** | **0.013** | 1.67 | 0.095 | 0.128 |
| left fusiform | -0.03 | 0.652 | 0.712 | 0.1 | 0.116 | 0.153 | 1.15 | 0.25 | 0.309 |
| left inferior parietal | **0.17** | **0.02** | **0.03** | **0.28** | **2.93e-04** | **5.77e-04** | 1.11 | 0.267 | 0.326 |
| left inferior temporal | -0.03 | 0.659 | 0.717 | **0.22** | **4.00e-03** | **7.00e-03** | **2.26** | **0.024** | **0.036** |
| left isthmus cingulate | -0.07 | 0.798 | 0.837 | **0.27** | **3.29e-04** | **6.43e-04** | **3.13** | **0.002** | **0.003** |
| left lateral occipital | -0.02 | 0.603 | 0.667 | -0.09 | 0.863 | 0.889 | -0.57 | 0.567 | 0.635 |
| left lateral orbitofrontal | -0.03 | 0.65 | 0.711 | 0.14 | 4.20e-02 | 6.10e-02 | 1.42 | 0.156 | 0.202 |
| left lingual | 0.08 | 0.156 | 0.201 | -0.06 | 0.78 | 0.822 | -1.2 | 0.229 | 0.285 |
| left medial orbitofrontal | -0.03 | 0.627 | 0.688 | **0.17** | **0.021** | **0.032** | 1.59 | 0.111 | 0.148 |
| left middle temporal | 0.14 | 0.045 | 0.064 | **0.3** | **1.07e-04** | **2.26e-04** | 1.52 | 0.129 | 0.17 |
| left parahippocampal | 0.04 | 0.313 | 0.377 | **0.56** | **5.18e-14** | **2.28e-13** | **5.16** | **2.43e-07** | **6.80e-07** |
| left paracentral | -0.03 | 0.658 | 0.716 | 0.13 | 6.00e-02 | 8.40e-02 | 1.51 | 0.131 | 0.171 |
| left pars triangularis | -8.00e-03 | 0.539 | 0.609 | **0.42** | **3.31e-08** | **1.00e-07** | **4.12** | **3.77e-05** | **8.37e-05** |
| left pars opercularis | 0.05 | 0.252 | 0.31 | **0.19** | **1.00e-02** | **1.60e-02** | 1.17 | 2.43e-01 | 3.00e-01 |
| left pars orbitalis | 0.03 | 0.345 | 0.412 | **0.37** | **1.43e-06** | **3.70e-06** | **3.46** | **5.32e-04** | **0.001** |
| left pericalcarine | 0.11 | 0.085 | 0.116 | -0.15 | 9.70e-01 | 9.81e-01 | **-2.43** | **1.50e-02** | **0.024** |
| left postcentral | 1.10e-01 | 0.087 | 0.118 | **0.20** | **7.00e-03** | **1.20e-02** | 0.79 | 4.30e-01 | 4.98e-01 |
| left posterior cingulate | 0.05 | 0.276 | 0.337 | **0.22** | **0.003** | **0.005** | 1.65 | 0.098 | 0.132 |
| left precentral | -0.01 | 0.554 | 0.624 | 0.11 | 0.083 | 0.113 | 1.13 | 0.257 | 0.315 |
| left precuneus | **0.17** | **0.016** | **0.026** | **0.44** | **5.41e-09** | **1.73e-08** | **2.86** | **0.004** | **0.007** |
| left rostral anterior cingulate | -0.02 | 0.575 | 0.641 | **0.39** | **5.20e-07** | **1.40e-06** | **3.81** | **1.40e-04** | **2.87e-04** |
| left rostral middle frontal | 0.08 | 0.16 | 0.206 | **0.28** | **2.85e-04** | **5.63e-04** | 2.1 | 0.036 | 0.052 |
| left superior frontal | 0.12 | 0.079 | 0.109 | **0.43** | **2.01e-08** | **6.17e-08** | **3.31** | **9.43e-04** | **2.00e-03** |
| left superior parietal | **0.17** | **0.018** | **0.028** | **0.40** | **1.87e-07** | **5.29e-07** | **2.37** | **0.018** | **0.027** |
| left superior temporal | 0.09 | 0.141 | 0.184 | **0.40** | **1.44e-07** | **4.11e-07** | **3.18** | **1.00e-03** | **0.003** |
| left supramarginal | 0.09 | 0.134 | 0.175 | **0.38** | **5.37e-07** | **1.45e-06** | **2.95** | **0.003** | **0.005** |
| left frontal pole | -0.08 | 0.835 | 0.867 | 0.10 | 1.12e-01 | 1.49e-01 | 1.57 | 0.116 | 0.153 |
| left temporal pole | -6.60e-03 | 0.532 | 0.602 | **0.17** | **2.10e-02** | **3.20e-02** | 1.5 | 0.134 | 0.175 |
| left transverse temporal | -1.70e-01 | 0.98 | 0.987 | **0.16** | **0.027** | **0.04** | **2.88** | **0.004** | **0.007** |
| left insula | -0.06 | 0.776 | 0.82 | 0.14 | 0.041 | 0.059 | 1.81 | 0.071 | 0.099 |
| right banks of superior temporal sulcus | 0.08 | 0.153 | 0.198 | **0.21** | **0.005** | **0.009** | 1.12 | 0.265 | 0.324 |
| right caudal anterior cingulate | -0.09 | 0.858 | 0.886 | **0.47** | **7.15e-10** | **2.43e-09** | **5.19** | **2.09e-07** | **5.85e-07** |
| right caudal middle frontal | 0.12 | 0.069 | 0.096 | **0.21** | **0.005** | **0.009** | 0.82 | 0.415 | 0.483 |

| | | | | | | | | | |
|---|---|---|---|---|---|---|---|---|---|
| right cuneus | 0.07 | 0.21 | 0.264 | 0.07 | 0.182 | 0.232 | 0.08 | 0.94 | 0.956 |
| right entorhinal | -0.03 | 0.639 | 0.7 | **0.21** | **0.004** | **0.007** | 2.04 | 0.041 | 0.059 |
| right fusiform | -0.03 | 0.622 | 0.685 | 0.05 | 0.274 | 0.334 | 0.62 | 0.533 | 0.604 |
| right inferior parietal | **0.15** | **0.031** | **0.046** | **0.27** | **4.16e-04** | **8.00e-04** | 1.09 | 0.277 | 0.337 |
| right inferior temporal | 0.02 | 0.414 | 0.482 | **0.23** | **2.00e-03** | **3.00e-03** | 1.93 | 0.054 | 0.076 |
| right isthmus cingulate | 0.06 | 0.224 | 0.28 | **0.41** | **8.13e-08** | **2.40e-07** | **3.16** | **0.002** | **0.003** |
| right lateral occipital | 0.08 | 0.16 | 0.205 | 0.09 | 0.147 | 0.191 | 0.04 | 0.967 | 0.979 |
| right lateral orbitofrontal | 8.50e-03 | 0.459 | 0.527 | **0.2** | **7.00e-03** | **1.20e-02** | 1.55 | 0.121 | 0.159 |
| right lingual | 0.11 | 0.091 | 0.124 | -0.08 | 0.824 | 0.859 | -1.77 | 0.077 | 0.106 |
| right medial orbitofrontal | -2.00e-01 | 0.992 | 0.995 | **0.3** | **7.71e-05** | **1.65e-04** | **4.31** | **1.63e-05** | **3.73e-05** |
| right middle temporal | 0.05 | 0.29 | 0.351 | **0.44** | **6.73e-09** | **2.14e-08** | **3.62** | **2.91e-04** | **5.73e-04** |
| right parahippocampal | 0.02 | 0.42 | 0.488 | **0.4** | **1.70e-07** | **4.85e-07** | **3.46** | **5.38e-04** | **1.00e-03** |
| right paracentral | -0.06 | 0.751 | 0.799 | **0.15** | **3.20e-02** | **4.70e-02** | 1.96 | 5.00e-02 | 7.10e-02 |
| right pars triangularis | 0.03 | 0.344 | 0.411 | **0.52** | **6.15e-12** | **2.37e-11** | **4.77** | **1.83e-06** | **4.64e-06** |
| right pars opercularis | 0.06 | 0.226 | 0.281 | **0.39** | **4.85e-07** | **1.31e-06** | **3.09** | **2.00e-03** | **0.004** |
| right pars orbitalis | 0.08 | 0.154 | 0.199 | **0.24** | **2.00e-03** | **3.00e-03** | 1.51 | 0.132 | 0.173 |
| right pericalcarine | 0.01 | 0.449 | 0.518 | -0.07 | 0.788 | 0.829 | -0.7 | 0.485 | 0.553 |
| right postcentral | 0.05 | 0.258 | 0.317 | **0.27** | **3.30e-04** | **6.45e-04** | 1.98 | 4.80e-02 | 6.80e-02 |
| right posterior cingulate | -0.02 | 0.582 | 0.648 | **0.32** | **4.00e-05** | **8.84e-05** | **3.12** | **0.002** | **0.003** |
| right precentral | -0.02 | 0.583 | 0.648 | 0.09 | 1.23e-01 | 1.62e-01 | 1.01 | 0.314 | 0.378 |
| right precuneus | **0.18** | **0.015** | **0.024** | **0.44** | **5.41e-09** | **1.73e-08** | **2.68** | **0.007** | **0.012** |
| right rostral anterior cingulate | 1.70e-03 | 0.492 | 0.561 | **0.31** | **5.38e-05** | **1.17e-04** | **2.84** | **0.005** | **0.008** |
| right rostral middle frontal | **0.2** | **0.007** | **0.012** | **0.25** | **9.86e-04** | **2.00e-03** | 0.5 | 0.618 | 0.682 |
| right superior frontal | **2.40e-01** | **0.001** | **0.002** | **0.39** | **4.27e-07** | **1.17e-06** | 1.47 | 0.141 | 0.184 |
| right superior parietal | **0.2** | **0.007** | **0.012** | **0.42** | **3.33e-08** | **1.01e-07** | **2.4** | **0.016** | **0.025** |
| right superior temporal | 0.12 | 0.074 | 0.102 | **0.49** | **5.89e-11** | **2.10e-10** | **3.96** | **7.47e-05** | **1.61e-04** |
| right supramarginal | 0.06 | 0.233 | 0.29 | **0.29** | **1.25e-04** | **2.59e-04** | **2.2** | **0.028** | **0.042** |
| right frontal pole | -0.11 | 0.913 | 0.934 | **0.17** | **1.60e-02** | **2.60e-02** | **2.56** | **1.10e-02** | **1.70e-02** |
| right temporal pole | 0.07 | 0.185 | 0.236 | **0.2** | **8.00e-03** | **1.30e-02** | 1.06 | 0.288 | 0.349 |
| right transverse temporal | -1.90e-03 | 0.509 | 0.579 | 0.12 | 0.073 | 0.101 | 1.11 | 0.265 | 0.325 |
| right insula | -3.00e-02 | 0.663 | 0.72 | **0.2** | **0.008** | **0.013** | 1.95 | 0.051 | 0.073 |

Intraclass correlation coefficients (ICCs) were calculated between 3T scans and 64mT scans processed using the standard coronal T1w+T2w pipeline and the recon-all-clinical multi T2w pipeline. Differences in correlation strength were assessed using Steiger's Z test. Positive *Z* values indicate stronger correspondence with 3T scans for recon-all-clinical–processed regions relative to standard regions. Results that are significant at $p_{FDR} < 0.05$ after correction for multiple comparisons are shown in bold.

**Table S17. Pearson correlations of regional measures from standard versus recon-all-clinical-processed multi T1w with 3T scans**

| Region | Standard coronal T1w and T2w vs. 3T | | | Recon-all-clinical multi T1w vs. 3T | | | Steiger | | |
|---|---|---|---|---|---|---|---|---|---|
| | *r* | *p* | $p_{FDR}$ | *r* | *p* | $p_{FDR}$ | *Z* | *p* | $p_{FDR}$ |
| left banks of superior temporal sulcus | 0.05 | 0.526 | 0.572 | **0.33** | **3.49e-05** | **6.98e-05** | **2.68** | **0.007** | **0.011** |
| left caudal anterior cingulate | 0.15 | 0.061 | 0.081 | **0.42** | **1.10e-07** | **2.77e-07** | **2.89** | **0.004** | **0.006** |
| left caudal middle frontal | 0.09 | 0.266 | 0.314 | **0.22** | **0.007** | **0.011** | 1.09 | 0.274 | 0.322 |
| left cuneus | **0.18** | **0.025** | **0.036** | -0.03 | 0.683 | 0.719 | -1.85 | 0.064 | 0.085 |
| left entorhinal | 0.02 | 0.849 | 0.869 | 0.10 | 0.213 | 0.257 | 0.74 | 0.460 | 0.508 |
| left fusiform | -0.04 | 0.620 | 0.662 | **0.23** | **0.004** | **0.007** | **2.31** | **0.021** | **0.030** |
| left inferior parietal | **0.20** | **0.016** | **0.023** | **0.36** | **4.74e-06** | **1.03e-05** | 1.61 | 0.108 | 0.138 |
| left inferior temporal | -0.04 | 0.628 | 0.671 | **0.32** | **7.15e-05** | **1.38e-04** | **3.11** | **0.002** | **0.003** |
| left isthmus cingulate | -0.08 | 0.334 | 0.385 | **0.28** | **5.19e-04** | **9.11e-04** | **3.39** | **7.01e-04** | **0.001** |
| left lateral occipital | -0.03 | 0.758 | 0.785 | 0.05 | 0.506 | 0.553 | 0.63 | 0.529 | 0.576 |
| left lateral orbitofrontal | -0.03 | 0.681 | 0.718 | **0.21** | **0.011** | **0.017** | 1.94 | 0.053 | 0.071 |
| left lingual | 0.10 | 0.235 | 0.280 | 0.07 | 0.421 | 0.469 | -0.25 | 0.801 | 0.824 |
| left medial orbitofrontal | -0.03 | 0.729 | 0.761 | **0.28** | **4.39e-04** | **7.76e-04** | **2.55** | **0.011** | **0.016** |
| left middle temporal | 0.15 | 0.068 | 0.089 | **0.43** | **3.99e-08** | **1.04e-07** | **2.62** | **0.009** | **0.014** |
| left parahippocampal | 0.09 | 0.274 | 0.322 | **0.57** | **2.26e-14** | **8.78e-14** | **5.17** | **2.37e-07** | **5.79e-07** |
| left paracentral | -0.05 | 0.566 | 0.612 | 0.13 | 0.113 | 0.144 | 1.58 | 0.113 | 0.144 |
| left pars triangularis | -0.01 | 0.906 | 0.918 | **0.53** | **4.97e-12** | **1.71e-11** | **4.92** | **8.68e-07** | **2.04e-06** |
| left pars opercularis | 0.06 | 0.478 | 0.526 | 0.14 | 0.092 | 0.119 | 0.72 | 0.473 | 0.521 |
| left pars orbitalis | 0.04 | 0.658 | 0.698 | **0.36** | **5.96e-06** | **1.28e-05** | **3.08** | **0.002** | **0.003** |
| left pericalcarine | 0.16 | 0.045 | 0.062 | -0.07 | 0.397 | 0.446 | **-2.20** | **0.028** | **0.040** |
| left postcentral | 0.14 | 0.099 | 0.127 | **0.35** | **1.19e-05** | **2.50e-05** | 1.82 | 0.068 | 0.090 |
| left posterior cingulate | 0.06 | 0.483 | 0.531 | **0.38** | **1.16e-06** | **2.70e-06** | **3.33** | **8.81e-04** | **0.002** |
| left precentral | -0.01 | 0.871 | 0.889 | **0.28** | **5.61e-04** | **9.76e-04** | **2.52** | **0.012** | **0.018** |
| left precuneus | **0.22** | **0.006** | **0.010** | **0.44** | **2.33e-08** | **6.19e-08** | **2.17** | **0.030** | **0.043** |
| left rostral anterior cingulate | -0.02 | 0.823 | 0.845 | **0.36** | **4.47e-06** | **9.76e-06** | **3.80** | **1.47e-04** | **2.74e-04** |
| left rostral middle frontal | 0.09 | 0.270 | 0.317 | **0.29** | **2.55e-04** | **4.65e-04** | 1.81 | 0.070 | 0.092 |
| left superior frontal | 0.12 | 0.134 | 0.168 | **0.28** | **5.24e-04** | **9.18e-04** | 1.35 | 0.176 | 0.215 |
| left superior parietal | **0.21** | **0.01** | **0.016** | **0.33** | **3.02e-05** | **6.09e-05** | 1.21 | 0.228 | 0.272 |
| left superior temporal | 0.10 | 0.233 | 0.278 | **0.49** | **2.39e-10** | **7.46e-10** | **3.94** | **8.23e-05** | **1.58e-04** |
| left supramarginal | 0.12 | 0.153 | 0.189 | **0.45** | **9.09e-09** | **2.52e-08** | **3.41** | **6.48e-04** | **0.001** |
| left frontal pole | -0.09 | 0.268 | 0.316 | 0.06 | 0.463 | 0.511 | 1.42 | 0.154 | 0.191 |
| left temporal pole | -0.01 | 0.886 | 0.902 | **0.25** | **0.002** | **0.004** | **2.18** | **0.029** | **0.041** |
| left transverse temporal | **-0.23** | **0.006** | **0.009** | 0.00 | 0.980 | 0.982 | 2.05 | 0.040 | 0.055 |
| left insula | -0.08 | 0.326 | 0.377 | **0.26** | **0.001** | **0.002** | **2.98** | **0.003** | **0.005** |
| right banks of superior temporal sulcus | 0.12 | 0.160 | 0.197 | **0.46** | **3.23e-09** | **9.25e-09** | **3.21** | **0.001** | **0.002** |
| right caudal anterior cingulate | -0.11 | 0.177 | 0.217 | **0.43** | **5.45e-08** | **1.40e-07** | **4.95** | **7.28e-07** | **1.72e-06** |
| right caudal middle frontal | 0.16 | 0.053 | 0.071 | **0.24** | **0.003** | **0.005** | 0.76 | 0.450 | 0.499 |
| right cuneus | 0.09 | 0.272 | 0.319 | 0.03 | 0.704 | 0.737 | -0.48 | 0.633 | 0.674 |

| | | | | | | | | | |
|---|---|---|---|---|---|---|---|---|---|
| right entorhinal | -0.05 | 0.565 | 0.611 | 0.01 | 0.905 | 0.918 | 0.47 | 0.636 | 0.677 |
| right fusiform | -0.03 | 0.684 | 0.720 | 0.15 | 0.062 | 0.082 | 1.49 | 0.136 | 0.169 |
| right inferior parietal | **0.19** | **0.017** | **0.025** | **0.36** | **7.14e-06** | **1.53e-05** | 1.48 | 0.139 | 0.173 |
| right inferior temporal | 0.02 | 0.783 | 0.808 | **0.32** | **5.21e-05** | **1.02e-04** | **2.56** | **0.010** | **0.016** |
| right isthmus cingulate | 0.07 | 0.379 | 0.429 | **0.31** | **1.29e-04** | **2.43e-04** | 2.05 | 0.041 | 0.056 |
| right lateral occipital | 0.09 | 0.26 | 0.307 | 0.04 | 0.614 | 0.656 | -0.42 | 0.673 | 0.712 |
| right lateral orbitofrontal | 0.01 | 0.907 | 0.919 | **0.18** | **0.030** | **0.042** | 1.31 | 0.191 | 0.231 |
| right lingual | 0.12 | 0.157 | 0.194 | 0.16 | 0.045 | 0.061 | 0.41 | 0.683 | 0.719 |
| right medial orbitofrontal | **-0.21** | **0.011** | **0.016** | **0.21** | **0.009** | **0.013** | **3.41** | **6.51e-04** | **0.001** |
| right middle temporal | 0.05 | 0.540 | 0.586 | **0.54** | **1.29e-12** | **4.61e-12** | **4.36** | **1.29e-05** | **2.68e-05** |
| right parahippocampal | 0.04 | 0.597 | 0.641 | **0.49** | **1.89e-10** | **5.93e-10** | **4.19** | **2.77e-05** | **5.61e-05** |
| right paracentral | -0.08 | 0.326 | 0.377 | 0.08 | 0.323 | 0.374 | 1.43 | 0.152 | 0.188 |
| right pars triangularis | 0.04 | 0.61 | 0.653 | **0.42** | **9.50e-08** | **2.40e-07** | **3.42** | **6.30e-04** | **0.001** |
| right pars opercularis | 0.07 | 0.401 | 0.45 | **0.43** | **5.23e-08** | **1.35e-07** | **3.62** | **2.98e-04** | **5.40e-04** |
| right pars orbitalis | 0.10 | 0.229 | 0.273 | **0.26** | **0.002** | **0.003** | 1.40 | 0.162 | 0.199 |
| right pericalcarine | 0.01 | 0.879 | 0.896 | -0.08 | 0.345 | 0.396 | -0.80 | 0.421 | 0.470 |
| right postcentral | 0.06 | 0.478 | 0.526 | **0.36** | **6.27e-06** | **1.35e-05** | **2.58** | **0.010** | **0.015** |
| right posterior cingulate | -0.02 | 0.80 | 0.823 | **0.29** | **3.14e-04** | **5.65e-04** | **2.82** | **0.005** | **0.008** |
| right precentral | -0.02 | 0.827 | 0.848 | **0.24** | **0.003** | **0.004** | **2.14** | **0.033** | **0.046** |
| right precuneus | **0.23** | **0.005** | **0.008** | **0.30** | **1.54e-04** | **2.89e-04** | 0.67 | 0.503 | 0.550 |
| right rostral anterior cingulate | 0.00 | 0.979 | 0.982 | 0.17 | 0.039 | 0.054 | 1.53 | 0.127 | 0.160 |
| right rostral middle frontal | **0.22** | **0.007** | **0.011** | 0.13 | 0.107 | 0.136 | -0.75 | 0.456 | 0.504 |
| right superior frontal | **0.26** | **0.001** | **0.002** | **0.22** | **0.006** | **0.009** | -0.31 | 0.756 | 0.783 |
| right superior parietal | **0.24** | **0.004** | **0.006** | **0.32** | **7.45e-05** | **1.43e-04** | 0.76 | 0.445 | 0.494 |
| right superior temporal | 0.13 | 0.103 | 0.132 | **0.48** | **5.06e-10** | **1.54e-09** | **3.60** | **3.14e-04** | **5.65e-04** |
| right supramarginal | 0.08 | 0.354 | 0.404 | **0.41** | **1.96e-07** | **4.84e-07** | **2.98** | **0.003** | **0.005** |
| right frontal pole | -0.11 | 0.162 | 0.20 | **0.23** | **0.005** | **0.008** | **3.05** | **0.002** | **0.004** |
| right temporal pole | 0.09 | 0.293 | 0.340 | **0.22** | **0.007** | **0.011** | 1.20 | 0.230 | 0.273 |
| right transverse temporal | 0.00 | 0.974 | 0.979 | 0.14 | 0.089 | 0.116 | 1.41 | 0.159 | 0.196 |
| right insula | -0.05 | 0.576 | 0.622 | **0.19** | **0.017** | **0.025** | 2.01 | 0.045 | 0.061 |

Pearson correlation coefficients were calculated between 3T scans and 64mT scans processed using the standard coronal T1w+T2w pipeline and the recon-all-clinical multi T1w pipeline. Differences in correlation strength were assessed using Steiger's Z test. Positive *Z* values indicate stronger correspondence with 3T scans for recon-all-clinical–processed regions relative to standard regions. Results that are significant at $p_{FDR} < 0.05$ after correction for multiple comparisons are shown in bold.

**Table S18. Intraclass correlations of regional measures from standard versus recon-all-clinical-processed multi T1w with 3T scans**

| Region | Standard coronal T1w and T2w vs. 3T | | | Recon-all-clinical multi T1w vs. 3T | | | Steiger | | |
|---|---|---|---|---|---|---|---|---|---|
| | ICC | *p* | $p_{FDR}$ | ICC | *p* | $p_{FDR}$ | *Z* | *p* | $p_{FDR}$ |
| left banks of superior temporal sulcus | 0.04 | 0.306 | 0.365 | **0.33** | **1.65e-05** | **3.54e-05** | **2.73** | **0.006** | **0.010** |
| left caudal anterior cingulate | 0.14 | 0.040 | 0.057 | **0.42** | **5.32e-08** | **1.49e-07** | **2.96** | **0.003** | **0.005** |
| left caudal middle frontal | 0.07 | 0.191 | 0.24 | **0.22** | **0.004** | **0.006** | 1.25 | 0.211 | 0.263 |
| left cuneus | 0.13 | 0.057 | 0.08 | -0.03 | 0.658 | 0.707 | -1.39 | 0.166 | 0.211 |
| left entorhinal | 6.50e-03 | 0.469 | 0.529 | 0.10 | 0.108 | 0.143 | 0.81 | 0.418 | 0.479 |
| left fusiform | -0.03 | 0.652 | 0.704 | **0.22** | **0.003** | **0.005** | **2.15** | **0.031** | **0.046** |
| left inferior parietal | **0.17** | **0.020** | **0.030** | **0.36** | **2.39e-06** | **5.58e-06** | 1.85 | 0.064 | 0.089 |
| left inferior temporal | -0.03 | 0.659 | 0.708 | **0.31** | **4.46e-05** | **9.10e-05** | **3.01** | **0.003** | **0.004** |
| left isthmus cingulate | -0.07 | 0.798 | 0.835 | **0.28** | **2.53e-04** | **4.75e-04** | **3.24** | **0.001** | **0.002** |
| left lateral occipital | -0.02 | 0.603 | 0.658 | 0.05 | 0.276 | 0.332 | 0.55 | 0.581 | 0.639 |
| left lateral orbitofrontal | -0.03 | 0.65 | 0.702 | **0.21** | **0.006** | **0.009** | 1.93 | 0.054 | 0.076 |
| left lingual | 0.08 | 0.156 | 0.20 | 0.06 | 0.22 | 0.272 | -0.16 | 0.874 | 0.898 |
| left medial orbitofrontal | -0.03 | 0.627 | 0.681 | **0.27** | **4.89e-04** | **8.88e-04** | **2.39** | **0.017** | **0.026** |
| left middle temporal | 0.14 | 0.045 | 0.064 | **0.43** | **1.82e-08** | **5.28e-08** | **2.71** | **0.007** | **0.0110** |
| left parahippocampal | 0.04 | 0.313 | 0.372 | **0.57** | **1.11e-14** | **4.57e-14** | **5.30** | **1.17e-07** | **3.13e-07** |
| left paracentral | -0.03 | 0.658 | 0.707 | 0.13 | 0.058 | 0.081 | 1.43 | 0.154 | 0.198 |
| left pars triangularis | -8.00e-03 | 0.539 | 0.599 | **0.51** | **1.07e-11** | **3.76e-11** | **4.76** | **1.92e-06** | **4.53e-06** |
| left pars opercularis | 0.05 | 0.252 | 0.307 | 0.14 | 0.048 | 0.067 | 0.73 | 0.463 | 0.523 |
| left pars orbitalis | 0.03 | 0.345 | 0.407 | **0.36** | **3.13e-06** | **7.19e-06** | **3.07** | **0.002** | **0.004** |
| left pericalcarine | 0.11 | 0.085 | 0.114 | -0.07 | 0.79 | 0.827 | -1.65 | 0.099 | 0.131 |
| left postcentral | 0.11 | 0.087 | 0.116 | **0.33** | **1.73e-05** | **3.70e-05** | 1.89 | 0.059 | 0.083 |
| left posterior cingulate | 0.05 | 0.276 | 0.332 | **0.38** | **5.39e-07** | **1.35e-06** | **3.34** | **8.43e-04** | **0.001** |
| left precentral | -0.01 | 0.554 | 0.615 | **0.27** | **3.59e-04** | **6.62e-04** | **2.45** | **0.014** | **0.022** |
| left precuneus | **0.17** | **0.016** | **0.025** | **0.44** | **1.17e-08** | **3.41e-08** | **2.59** | **0.010** | **0.015** |
| left rostral anterior cingulate | -0.02 | 0.575 | 0.633 | **0.35** | **4.95e-06** | **1.12e-05** | **3.60** | **3.18e-04** | **5.91e-04** |
| left rostral middle frontal | 0.08 | 0.16 | 0.205 | **0.29** | **1.25e-04** | **2.41e-04** | 1.89 | 0.059 | 0.082 |
| left superior frontal | 0.12 | 0.079 | 0.108 | **0.28** | **2.54e-04** | **4.77e-04** | 1.42 | 0.156 | 0.20 |
| left superior parietal | **0.17** | **0.018** | **0.027** | **0.33** | **1.58e-05** | **3.42e-05** | 1.52 | 0.128 | 0.167 |
| left superior temporal | 0.09 | 0.141 | 0.183 | **0.49** | **1.14e-10** | **3.83e-10** | **3.98** | **6.96e-05** | **1.39e-04** |
| left supramarginal | 0.09 | 0.134 | 0.175 | **0.44** | **1.04e-08** | **3.07e-08** | **3.40** | **6.64e-04** | **0.001** |
| left frontal pole | -0.08 | 0.835 | 0.866 | 0.06 | 0.238 | 0.291 | 1.30 | 0.195 | 0.245 |
| left temporal pole | -6.60e-03 | 0.532 | 0.593 | **0.25** | **0.001** | **0.002** | **2.17** | **0.030** | **0.044** |
| left transverse temporal | -0.17 | 0.980 | 0.989 | 1.80e-03 | 0.491 | 0.553 | 1.48 | 0.138 | 0.179 |
| left insula | -0.06 | 0.776 | 0.816 | **0.26** | **6.65e-04** | **0.001** | **2.81** | **0.005** | **0.008** |
| right banks of superior temporal sulcus | 0.08 | 0.153 | 0.197 | **0.45** | **2.26e-09** | **6.95e-09** | **3.43** | **6.03e-04** | **0.001** |
| right caudal anterior cingulate | -0.09 | 0.858 | 0.886 | **0.42** | **3.81e-08** | **1.08e-07** | **4.67** | **2.97e-06** | **6.86e-06** |
| right caudal middle frontal | 0.12 | 0.069 | 0.095 | **0.23** | **0.002** | **0.004** | 0.98 | 0.329 | 0.389 |

| | | | | | | | | | |
|---|---|---|---|---|---|---|---|---|---|
| right cuneus | 0.07 | 0.210 | 0.261 | 0.03 | 0.357 | 0.417 | -0.29 | 0.769 | 0.809 |
| right entorhinal | -0.03 | 0.639 | 0.692 | 9.70e-03 | 0.453 | 0.512 | 0.33 | 0.745 | 0.789 |
| right fusiform | -0.03 | 0.622 | 0.677 | **0.15** | **0.033** | **0.048** | 1.42 | 0.155 | 0.199 |
| right inferior parietal | **0.15** | **0.031** | **0.045** | **0.35** | **4.04e-06** | **9.23e-06** | 1.82 | 0.069 | 0.095 |
| right inferior temporal | 0.02 | 0.414 | 0.475 | **0.32** | **2.57e-05** | **5.38e-05** | **2.62** | **0.009** | **0.014** |
| right isthmus cingulate | 0.06 | 0.224 | 0.276 | **0.31** | **6.13e-05** | **1.23e-04** | **2.14** | **0.033** | **0.048** |
| right lateral occipital | 0.08 | 0.160 | 0.204 | 0.04 | 0.308 | 0.366 | -0.33 | 0.738 | 0.783 |
| right lateral orbitofrontal | 8.50e-03 | 0.459 | 0.519 | **0.18** | **0.015** | **0.023** | 1.33 | 0.183 | 0.231 |
| right lingual | 0.11 | 0.091 | 0.122 | **0.16** | **0.024** | **0.036** | 0.44 | 0.660 | 0.708 |
| right medial orbitofrontal | -0.2 | 0.992 | 0.995 | **0.20** | **0.006** | **0.010** | **3.22** | **0.001** | **0.002** |
| right middle temporal | 0.05 | 0.290 | 0.346 | **0.53** | **7.51e-13** | **2.84e-12** | **4.41** | **1.05e-05** | **2.30e-05** |
| right parahippocampal | 0.02 | 0.420 | 0.480 | **0.49** | **1.09e-10** | **3.67e-10** | **4.37** | **1.24e-05** | **2.69e-05** |
| right paracentral | -0.06 | 0.751 | 0.793 | 0.08 | 0.161 | 0.206 | 1.20 | 0.231 | 0.283 |
| right pars triangularis | 0.03 | 0.344 | 0.406 | **0.40** | **1.90e-07** | **4.97e-07** | **3.30** | **9.61e-04** | **0.002** |
| right pars opercularis | 0.06 | 0.226 | 0.278 | **0.43** | **2.54e-08** | **7.25e-08** | **3.62** | **2.92e-04** | **5.44e-04** |
| right pars orbitalis | 0.08 | 0.154 | 0.198 | **0.25** | **0.001** | **0.002** | 1.47 | 0.142 | 0.185 |
| right pericalcarine | 0.01 | 0.449 | 0.509 | -0.07 | 0.820 | 0.854 | -0.76 | 0.448 | 0.507 |
| right postcentral | 0.05 | 0.258 | 0.313 | **0.32** | **3.08e-05** | **6.40e-05** | **2.30** | **0.022** | **0.032** |
| right posterior cingulate | -0.02 | 0.582 | 0.639 | **0.28** | **2.51e-04** | **4.71e-04** | **2.68** | **0.007** | **0.012** |
| right precentral | -0.02 | 0.583 | 0.639 | **0.22** | **0.004** | **0.006** | 1.94 | 0.052 | 0.073 |
| right precuneus | **0.18** | **0.015** | **0.023** | **0.29** | **1.32e-04** | **2.54e-04** | 1.01 | 0.310 | 0.369 |
| right rostral anterior cingulate | 1.70e-03 | 0.492 | 0.553 | **0.17** | **0.019** | **0.029** | 1.51 | 0.131 | 0.171 |
| right rostral middle frontal | **0.2** | **0.007** | **0.012** | 0.13 | 0.054 | 0.075 | -0.57 | 0.569 | 0.628 |
| right superior frontal | **0.24** | **0.001** | **0.002** | **0.22** | **0.003** | **0.005** | -0.19 | 0.850 | 0.878 |
| right superior parietal | **0.2** | **0.007** | **0.012** | **0.32** | **3.67e-05** | **7.59e-05** | 1.10 | 0.272 | 0.328 |
| right superior temporal | 0.12 | 0.074 | 0.101 | **0.47** | **3.62e-10** | **1.17e-09** | **3.62** | **2.98e-04** | **5.54e-04** |
| right supramarginal | 0.06 | 0.233 | 0.286 | **0.39** | **4.92e-07** | **1.24e-06** | **2.91** | **0.004** | **0.006** |
| right frontal pole | -0.11 | 0.913 | 0.934 | **0.22** | **0.003** | **0.005** | **2.96** | **0.003** | **0.005** |
| right temporal pole | 0.07 | 0.185 | 0.234 | **0.22** | **0.004** | **0.006** | 1.27 | 0.203 | 0.254 |
| right transverse temporal | -1.90e-03 | 0.509 | 0.570 | 0.12 | 0.064 | 0.089 | 1.16 | 0.247 | 0.301 |
| right insula | -0.03 | 0.663 | 0.711 | **0.19** | **0.008** | **0.013** | 1.93 | 0.054 | 0.075 |

Intraclass correlation coefficients (ICCs) were calculated between 3T scans and 64mT scans processed using the standard coronal T1w+T2w pipeline and the recon-all-clinical multi T1w pipeline. Differences in correlation strength were assessed using Steiger's Z test. Positive *Z* values indicate stronger correspondence with 3T scans for recon-all-clinical–processed regions relative to standard regions. Results that are significant at $p_{FDR} < 0.05$ after correction for multiple comparisons are shown in bold.

**Table S19. Regional Pearson correlation improvements: current vs. previous best-performing pipelines**

| | Previous | Current | Fisher's | | |
|---|---|---|---|---|---|
| **Region** | ***r*1** | ***r*2** | ***Z*** | ***p*** | **$p_{FDR}$** |
| left banks of superior temporal sulcus | -0.07 | 0.28 | 2.39 | 0.017 | 0.103 |
| left caudal anterior cingulate | 0.25 | 0.32 | 0.50 | 0.615 | 0.911 |
| left caudal middle frontal | 0.35 | 0.43 | 0.65 | 0.515 | 0.911 |
| left cuneus | 0.14 | 0.10 | -0.25 | 0.806 | 0.958 |
| left entorhinal | 0.01 | 0.22 | 1.49 | 0.137 | 0.390 |
| left frontal pole | -0.35 | 0.10 | **3.15** | **0.002** | **0.048** |
| left fusiform | 0.17 | 0.10 | -0.48 | 0.631 | 0.911 |
| left inferior parietal | 0.16 | 0.28 | 0.85 | 0.395 | 0.840 |
| left inferior temporal | 0.18 | 0.22 | 0.28 | 0.780 | 0.958 |
| left insula | -0.10 | 0.14 | 1.66 | 0.097 | 0.330 |
| left isthmus cingulate | 0.08 | 0.28 | 1.41 | 0.159 | 0.433 |
| left lateral occipital | 0.07 | -0.09 | -1.08 | 0.280 | 0.680 |
| left lateral orbitofrontal | 0.19 | 0.14 | -0.36 | 0.717 | 0.911 |
| left lingual | -0.06 | -0.06 | -0.02 | 0.986 | 0.986 |
| left medial orbitofrontal | 0.18 | 0.17 | -0.10 | 0.924 | 0.971 |
| left middle temporal | 0.30 | 0.31 | 0.08 | 0.938 | 0.971 |
| left paracentral | 0.04 | 0.13 | 0.62 | 0.538 | 0.911 |
| left parahippocampal | 0.36 | 0.58 | 1.95 | 0.051 | 0.215 |
| left pars opercularis | 0.33 | 0.43 | 0.77 | 0.443 | 0.860 |
| left pars orbitalis | -0.02 | 0.19 | 1.49 | 0.137 | 0.390 |
| left pars triangularis | 0.00 | 0.38 | 2.74 | 0.006 | 0.069 |
| left pericalcarine | -0.20 | -0.15 | 0.35 | 0.724 | 0.911 |
| left postcentral | 0.24 | 0.21 | -0.20 | 0.838 | 0.958 |
| left posterior cingulate | 0.24 | 0.23 | -0.10 | 0.922 | 0.971 |
| left precentral | 0.19 | 0.13 | -0.41 | 0.682 | 0.911 |
| left precuneus | 0.40 | 0.45 | 0.42 | 0.677 | 0.911 |
| left rostral anterior cingulate | -0.02 | 0.39 | 2.91 | 0.004 | 0.062 |
| left rostral middle frontal | 0.09 | 0.28 | 1.38 | 0.167 | 0.436 |
| left superior frontal | 0.36 | 0.43 | 0.56 | 0.575 | 0.911 |
| left superior parietal | 0.11 | 0.41 | 2.23 | 0.025 | 0.144 |
| left superior temporal | 0.31 | 0.42 | 0.87 | 0.382 | 0.838 |
| left supramarginal | 0.41 | 0.4 | -0.08 | 0.939 | 0.971 |
| left temporal pole | 0.14 | 0.17 | 0.22 | 0.828 | 0.958 |
| left transverse temporal | 0.27 | 0.17 | -0.69 | 0.489 | 0.899 |
| right banks of superior temporal sulcus | 0.10 | 0.21 | 0.75 | 0.451 | 0.860 |
| right caudal anterior cingulate | 0.01 | 0.47 | **3.40** | **6.84e-04** | **0.046** |
| right caudal middle frontal | 0.29 | 0.22 | -0.52 | 0.603 | 0.911 |
| right cuneus | 0.14 | 0.07 | -0.48 | 0.630 | 0.911 |
| right entorhinal | -0.05 | 0.22 | 1.86 | 0.063 | 0.242 |
| right frontal pole | -0.14 | 0.17 | 2.12 | 0.034 | 0.166 |

| | | | | | |
|---|---|---|---|---|---|
| right fusiform | -0.35 | 0.05 | 2.78 | 0.005 | 0.069 |
| right inferior parietal | 0.06 | 0.28 | 1.56 | 0.118 | 0.363 |
| right inferior temporal | 0.00 | 0.24 | 1.67 | 0.096 | 0.330 |
| right insula | 0.17 | 0.20 | 0.26 | 0.792 | 0.958 |
| right isthmus cingulate | 0.06 | 0.41 | 2.55 | 0.011 | 0.086 |
| right lateral occipital | 0.06 | 0.09 | 0.20 | 0.845 | 0.958 |
| right lateral orbitofrontal | 0.09 | 0.20 | 0.78 | 0.436 | 0.860 |
| right lingual | 0.06 | -0.08 | -0.92 | 0.355 | 0.820 |
| right medial orbitofrontal | -0.05 | 0.30 | 2.47 | 0.014 | 0.093 |
| right middle temporal | 0.17 | 0.45 | 2.13 | 0.034 | 0.166 |
| right paracentral | 0.09 | 0.15 | 0.43 | 0.670 | 0.911 |
| right parahippocampal | 0.40 | 0.40 | 0.03 | 0.979 | 0.986 |
| right pars opercularis | 0.20 | 0.52 | 2.53 | 0.011 | 0.086 |
| right pars orbitalis | 0.28 | 0.39 | 0.91 | 0.362 | 0.820 |
| right pars triangularis | 0.25 | 0.24 | -0.07 | 0.942 | 0.971 |
| right pericalcarine | -0.01 | -0.07 | -0.40 | 0.692 | 0.911 |
| right postcentral | 0.27 | 0.31 | 0.36 | 0.716 | 0.911 |
| right posterior cingulate | 0.24 | 0.32 | 0.57 | 0.566 | 0.911 |
| right precentral | -0.06 | 0.12 | 1.23 | 0.219 | 0.551 |
| right precuneus | 0.20 | 0.45 | 1.85 | 0.064 | 0.242 |
| right rostral anterior cingulate | -0.13 | 0.31 | **3.07** | **0.002** | **0.048** |
| right rostral middle frontal | 0.03 | 0.25 | 1.57 | 0.116 | 0.363 |
| right superior frontal | 0.11 | 0.39 | 2.07 | 0.038 | 0.173 |
| right superior parietal | 0.08 | 0.43 | 2.57 | 0.010 | 0.086 |
| right superior temporal | 0.43 | 0.51 | 0.75 | 0.455 | 0.860 |
| right supramarginal | 0.39 | 0.31 | -0.60 | 0.546 | 0.911 |
| right temporal pole | 0.22 | 0.20 | -0.17 | 0.863 | 0.962 |
| right transverse temporal | 0.06 | 0.13 | 0.42 | 0.676 | 0.911 |

Improvements in regional Pearson correlation coefficients were assessed by comparing the current study's best-performing pipeline (recon-all-clinical with multi T2w; $r_2$) with that of the previous study (SynthSR v1 with axial T1w and T2w; $r_1$) (Cooper et al., 2024) using Fisher's Z test. Positive $Z$ values indicate stronger correspondence with 3T MRI for the current study relative to the previous study. Results that remained significant after multiple comparison correction ($p_{FDR} < 0.05$) are shown in bold.

**Table S20. Regional intraclass correlation improvements: current vs. previous best-performing pipelines**

| Region | Previous ICC1 | Current ICC2 | Fisher's Z | p | $p_{FDR}$ |
|---|---|---|---|---|---|
| left banks of superior temporal sulcus | -0.06 | 0.28 | 2.36 | 0.018 | 0.113 |
| left caudal anterior cingulate | 0.23 | 0.32 | 0.66 | 0.509 | 0.887 |
| left caudal middle frontal | 0.35 | 0.41 | 0.48 | 0.634 | 0.909 |
| left cuneus | 0.14 | 0.10 | -0.28 | 0.783 | 0.934 |
| left entorhinal | 0.00 | 0.20 | 1.35 | 0.178 | 0.476 |
| left frontal pole | -0.33 | 0.10 | 3.01 | 0.003 | 0.058 |
| left fusiform | 0.16 | 0.10 | -0.41 | 0.679 | 0.909 |
| left inferior parietal | 0.16 | 0.28 | 0.86 | 0.392 | 0.859 |
| left inferior temporal | 0.16 | 0.22 | 0.42 | 0.673 | 0.909 |
| left insula | -0.09 | 0.14 | 1.57 | 0.117 | 0.397 |
| left isthmus cingulate | 0.08 | 0.27 | 1.33 | 0.182 | 0.476 |
| left lateral occipital | 0.06 | -0.09 | -1.02 | 0.308 | 0.775 |
| left lateral orbitofrontal | 0.19 | 0.14 | -0.35 | 0.727 | 0.916 |
| left lingual | -0.06 | -0.06 | 0.00 | 1.000 | 1.000 |
| left medial orbitofrontal | 0.17 | 0.17 | 0.00 | 1.000 | 1.000 |
| left middle temporal | 0.29 | 0.30 | 0.07 | 0.941 | 0.987 |
| left paracentral | 0.04 | 0.13 | 0.62 | 0.538 | 0.909 |
| left parahippocampal | 0.35 | 0.56 | 1.81 | 0.070 | 0.279 |
| left pars opercularis | -0.02 | 0.19 | 1.44 | 0.150 | 0.443 |
| left pars orbitalis | 0.00 | 0.37 | 2.64 | 0.008 | 0.096 |
| left pars triangularis | 0.33 | 0.42 | 0.71 | 0.477 | 0.887 |
| left pericalcarine | -0.20 | -0.15 | 0.35 | 0.726 | 0.916 |
| left postcentral | 0.22 | 0.20 | -0.14 | 0.887 | 0.958 |
| left posterior cingulate | 0.23 | 0.22 | -0.07 | 0.943 | 0.987 |
| left precentral | 0.19 | 0.11 | -0.56 | 0.579 | 0.909 |
| left precuneus | 0.40 | 0.44 | 0.33 | 0.742 | 0.917 |
| left rostral anterior cingulate | -0.02 | 0.39 | 2.93 | 0.003 | 0.058 |
| left rostral middle frontal | 0.09 | 0.28 | 1.34 | 0.180 | 0.476 |
| left superior frontal | 0.36 | 0.43 | 0.56 | 0.573 | 0.909 |
| left superior parietal | 0.11 | 0.40 | 2.12 | 0.034 | 0.190 |
| left superior temporal | 0.31 | 0.40 | 0.70 | 0.484 | 0.887 |
| left supramarginal | 0.41 | 0.38 | -0.24 | 0.809 | 0.949 |
| left temporal pole | 0.13 | 0.17 | 0.28 | 0.781 | 0.934 |
| left transverse temporal | 0.27 | 0.16 | -0.78 | 0.433 | 0.865 |
| right banks of superior temporal sulcus | 0.10 | 0.21 | 0.77 | 0.444 | 0.865 |
| right caudal anterior cingulate | 0.01 | 0.47 | **3.40** | **6.72e-04** | **0.046** |
| right caudal middle frontal | 0.29 | 0.21 | -0.58 | 0.562 | 0.909 |
| right cuneus | 0.14 | 0.07 | -0.48 | 0.631 | 0.909 |

| | | | | | |
|---|---|---|---|---|---|
| right entorhinal | -0.05 | 0.21 | 1.79 | 0.074 | 0.280 |
| right frontal pole | -0.12 | 0.17 | 1.98 | 0.047 | 0.215 |
| right fusiform | -0.32 | 0.05 | 2.59 | 0.010 | 0.096 |
| right inferior parietal | 0.06 | 0.27 | 1.47 | 0.141 | 0.437 |
| right inferior temporal | 0.00 | 0.23 | 1.61 | 0.108 | 0.387 |
| right insula | 0.15 | 0.20 | 0.35 | 0.726 | 0.916 |
| right isthmus cingulate | 0.06 | 0.41 | 2.55 | 0.011 | 0.096 |
| right lateral occipital | 0.06 | 0.09 | 0.20 | 0.838 | 0.950 |
| right lateral orbitofrontal | 0.09 | 0.20 | 0.76 | 0.445 | 0.865 |
| right lingual | 0.06 | -0.08 | -0.95 | 0.341 | 0.801 |
| right medial orbitofrontal | -0.05 | 0.30 | 2.44 | 0.015 | 0.100 |
| right middle temporal | 0.17 | 0.44 | 2.04 | 0.041 | 0.201 |
| right paracentral | 0.09 | 0.15 | 0.41 | 0.680 | 0.909 |
| right parahippocampal | 0.38 | 0.40 | 0.16 | 0.873 | 0.958 |
| right pars opercularis | 0.27 | 0.39 | 0.92 | 0.360 | 0.816 |
| right pars orbitalis | 0.24 | 0.24 | 0.00 | 1.000 | 1.000 |
| right pars triangularis | 0.20 | 0.52 | 2.53 | 0.011 | 0.096 |
| right pericalcarine | -0.01 | -0.07 | -0.43 | 0.671 | 0.909 |
| right postcentral | 0.24 | 0.27 | 0.22 | 0.828 | 0.950 |
| right posterior cingulate | 0.23 | 0.32 | 0.66 | 0.509 | 0.887 |
| right precentral | -0.05 | 0.09 | 0.95 | 0.341 | 0.801 |
| right precuneus | 0.20 | 0.44 | 1.83 | 0.068 | 0.279 |
| right rostral anterior cingulate | -0.12 | 0.31 | 2.99 | 0.003 | 0.058 |
| right rostral middle frontal | 0.03 | 0.25 | 1.53 | 0.126 | 0.409 |
| right superior frontal | 0.11 | 0.39 | 2.04 | 0.041 | 0.201 |
| right superior parietal | 0.08 | 0.42 | 2.49 | 0.013 | 0.096 |
| right superior temporal | 0.43 | 0.49 | 0.52 | 0.605 | 0.909 |
| right supramarginal | 0.39 | 0.29 | -0.77 | 0.442 | 0.865 |
| right temporal pole | 0.22 | 0.20 | -0.14 | 0.887 | 0.958 |
| right transverse temporal | 0.06 | 0.12 | 0.41 | 0.681 | 0.909 |

Improvements in regional intraclass correlation coefficients (ICCs) were assessed by comparing the current study's best-performing pipeline (recon-all-clinical with multi T2w; $ICC_2$) with that of the previous study (SynthSR v1 with axial T1w and T2w; $ICC_1$) (Cooper et al., 2024) using Fisher's Z test. Positive *Z* values indicate stronger correspondence with 3T MRI for the current study relative to the previous study. Results that remained significant after multiple comparison correction ($p_{FDR} < 0.05$) are shown in bold.
